# Genetic, transcriptome, proteomic and epidemiological evidence for blood brain barrier disruption and polymicrobial brain invasion as determinant factors in Alzheimer’s disease

**DOI:** 10.1101/080333

**Authors:** C.J. Carter

## Abstract

Multiple pathogens have been detected in Alzheimer’s disease (AD) brains. A bioinformatics approach was used to assess relationships between pathogens and AD genes (GWAS), the AD hippocampal transcriptome and plaque or tangle proteins. Host/pathogen interactomes (*C.albicans*, *C.Neoformans*, Bornavirus, *B.Burgdorferri*, cytomegalovirus, Ebola virus, HSV-1, HERV-W, HIV-1, Epstein-Barr, hepatitis C, influenza, *C.Pneumoniae*, *P.Gingivalis*, *H.Pylori*, *T.Gondii*, *T.Cruzi*) significantly overlap with misregulated AD hippocampal genes, with plaque and tangle proteins and, except Bornavirus, Ebola and HERV-W, with AD genes. Upregulated AD hippocampal genes match those upregulated by multiple bacteria, viruses, fungi or protozoa in immunocompetent blood cells. AD genes are enriched in bone marrow and immune locations and in GWAS datasets reflecting pathogen diversity, suggesting selection for pathogen resistance. The age of AD patients implies resistance to infections afflicting the younger. APOE4 protects against malaria and hepatitis C, and immune/inflammatory gain of function applies to APOE4, CR1, TREM2 and presenilin variants. 30/78 AD genes are expressed in the blood brain barrier (BBB), which is disrupted by AD risk factors (ageing, alcohol, aluminium, concussion, cerebral hypoperfusion, diabetes, homocysteine, hypercholesterolaemia, hypertension, obesity, pesticides, pollution, physical inactivity, sleep disruption and smoking). The BBB and AD benefit from statins, NSAIDs, oestrogen, melatonin and the Mediterranean diet. Polymicrobial involvement is supported by the upregulation of pathogen sensors/defenders (bacterial, fungal, viral) in the AD brain, blood or CSF. Cerebral pathogen invasion permitted by BBB inadequacy, activating a hyper-efficient immune/inflammatory system, beta-amyloid and other antimicrobial defence may be responsible for AD which may respond to antibiotic, antifungal or antiviral therapy.

## Introduction

Multiple pathogens have been implicated in Alzheimer’s disease (AD) either via detection in the AD brain, or in epidemiological studies relating to serum antibodies. Pathological burden (cytomegalovirus, Herpes simplex (HSV-1), *Borrelia burgdorferi*, *Chlamydia pneumoniae* and *Helicobacter pylori*) rather than any individual pathogen may also be associated with AD [1]. Many pathogens are able to increase beta-amyloid deposition and tau phosphorylation in animal models, *in vitro* or *in vivo* and beta-amyloid itself is an antimicrobial peptide active against bacteria and fungi [2,3]and the influenza[4] and herpes simplex viruses [5,6]. These effects are summarised in Table 1 for a number of pathogens and for beta-amyloid.

**Table 1:**
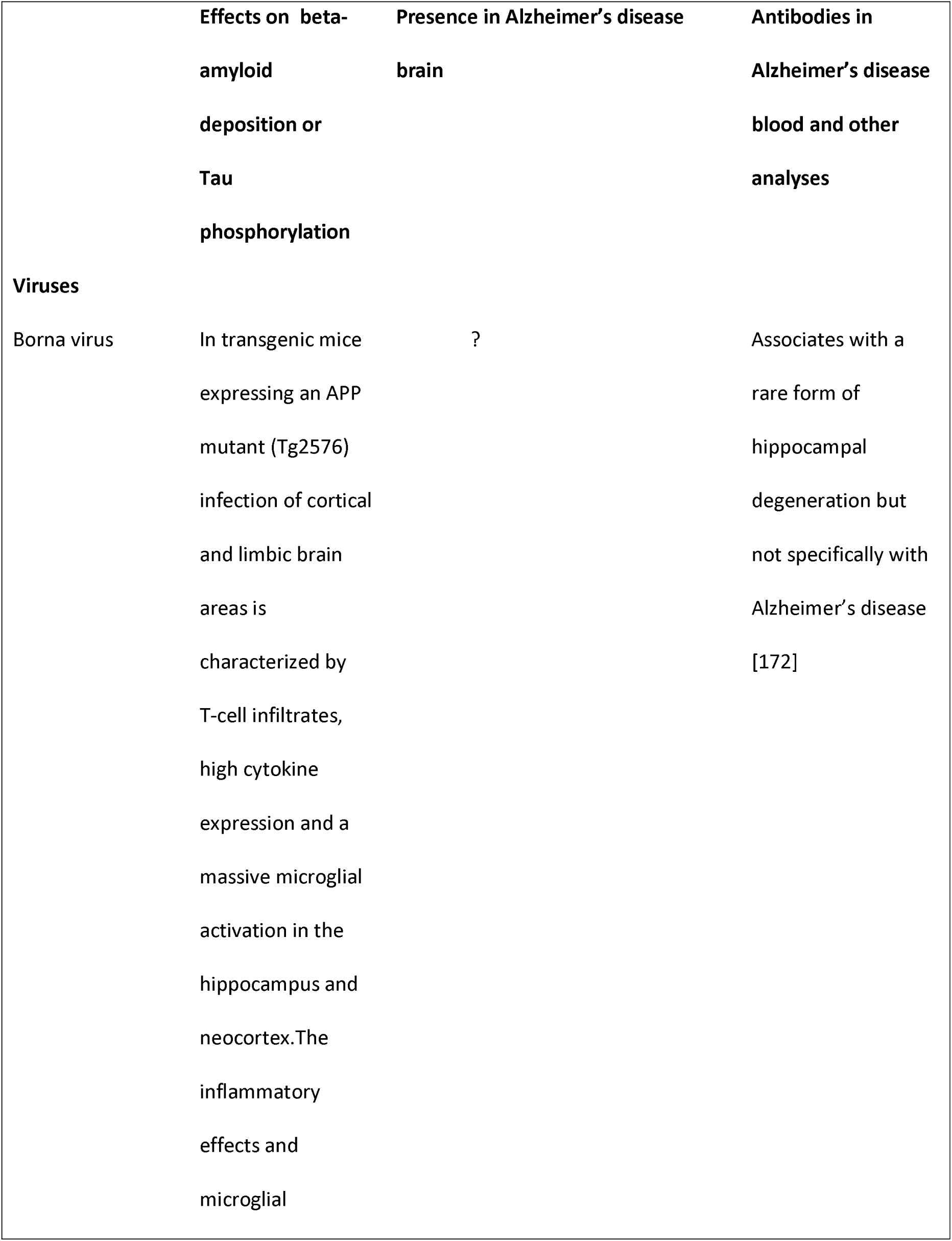
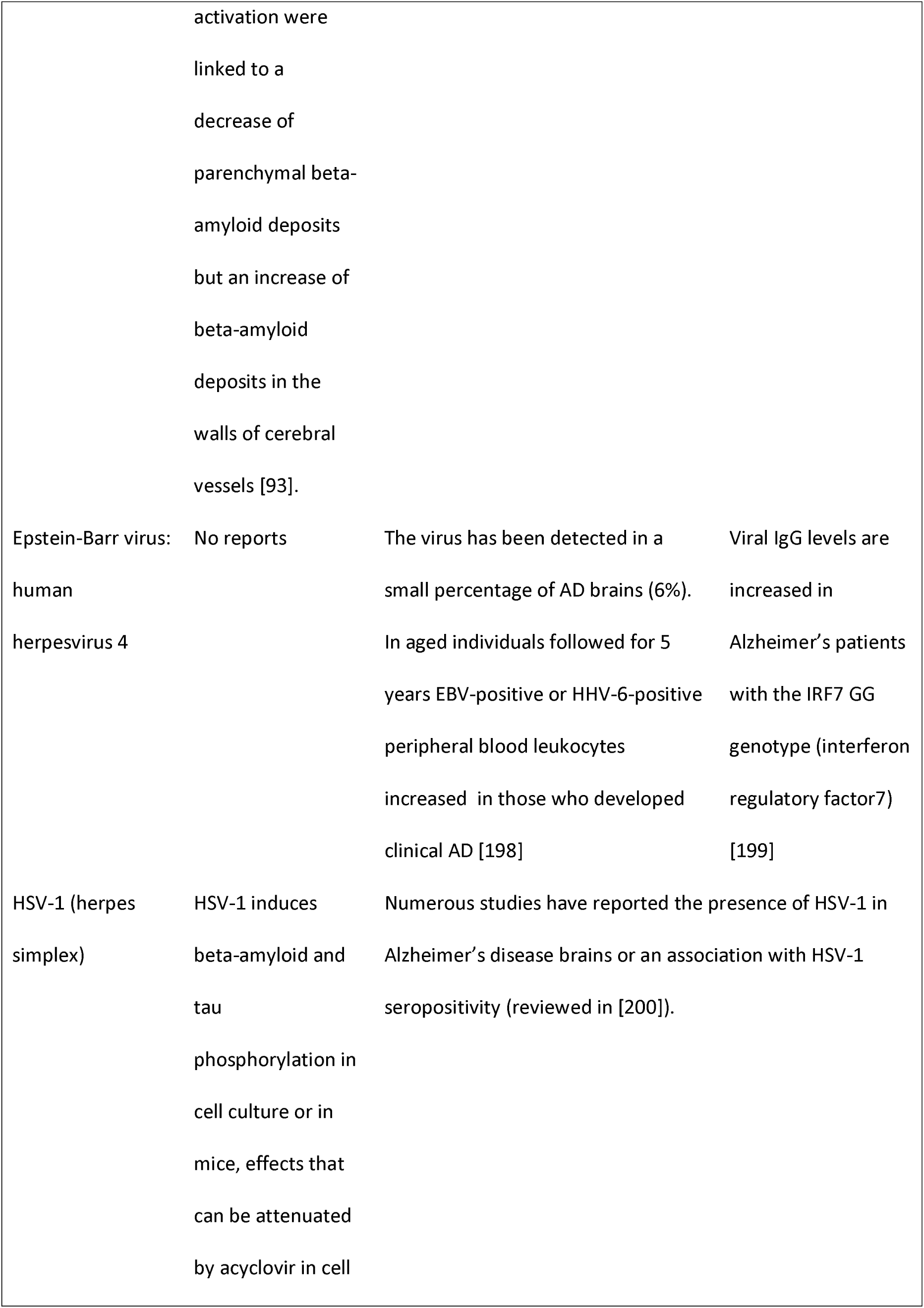
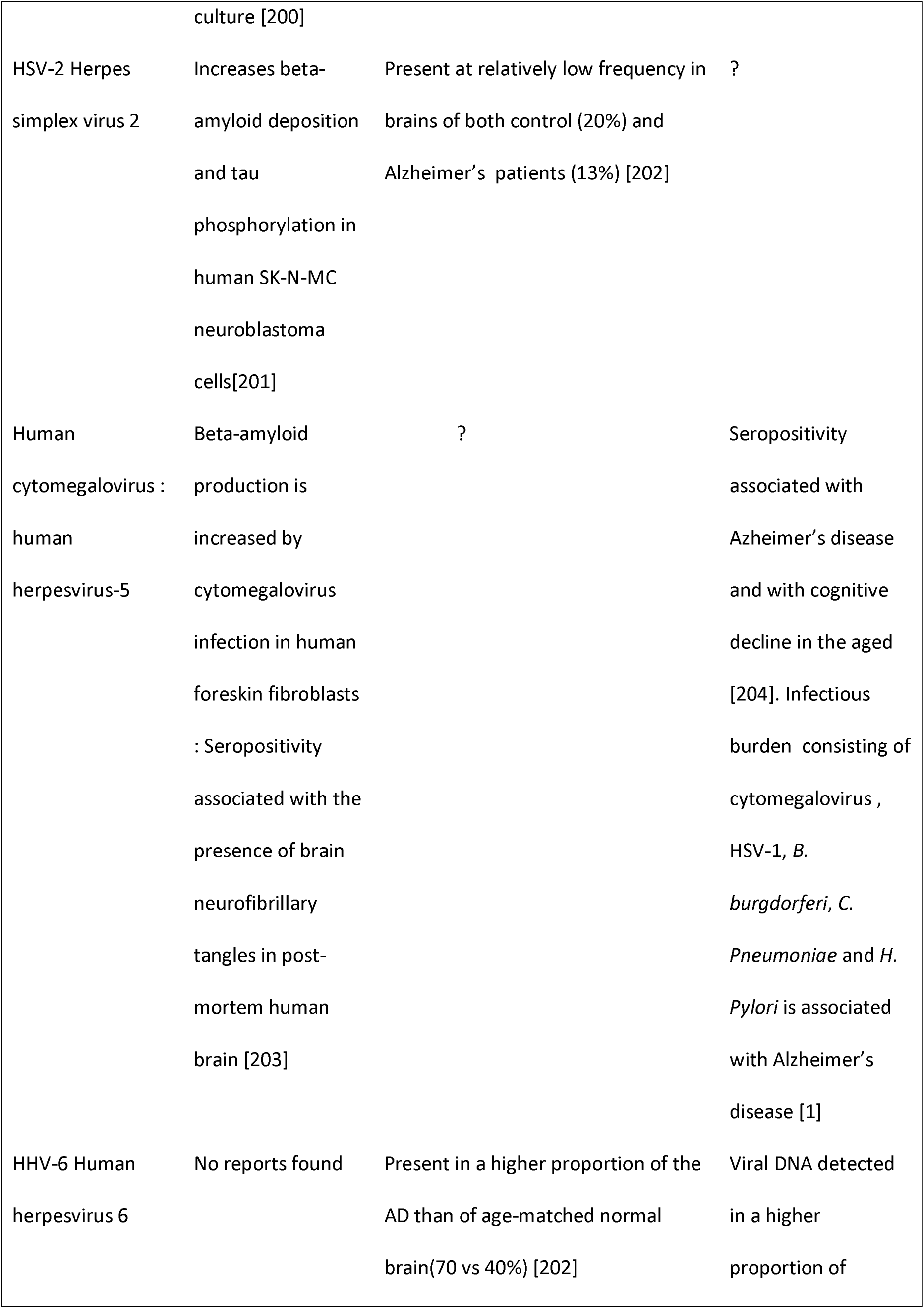
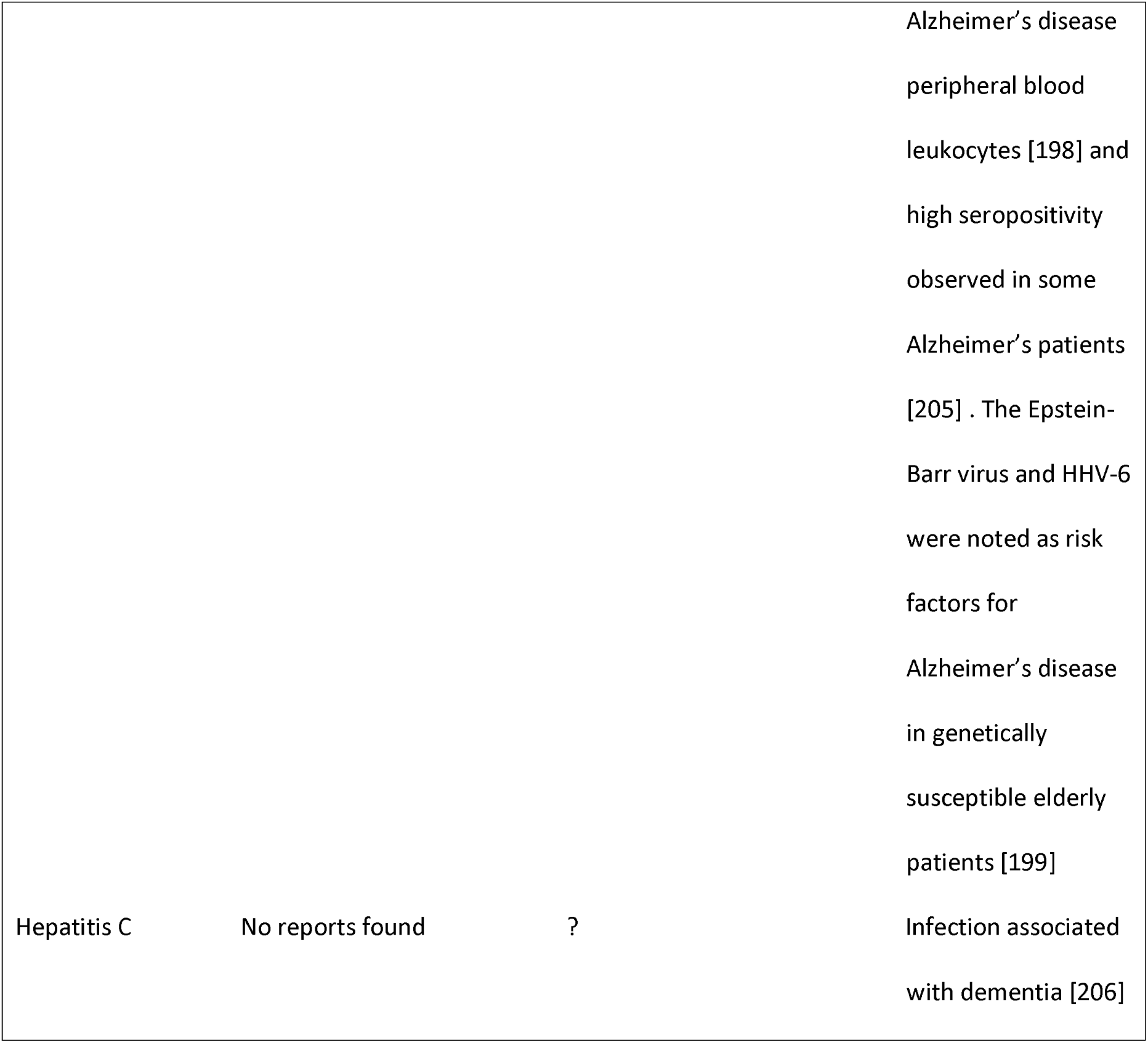
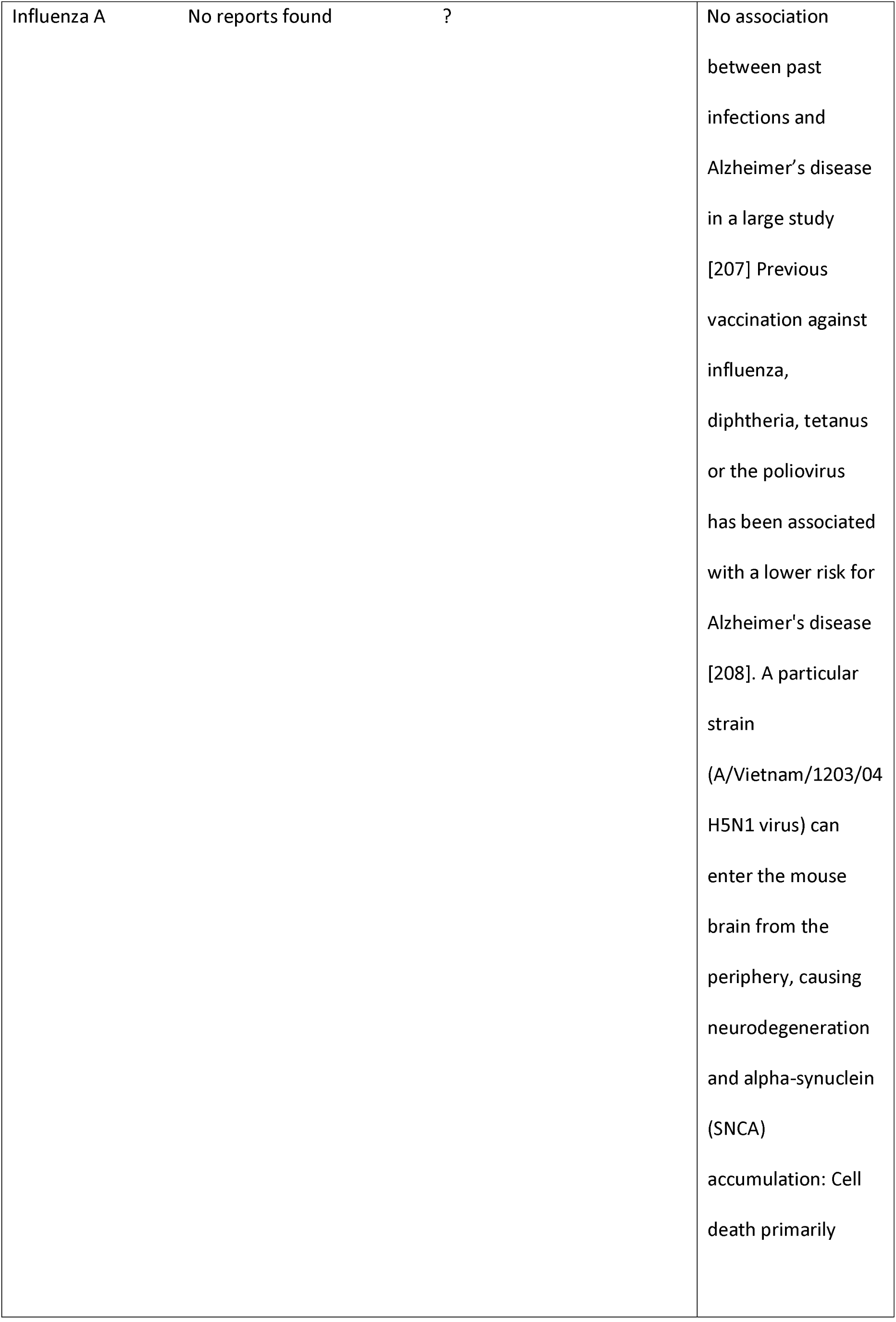
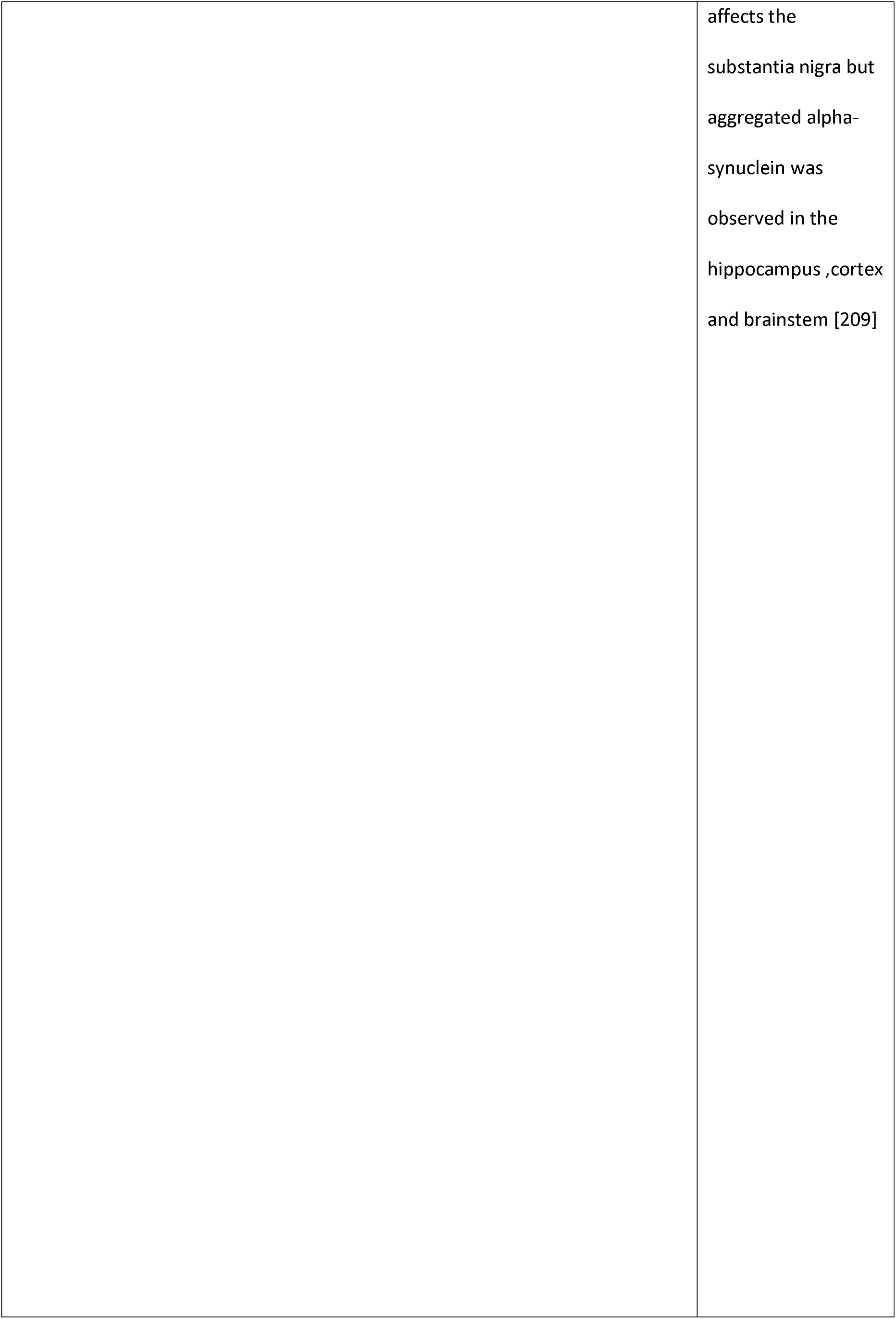
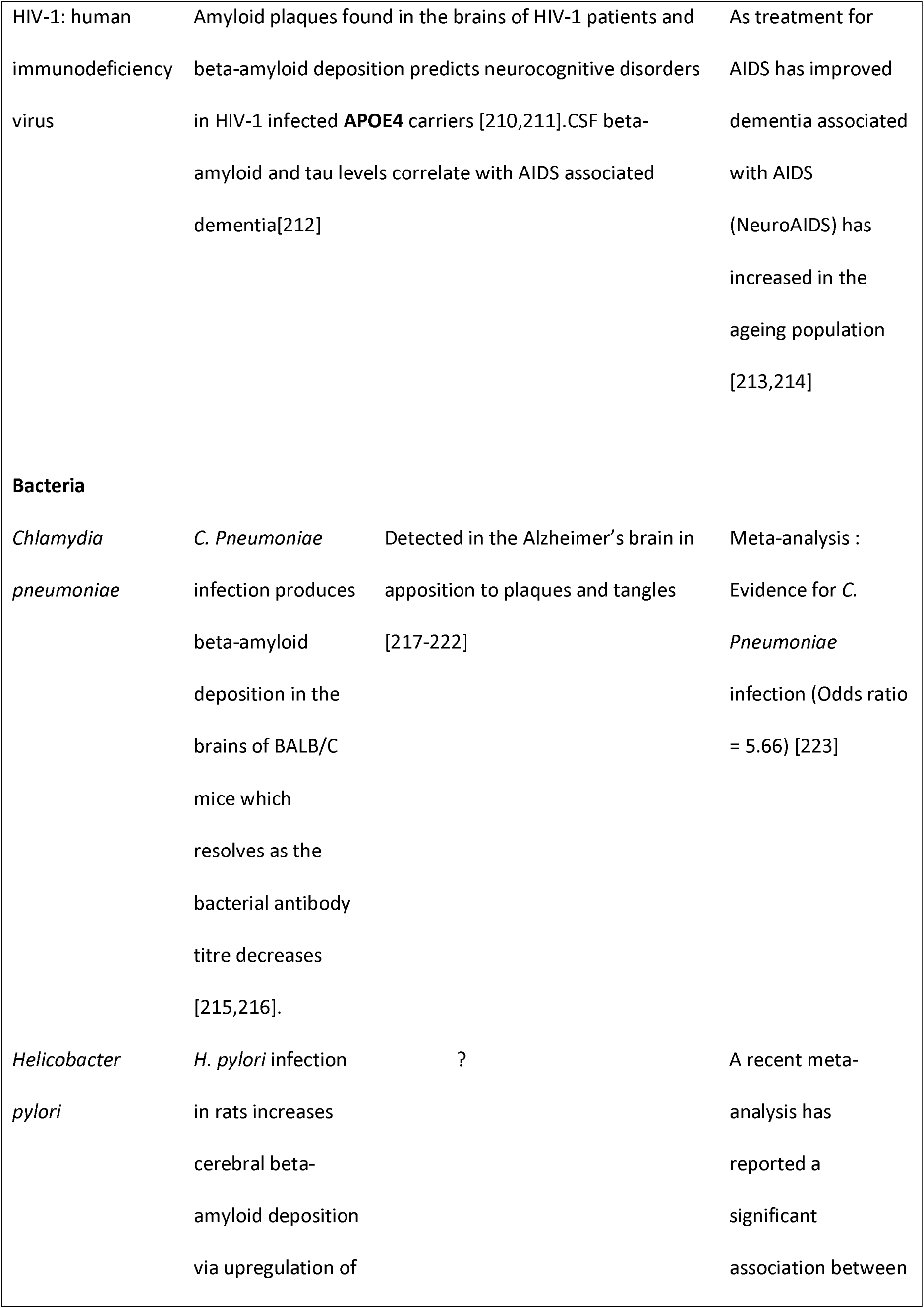
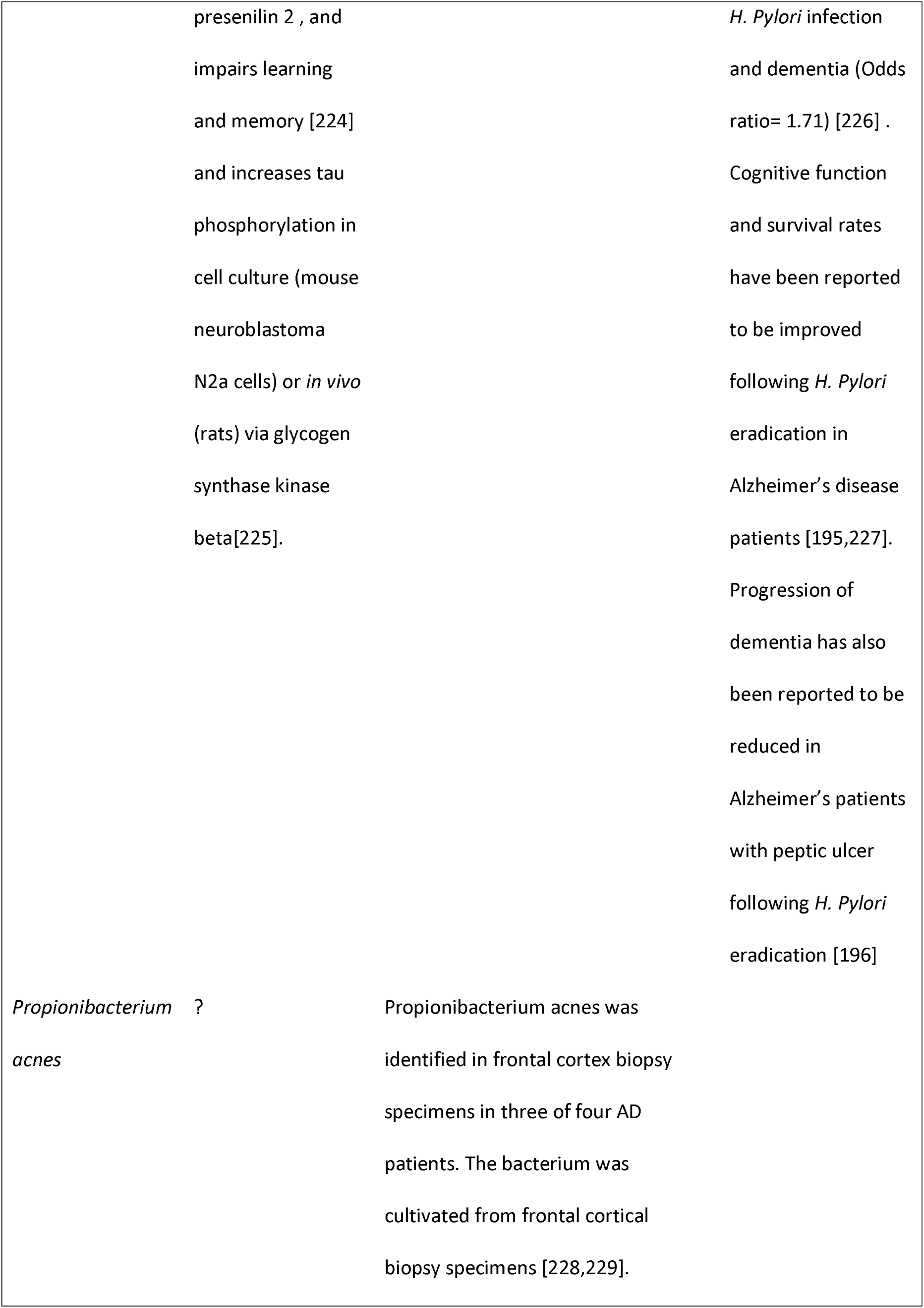
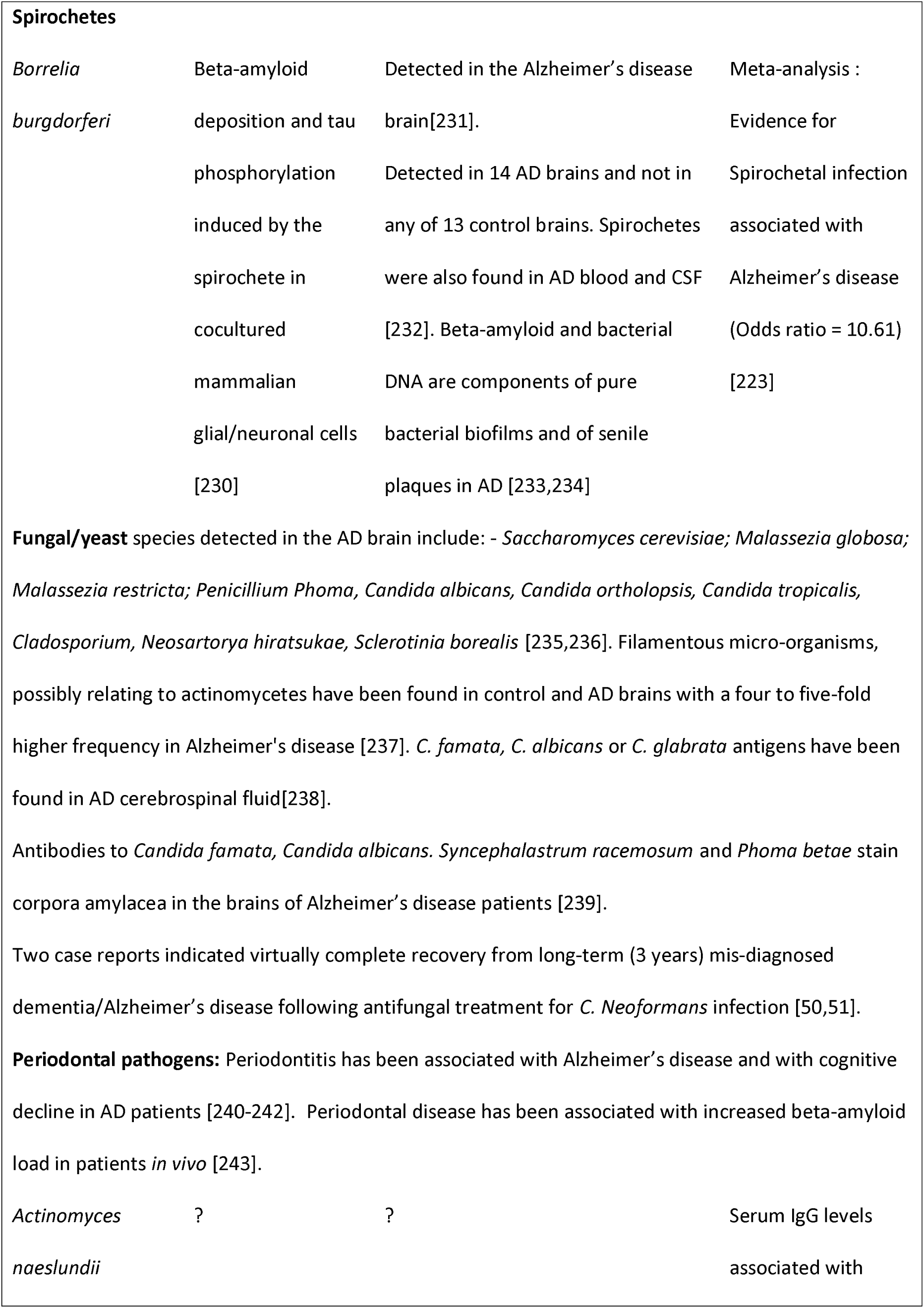
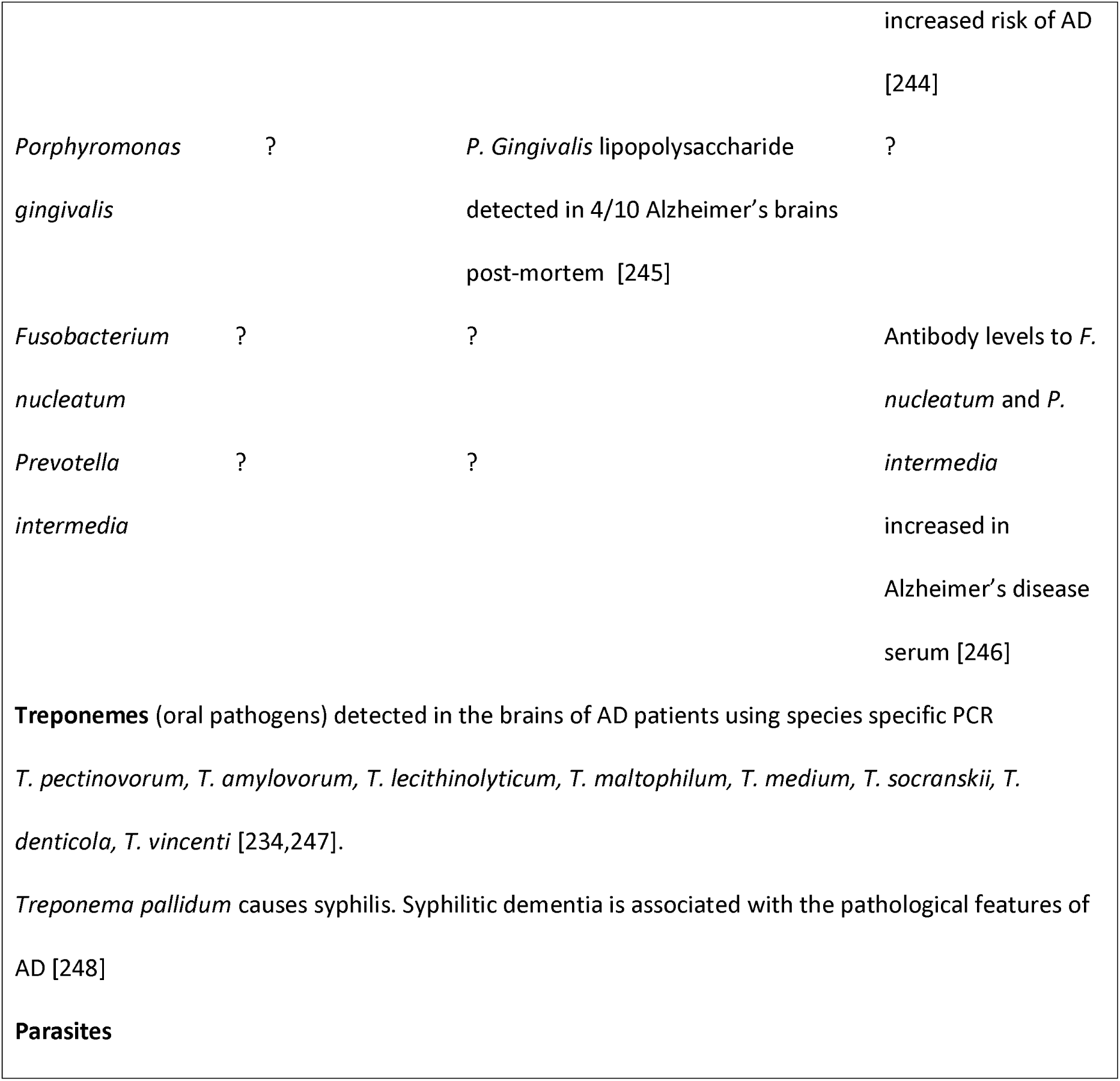
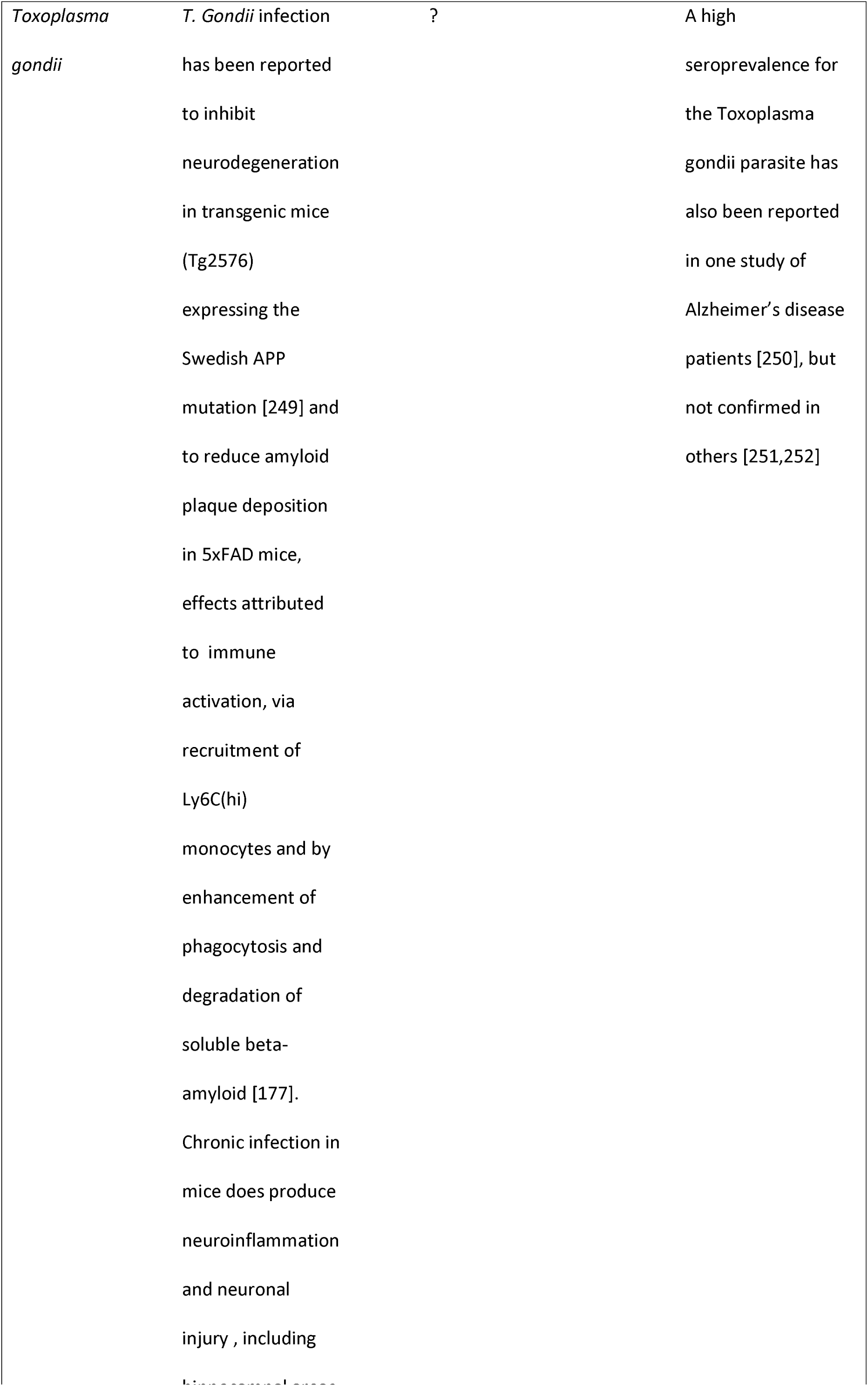
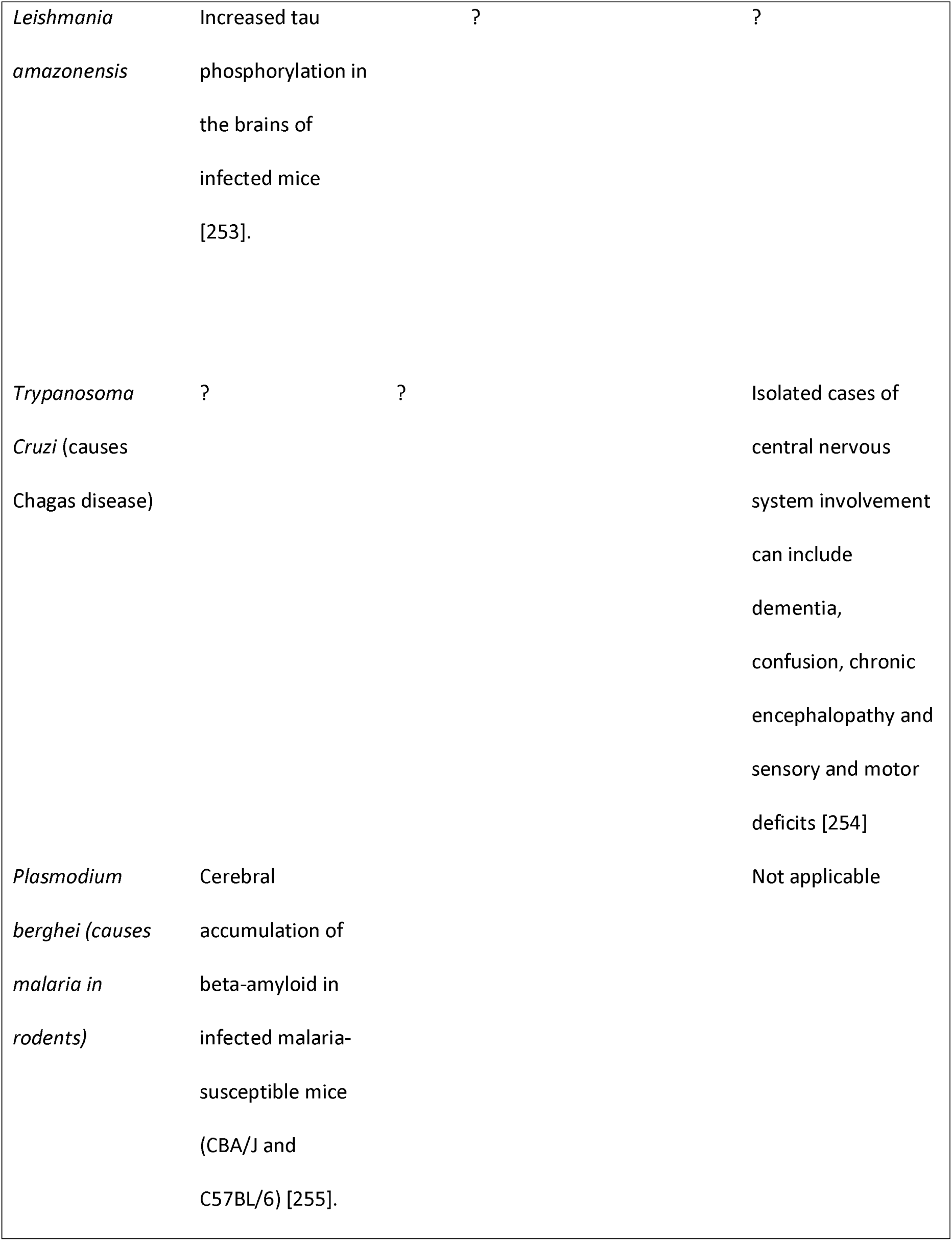
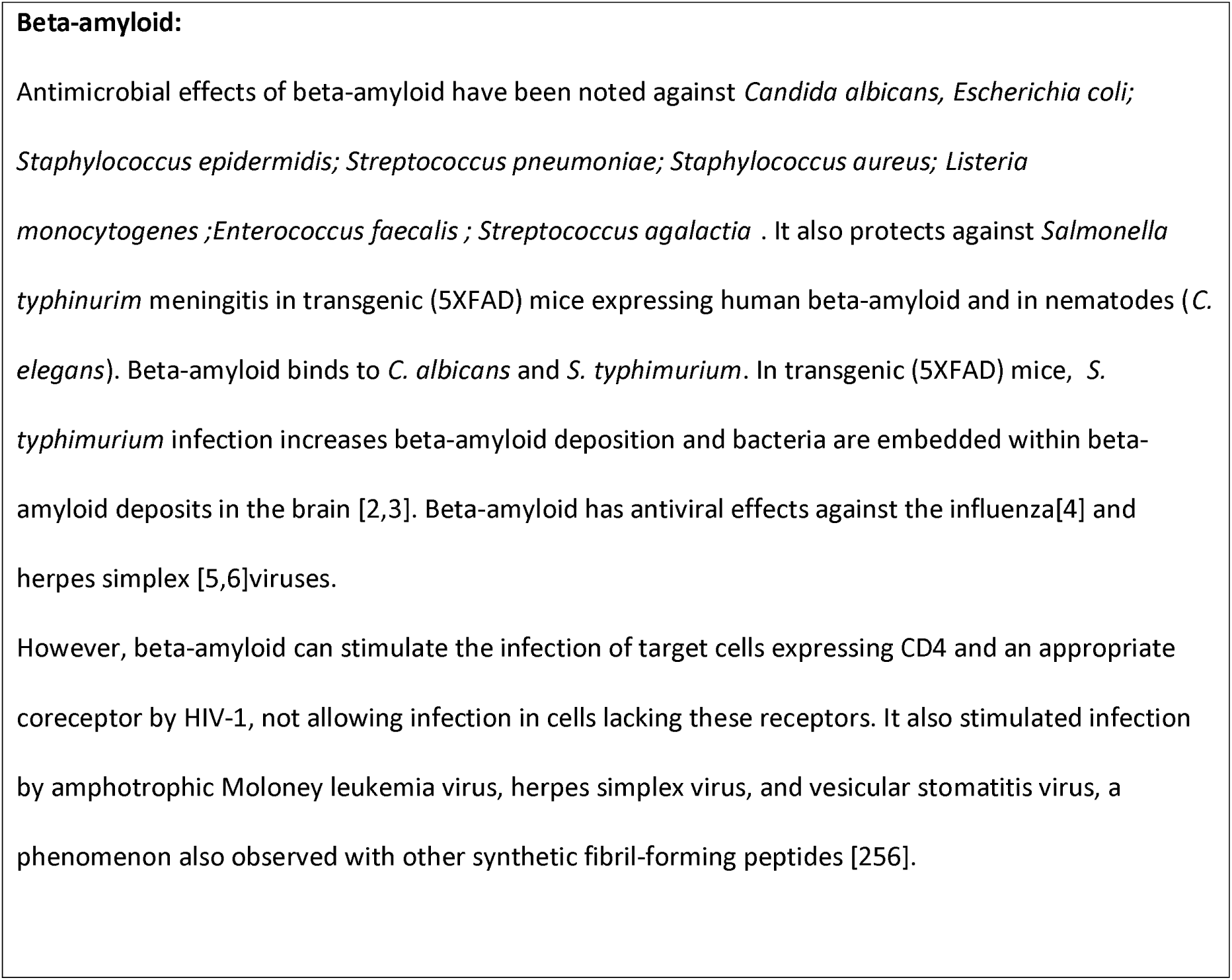
The effects of diverse pathogens on beta-amyloid deposition, tau phosphorylation and their relationships with Alzheimer’s disease.

Previous studies have shown that the life cycles of several pathogens implicated in AD relate to AD susceptibility genes [7]. The proteins found in AD plaques and tangles are also enriched in those used by HSV-1 during its life cycle [8] and the HSV-1 or *Toxoplasma Gondii* host interactomes are also enriched in AD susceptibility genes [9,10]

Similar studies have noted significant overlaps between the Epstein-Barr viral/host interactome and diseases in which the virus is implicated, including B cell lymphoma [11]or multiple sclerosis [12]. The interactomes of oncogenic viruses also relate to cancer genes [13] suggesting important gene/environment interactions that may condition disease susceptilbility.

In this study, the host pathogen interactomes of 17 fungal, bacterial, viral and parasite pathogens were analysed in relation to 78 AD genes derived from genome-wide association studies (GWAS). The anatomical location of these genes was also queried against proteomic /genomic datasets from multiple tissues.

The host genes of the pathogen interactomes were also compared with the combined up and downregulated genes from a study of the AD hippocampus, post-mortem [14] and to the proteins found in plaques or neurofibrillary tangles. The upregulated genes from this AD hippocampal study were also compared with upregulated genes from numerous infection microarray datasets (viral, bacterial, fungal and protozoan) housed at the Molecular signatures database [15] or the Gene Expression Omnibus [16].

Pathogens have shaped human evolution, as the survivors of dangerous infections are endowed, via natural selection, with genes conveying resistance. The AD genes were also compared against a series of genome-wide association datasets related to general pathogen or protozoan diversity, viral diversity and the immune response to parasitic worms, across multiple human populations in different geographical locations. Such genes are likely to have been selected for pathogen resistance. [17–20].

The results show that host genes related to pathogens are enriched in all these AD parameters and that many AD susceptibility genes also relate to pathogens, but more likely to pathogen resistance than susceptibility. The anatomical data point to an immune function of many AD genes, while others are localised in the blood-brain barrier, which is disrupted by other environmental risk factors associated with AD.

## Methods

The host/pathogen interactomes of two fungal species (*Candida albicans*, *Cryptococcus Neoformans*), the Borna virus, human cytomegalovirus, Ebola virus, Herpes simplex (HSV-1), human endogenous retroviruses HERV-W, the human immunodeficiency virus (HIV-1) (the latter from the HIV-1, human interaction database [21]

http://www.ncbi.nlm.nih.gov/genome/viruses/retroviruses/hiv-1/interactions, Epstein-Barr, hepatitis C and influenza A viruses, 3 bacterial species (*Chlamydia Pneumoniae*, *Porphyromonas Gingivalis*, *Helicobacter Pylori*) and 2 protozoans (*Toxoplasma Gondii* and *Trypanosoma Cruzi*) were obtained by literature survey and from extant databases. These referenced interactomes can be accessed at http://www.polygenicpathways.co.uk/HPI.htm.

Genes misregulated in the AD hippocampus are those reported from a post-mortem microarray study [14]. Up-and downregulated genes (N=2879) were combined for comparison with the pathogen interactomes. These interactomes contain multiple types of interaction (protein/protein, viral microRNA, and effects on transcription etc.) and it is not possible to compare like with like for this aspect.

The upregulated genes (N= 1690) from this AD hippocampal study contain the pathways relevant to pathogens and immune activation (inflammation, complement activation and the defence response) [14] and these were chosen for comparison with upregulated genes from infection datasets at the Molecular signatures database (MSigDB) http://software.broadinstitute.org/gsea/msigdb/index.jsp. MSigDB contains several thousand microarray gene sets which can be compared against the AD input [15]. Infection related datasets, and those related to Toll-like receptor ligands, were identified using search terms (e.g. infection, virus, bacteria, TLR1, lipopolysaccharide, etc.). Microarray viral infection datasets (upregulated gene sets) from the gene expression omnibus (GEO) [22] were also downloaded from the Harmonizome database http://amp.pharm.mssm.edu/Harmonizome/ from the Ma'ayan laboratory of computational systems. [23]. For the searched gene sets, most of the data outputs were restricted at source (by MSigDb or GEO) to the top upregulated genes (usually ~ 200-300).

The proteins found in plaques or neurofibrillary tangles are from two proteomics studies yielding 488 proteins in plaques [24] and 90 in tangles [25].

Seventy eight genes associated with Alzheimer’s disease (Reported genes) were obtained from the NHGRI-EBI Catalog of published genome-wide association studies (GWAS) [26], Available at: www.ebi.ac.uk/gwas. Accessed January, 2016, version 1.0 using studies labelled as “Alzheimer’s disease” or “Alzheimer’s disease late-onset”. These genes and their relationships with pathogens or the immune system are catalogued in Supplementary Table 1. These genes are highlighted in bold throughout the text.

Genes related to general pathogen diversity, protozoan and viral diversity and to the immune response to parasitic worms are from a series of papers concerning evolutionary selection pressure relevant to pathogen resistance [17–20].

The tissue and cellular distribution of the 78 AD genes were analysed using the functional enrichment analysis tool (FUNRICH) [27]. http://funrich.org/index.html. This tool derives proteomic and genomic distribution data from >1.5 million annotations. It provides the total number of genes in datasets from each region sampled and returns the significance of any enrichment for members of the uploaded AD genes, using the hypergeometric probability test, with p values corrected using the the Storey and Tibshirani method (Q values) [27]. AD gene enrichment was also analysed in a published blood brain barrier proteome dataset of mouse cerebral arteries (6620 proteins) [28].

The presence of the AD genes in exosomes, a means of transit through cells allowing intercellular communication[29,30], was assessed using ExoCarta (http://www.exocarta.org) a manually curated database of exosomal proteins, RNA and lipids [31].The exosomal pathway is hijacked by several viruses, contributing to intercellular spread and immune evasion [32,33].

Assuming a human genome of 26846 coding genes and an interactome or other gene set of N genes one would expect N/26846 to exist in the comparator dataset. For example, when comparing 2879 misregulated AD hippocampal genes against any pathogen interactome one would expect 2879/26846 (10.7%) to figure in the pathogen interactome. This calculation was used to define expected values and the enrichment values (observed/expected) in relation to other datasets. Significance of the enrichment was calculated using the hypergeometric probability test. The resultant p values from each analysis series were corrected for false discovery (FDR) [34]. Nominally significant FDR corrected values are considered at P <0.05 and a corrected Bonferroni p value threshold is illustrated on each set of graphs. (Bonferroni P = 0.05/N, where N is the maximum number of possible comparisons for each situation (e.g. 78 AD genes or 1690 upregulated genes in the AD hippocampus).

## Results

### The anatomical location of the AD genes (Fig 1)

**Fig 1:**
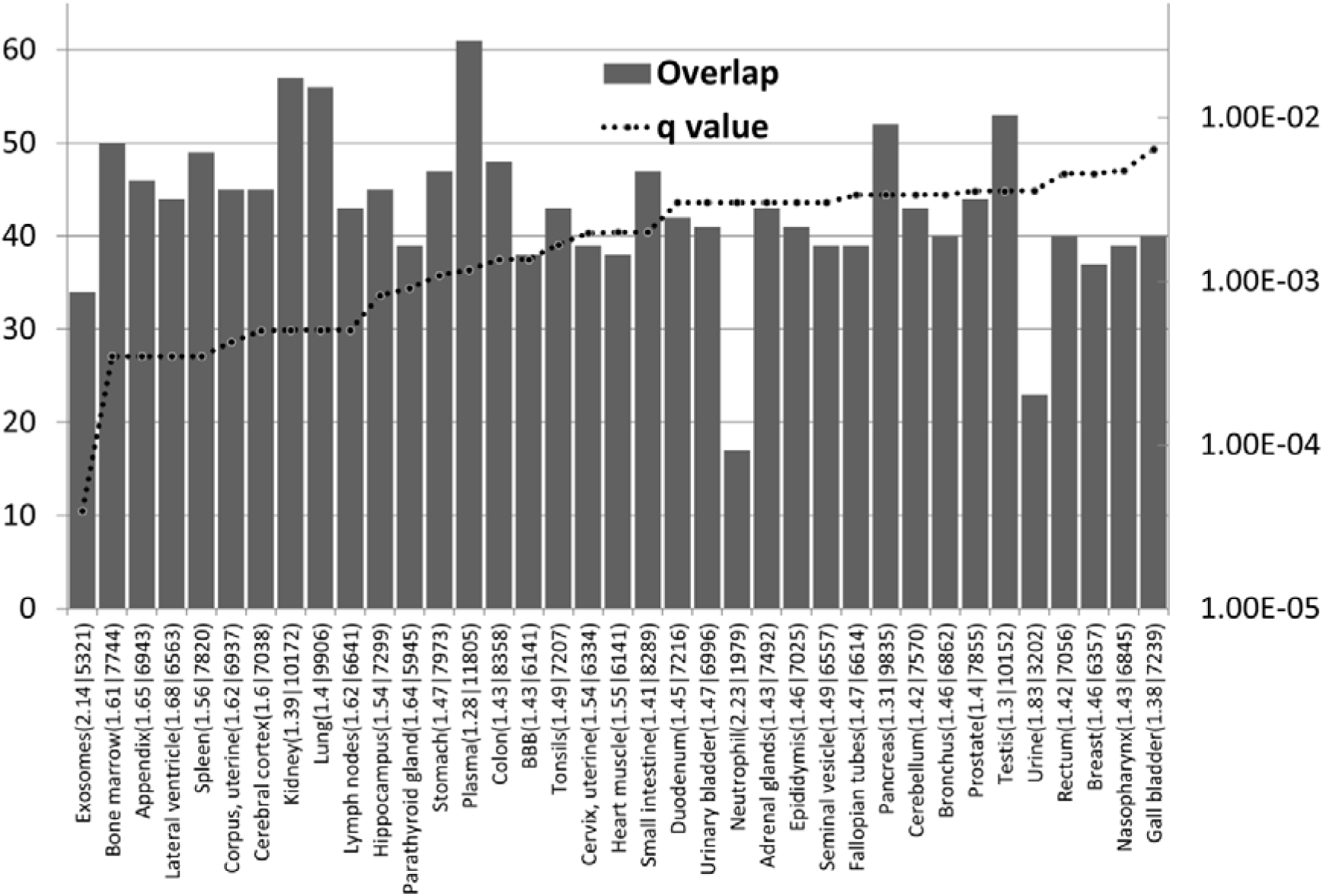
The distribution and enrichment of 78 AD genes in diverse proteomic and genomic datasets (Funrich and Exocarta data). The bars indicate the number of genes (from 78) in each tissue and the dotted line the corrected p value (q value). The maximum on this axis is set to q = 0.05. Observed/expected values, followed by the total number of genes expressed are appended after the identities of each sample. BBB refers to a separate blood brain barrier proteomics dataset. Cancer or cell line datasets are omitted and the data are limited to anatomical datasets containg more than 10 AD genes (Not all data are shown).

The AD genes are most significantly enriched in the exosome and bone marrow datasets. As noted above, exosomes are hijacked by many viruses for intercellular spread. Exosomes are prevalent in plasma [35](also enriched in AD genes) and are also the means by which intracellulary generated beta-amyloid is conveyed to the extracellular space [36]. In this context, and in relation to the antimicrobial effects of beta-amyloid, APP and gamma-secretase are highly expressed in the immune dendritic cells that scout for invading pathogens [7]. The bone marrow is the hematopoietic source of red and white blood cells and platelets [37]. B cells in the bone marrow rapidly respond to infection [38] and the bone marrow is also a source of angiogenic cells that are involved in vascular endothelial repair, a process that is disrupted in Alzheimer’s disease [39,40]. The parathyroid gland expresses many AD genes and also plays a role in hematopoesis [41,42]. Other immune related areas enriched in AD genes include the appendix, spleen, tonsils, the lymph nodes and the bronchus and neutrophils. The appendix is an important component of mucosal immune function, particularly B cell-mediated immune responses and extrathymically derived T-lymphocytes [43]. The tonsils and nasopharynx, also enriched in AD genes, play an important role in the initial defence against respiratory pathogens [44].

AD genes are enriched in the lateral ventricle, a site of the choroid plexus [45]. This provides cerebrospinal fluid (CSF) and is the location of the blood-CSF barrier, which is exploited by pathogens to gain access to the brain. The choroid plexus plays an important role in pathogen defence [46]. Post-mortem gene expression studies of the choroid plexus epithelium in AD patients show changes indicative of increased permeability of the blood-cerebrospinal fluid barrierand a reduction of macrophage recruitment [47], factors that woud favour pathogen entry and reduce their phagocytosis by macrophages. The hippocampus bulges into the temporal horn of the lateral ventricle [48] and this area, a keystone of AD pathology, is thus in close proximity to a major site of cerebral pathogen entry. AD genes are also enriched in a separate BBB dataset from mouse cerebral arteries. This is discussed in greater detail below. Other barriers in intestinal and pulmonary tissues, also enriched in AD genes (Fig 1), might also be considered as potential sites of pathogen entry. Immune systems play an important role at barrier interfaces [49].

Although AD genes are expressed in other sites, the main focus, in terms of enrichment, relates to immune and barrier systems.

A number of the 78 AD genes (referenced in supplementary Table 1) are primarily concerned with immune function (**HLA-DRB1, HLA-DRB5, HMHA1, IGH**) while many others with diverse primary effects also possess relevant properties in relation to the immune system (**ACE, ADAMTS20, AP2A2, BCL3, BIN1, CR1, CLU, CUGBP2, DISC1, EPHA1, GAB2, INPP5D, MEF2C. MS4A3, MS4A4A, RIN3, SCIMP, SPPL2A, STK24, TREM2, TREML2, ZNF224**) or pathogen defence (e.g. phagocytosis or autophagy) (**ABCA7, APOC1, APOE, BCAM, CD2AP, CD33, CDON, CELF1, PAX2, PTK2B, SASH1, SQSTM1**). A number of the AD genes also act as primary receptors for pathogens. These include the poliovirus receptor **PVR**, the HSV-1 receptor **PVRL2**, and complement receptor (**CR1**), which binds to many opsonised pathogens but which may also act as an entry receptor for *Plasmodium falciparum*, *Legionella pneumophila and Mycobacterium tuberculosis*. **CD33** binds to the HIV-1 gp120 protein and to diverse forms of sialic acid which coats many pathogens. Others bind bacterial lipopolysaccharides (**APOC1** and **TREM2**) or the Escherichia coli cytotoxic necrotizing factor 1 (**BCAM**). Others (**AP2A2**, **BIN1, CD2AP**, and **PICALM**) are involved in endocytosis, an obligate requirement for pathogen entry following binding to cognate receptors (see supplementary Table 1 for references).

### Host/pathogen interactomes are enriched in AD genes (Fig 2)

**Fig 2.**
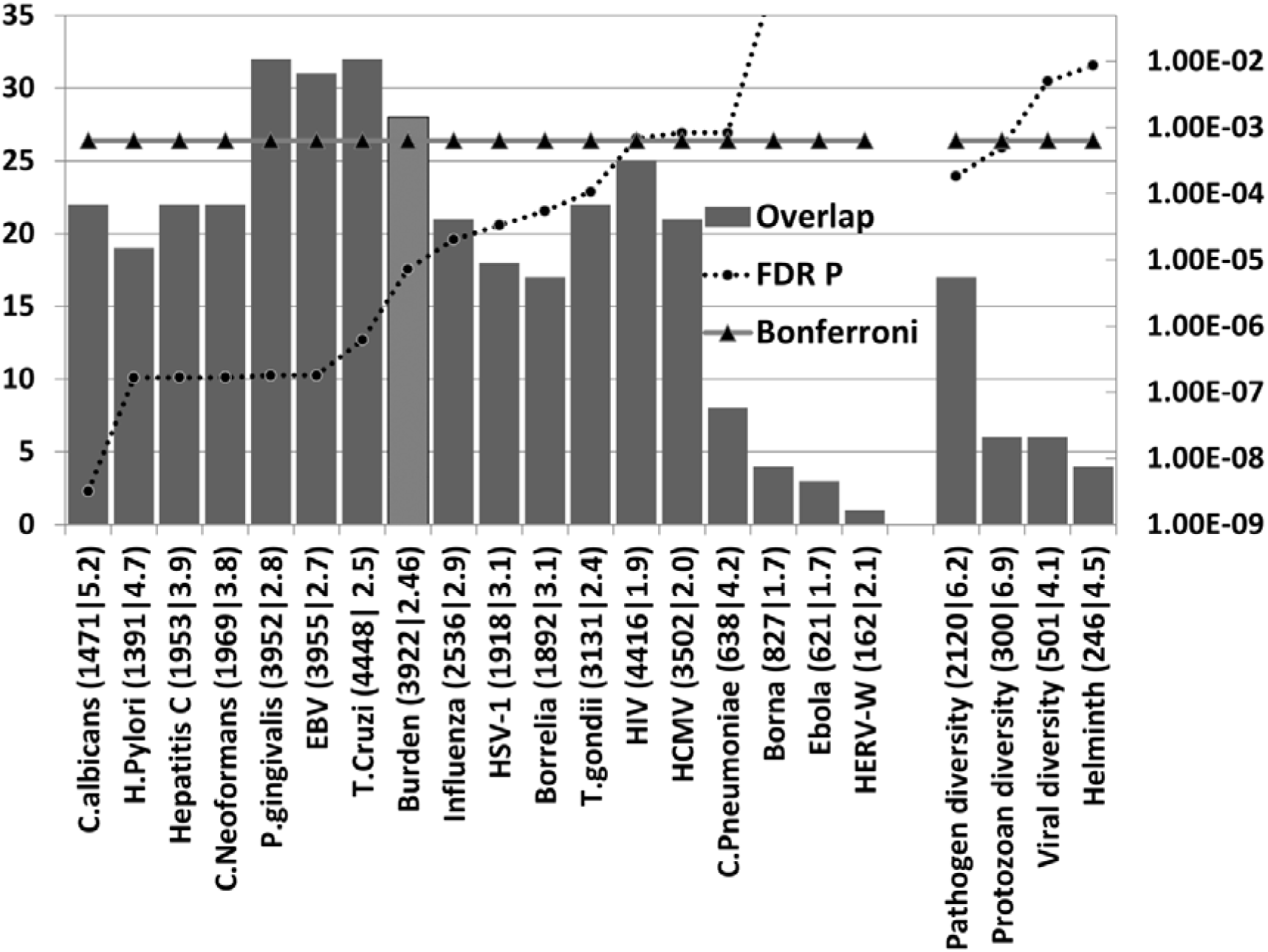
The number of AD genes (of 78) overlapping with diverse host/pathogen interactomes, or with those implicated in pathogen, protozoan or viral diversity or with the immune response to parasitic worms (Helminth) (Bars). The identities on the X-axis (e.g. *C. albicans* (1471|5.2) are appended with the total number of genes in each interactome (1471 in this case) or genetics dataset followed by the enrichment ratio (5.2 fold). The FDR-corrected p value for enrichment, derived from the hypergeometric distribution, is shown on the right hand axis (log scale) which is set to a maximum of 0.05. Invisible points are above this value. The Bonferroni cut-off level (p=0.05/78) is also shown. The Burden data (lighter shaded bar) correspond to the combined interactomes and AD gene overlaps of the human cytomegalovirus (HCMV), HSV-1, *Borrelia burgdorferi*, *Chlamydia pneumoniae* and *Helicobacter pylori*. EBV= Epstein-Barr virus.

All host/pathogen interactomes, with the exception of those of the Borna virus, Ebola virus and the HERV-W retrovirus were significantly enriched in AD genes (FDR p <0.05) with all but HIV-1, the cytomegalovirus and *C. pneumoniae* below the Bonferroni corrected value (P=6.41E-4). Pathogen burden (cytomegalovirus, HSV-1, *B. burgdorferi*, *C. Pneumoniae* and *H. Pylori*) has been associated with Alzheimer’s disease [1] and the pooled interactomes of these five pathogens (3922 host genes) were significantly enriched in AD genes (p= 7.3E-6). Given the variety of pathogens reported in AD brains (Table 1) other cumulative effects might be expected for various permutations.

The most significant pathogens related to fungi (*C. albicans* and *C. Neoformans*), the gum disease pathogen *P. Gingivalis* and the Epstein-Barr and hepatitis C viruses. Numerous fungal species, including *C.albicans*, have been detected in the AD brain (Table 1), although *C. Neoformans* was not one of the species studied. Two case reports have demonstrated virtually complete recovery from long-term (3 years) mis-diagnosed dementia/Alzheimer’s disease following antifungal treatment for *C. Neoformans* infection [50,51].

The Epstein-Barr virus has been associated with AD and hepatitis C associated with dementia (table 1). *In vivo* studies for the Epstein-Barr and Hepatitis C viruses are however limited by their inability to infect rats or mice. Several of these pathogens including *C. pneumoniae*, HSV-1, cytomegalovirus and the Epstein-Barr and hepatitis C viruses or *H. pylori* and *B. Burgdorferri* [52–59], periodontitis and *P.Gingivalis* [57]have also been associated wiith atherosclerosis, an important endophenotype in AD [60].

Apart from **APOE4** no AD genetic variants seem to have been studied in relation to effects on pathogens and it is impossible to note whether the variants favour or oppose their destructive potential. The apolipoprotein E (**APOE4**) variant protects against hepatitis C [61], but favours the cerebral entry of HSV-1 [62]and enhances the attachment of *C. pneumoniae* elementary bodies to host cells [63].

### AD genes overlap with those implicated in pathogen, protozoan or viral diversity or with the immune response to parasitic worms (Fig 2)

The AD genes are enriched in a series of genome-wide and global-wide datasets related to general pathogen diversity, protozoan or viral diversity (the number of different pathogens in a geographic region) or with the immune response to parasitic worms, most significantly so for general pathogen and protozoan diversity (FDR p < 0.05). The overlaps in relation to viral diversity or the response to parasitic worms exceeded the Bonferroni cut-off.

In evolutionary terms, these pathogen-related genes likely reflect pathogen resistance rather than susceptibility [17–20].

It has also been noted that genes related to inflammatory diseases [64] or to the AD gene network [65]are subject to positive selection pressure. While many pathogens have been implicated in AD, the selection of AD genes for pathogen resistance rather than susceptibility seems logical in relation to several considerations, as already proposed [66,67]. Firstly, the old age of AD patients indicates survival from the many infectious diseases that are among the principal causes of death in adults and children. In the USA, the leading non-accidental causes of death in adults (2013 figures) include heart disease; cancers; chronic lower respiratory diseases; cerebrovascular diseases; Diabetes mellitus; Influenza and pneumonia; nephritis, nephrotic syndrome and nephrosis [68].

Certain viruses, helminths and bacteria are oncogenic and it has been estimated that 15-20% of cancers are due to infections [69]. The inverse association between the incidence of cancer and Alzheimer’s disease [70] suggests that AD genes might well be cancer protective (but also that death due to cancer precludes AD). Inflammatory heart diseases [71] and atherosclerosis, cerebrovascular disorders and stroke have also been linked to infection [72,73]. Enteroviruses have been implicated in Type 1 diabetes mellitus [74].

The leading non-accidental causes of infant deaths were congenital malformations, deformations and chromosomal abnormalities; disorders related to short gestation and low birth weight, not elsewhere classified; newborn affected by maternal complications of pregnancy; sudden infant death syndrome; newborn affected by complications of placenta, cord and membranes; bacterial sepsis of newborn; respiratory distress of newborn; diseases of the circulatory system; and neonatal haemorrhage. Again, many of these relate to infections. In evolutionary terms, pandemics and infectious diseases have been, and in poorer countries still are, associated with high mortality.

In relation to Alzheimer’s disease, the apolipoprotein E (**APOE4**) variant protects against malaria [75] and hepatitis C [61], although **APOE4** favours cerebral entry of the herpes simplex virus [62]and enhances the attachment of Chlamydia pneumoniae elementary bodies to host cells [63]. Malaria and hepatitis C are both associated with high mortality [76,77] and the protective effects of **APOE4** would encourage its maintenance in the population, to the detriment of infection by the less virulent agents.

The **APOE4** variant is also associated with enhanced immune/inflammatory responses. For example, Toll-like receptor activation (TLR3, 4) in microglia induces cyclooxygenase-2 (PTGS2), microsomal prostaglandin E synthase (PTGES), and prostaglandin E2, an effect exaggerated in **APOE4/APOE4** mice [78]. **APOE4** is also associated with enhanced *in vivo* innate immune responses in human subjects. Whole blood from healthy APOE3/**APOE4** volunteers induced higher cytokine levels on ex vivo stimulation with Toll-like receptor (TLR2, 4 or 5) ligands than blood from APOE3/APOE3 patients [79]. Gain of function also applies to AD variant forms of complement receptor **CR1**, which are better able to bind complement component C1q or C3B [80]. C1q and C3B are opsonins that interact with complement cell-surface receptors (C1qRp, **CR1**, CR3 and CR4) to promote phagocytosis (including that of infectious agents) and a local pro-inflammatory response [81]. **TREM2** variants in AD are also associated with enhanced inflammatory responses (upregulation of proinflammatory cytokines) [82]. In presenilin (**PSEN1**) mutant knockin mice, microglial challenge with bacterial lipopolysaccharide results in enhanced nitric oxide and inflammatory cytokine responses, relative to normal mice [83]. For these genes at least, this gain of immune/inflammatory function concords with selection for pathogen resistance.

It has also been noted that unaffected offspring with a parental history of AD have an enhanced inflammatory response in lipopolysaccharide-stimulated whole blood samples, producing higher levels of interleukin 1beta, tumor necrosis factor alpha and interferon gamma in response to LPS. This effect was independent of the **APOE4** variant [84] suggesting that other AD genes are also endowed with gain of function in relation to the immune/inflammatory system. Monocyte-derived dendritic cells from Alzheimer’s disease patients also produce more interleukin 6 than those from healthy controls. AD monocytes stimulated with LPS also show a higher induced expression of the pro-inflammatory ICAM-1 adhesion molecule than controls [85]. Beta-amyloid also stimulates cytokine production in peripheral blood mononuclear cells (PBMC) and the production of the chemokines, RANTES, MIP-1beta, and eotaxin as well as that of CSF2 (colony stimulating factor 2 (granulocyte-macrophage)) and CSF3 (colony stimulating factor 3) is greater than controls in AD-derived PBMC stimulated with beta-amyloid [86].

Given the antimicrobial properties of beta-amyloid, any genetic variant that increase its production, at least in the periphery, might also be considered as desirable, in evolutionary terms, in relation to pathogen defence. A high percentage of AD GWAS genes are involved in APP processing [87]. The AD genetic variant of **ABCA7** results in increased secretion of beta amyloid and raised beta-secretase activity in CHO-and HEK cells with the Swedish APP mutation [88], but the effects of late-onset AD variant genes on the beta-amyloid response to pathogens remain to be determined.

### Host/pathogen interactome enrichment in misregulated genes of the Alzheimer’s disease hippocampal transcriptome (Fig 3)

**Fig 3.**
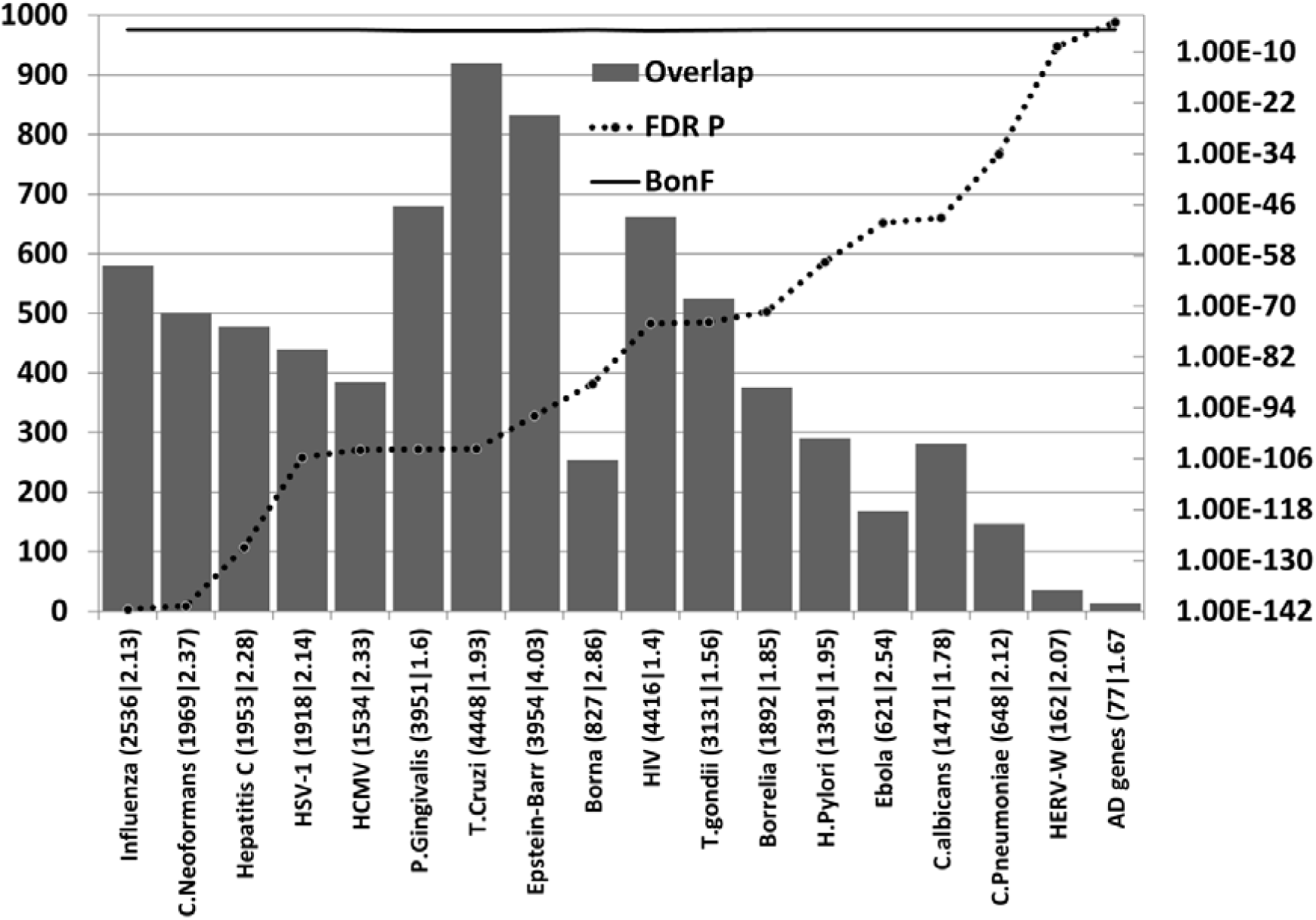
The number of genes misregulated (combined up and down) in a microarray study of the AD hippocampus overlapping with diverse host/pathogen interactomes. The identities on the X-axis (e.g. C. albicans (1471|5.2) are appended with the total number of genes in each interactome (1471 in this case) or genetics dataset followed by the enrichment ratio (5.2 fold). The p value for enrichment, derived from the hypergeometric distribution, is shown on the right hand axis (log scale) which is set to a maximum of 0.05. The Bonferroni cut off (1.74E-05) is also shown.

All pathogen interactomes, most notably relating to influenza, *C. Neoformans* and Hepatitis C were highly enriched in genes relating to this microarray dataset (combined up and downregulated genes). The significance level of the interactome enrichment for most pathogens was several orders of magnitude below the Bonferroni cut off (p=1.74E-05) (Fig 3). 14/78 AD genes appear in this microarray dataset (FDR p = 0.001). Two case reports have demonstrated virtually complete recovery from long-term (3 years) mis-diagnosed dementia/Alzheimer’s disease following antifungal treatment for C. Neoformans infection [50,51]. Regarding the influenza data, bronchopneumonia, often caused by influenza, is a common final cause of death in dementia patients [89] and such recent infections close to death may well influence the data.

Regardless of the rank order, it is clear that many diverse pathogen interactomes affect several hundred genes of the 2879 misregulated in the AD hippocampus and/or that these misregulated AD genes represent a substantial percentage of the individual pathogens' interactomes (Fig 3).

Kegg pathway analysis of these misregulated hippocampal genes using the consensus path database [90] showed that many infection-related pathways were also significantly enriched (FDR p < 0.05). These included (pathogen with N genes followed by the FDR corrected p value): Epstein-Barr virus infection (74,5.5E-7); Salmonella infection (36,0.0001); Tuberculosis (57,0.0009); Epithelial cell signaling in Helicobacter pylori infection (28,0.00097); Shigellosis (27,0.001); Influenza A (54,0.003); Herpes simplex infection (56,0.0036); Vibrio cholerae infection (21,0.0089); HTLV-I infection (71,0.0096); Toxoplasmosis (37,0.013); Hepatitis B (43,0.018); Pathogenic Escherichia coli infection (20,0.02); Bacterial invasion of epithelial cells (26,0.02); Measles (38,0.04).

### Upregulated genes in the AD hippocampus are enriched in genes upregulated by multiple viral, bacterial and fungal pathogens or Toll-like receptor ligands

Numerous infection-related microarray datasets exist in the Molecular signatures database or in the Gene expression omnibus (see methods), using blood cells taken from infected patients, or cells or tissues infected under laboratory conditions.

The hippocampal genes upregulated in Alzheimer’s disease were significantly enriched in upregulated genes in datasets from multiple viral species and to double stranded RNA and the viral mimic/TLR3 agonist, Polyinosinic:polycytidylic acid (poly I:C) (Fig 4). The viruses ranged from the benign (e.g. the rhinovirus that causes the common cold) to the highly malignant (e.g. the ebolavirus, rabies virus or HIV-1). They include common human infectious agents (e.g. adenovirus 5, influenza, Epstein-Barr virus, herpes simplex virus (HSV-1), measles or the Norwalk virus). Apart from HSV-1, the human cytomegalovirus, HIV-1 or hepatitis C (See Table 1) none of these have been implicated in Alzheimer’s disease or dementia. Most microarray experiments related to immunocompetent blood cells (B cells, T cells, dendritic cells, monocytes and macrophages) or to cultured cell lines. No infection-related datasets were found for microglia, the brain resident immunocompetent cells, but significant enrichment of the AD upregulated genes was observed for genes upregulated by interferon gamma in microglial cells (Fig 4). Interferon gamma plays an important role in the response to viral, bacterial and parasitic infections [91].

**Figure 4:**
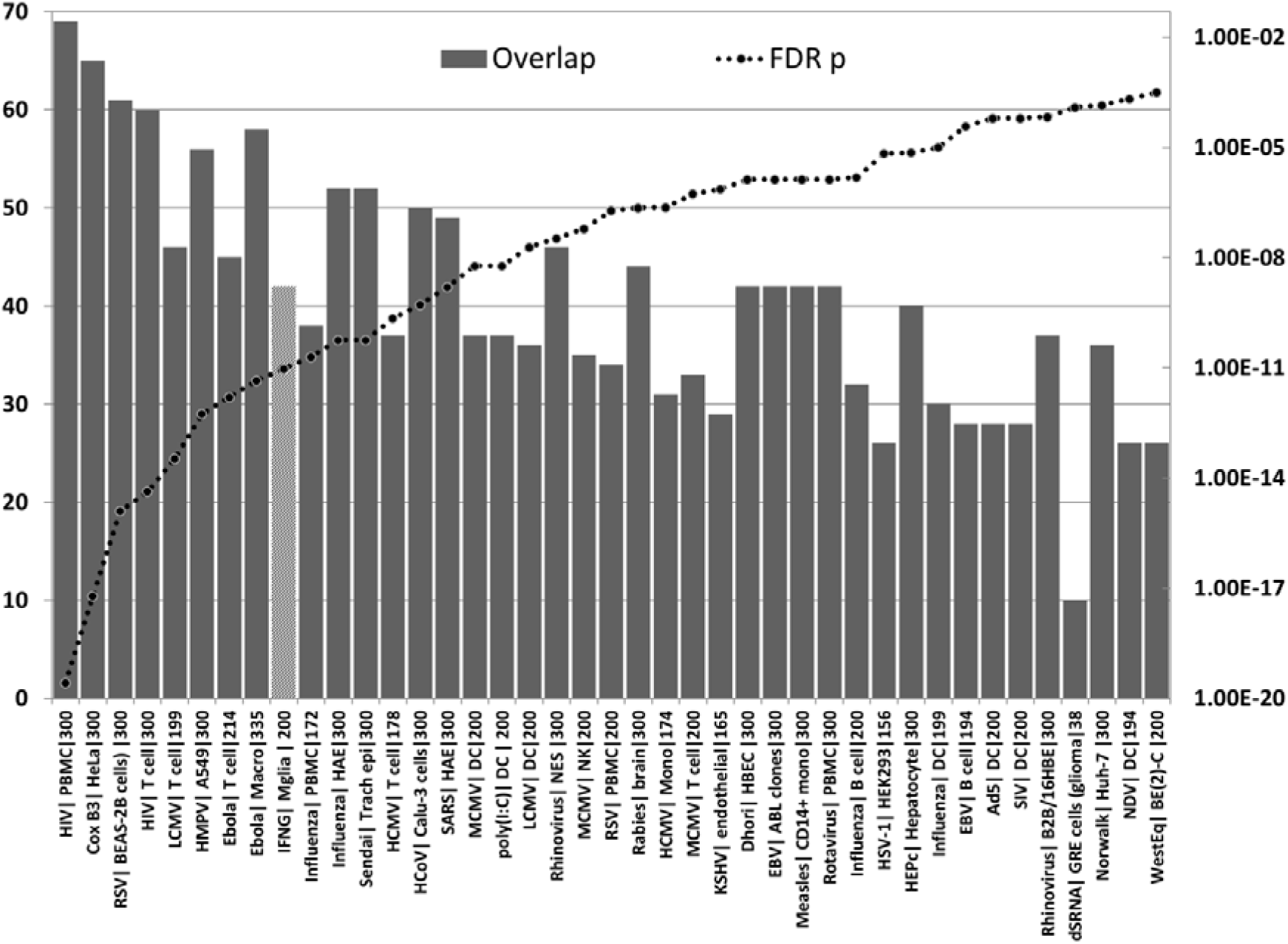
The number of upregulated genes (bars) from the AD hippocampal transcriptome that overlap with upregulated genes in viral infection datasets from the Molecular signatures database or the Gene expression omnibus (see methods). The effects of the mimic poly(I:C) are also shown, as is the effect of interferon gamma on gene expression in microglial cells. For each datapoint, the name of the virus is shown, followed by the cell type and the total number of upregulated genes in the viral datasets (limited by MSigDb or GEO). The significance of enrichment (right axis) represents the FDR corrected p value from the hypergeometric test. All values are below the Bonferroni corection (0.05/300 = 1.67E-04). Because the number of downloaded genes is mostly limited to 300, this is the maximum number of possible overlaps. The pale bar represents the microglial response to interferon gamma. Tissue/cell abbreviations; A549= adenocarcinomic human alveolar basal epithelial cells; ABL = Akata Burkitt’s lymphoma cells; B2B/16HBE, BE(2)C or BEAS-2B = human bronchial epithelial cells; BroLav = human bronchial lavage; Calu-3 = Cultured Human Airway Epithelial Cells; DC = dendritic cells; GRE = glioma cell line; HAE = human airway epithelial cells; HBEC = Human Bronchial Epithelial Cells; HEK = human embryonic kidney cells; HeLa = cervical cancer cell line; HuH-7 = hepatocarcinoma cell line; Macro = macrophage; Mgli = microglia; Mono =monocytes; NES = human nasal epithelial scrapings; NK = natural killer cells; PBMC = peripheral blood mononuclear cells; PLC/PRF/5 cells = human liver hepatoma cells; Trach epi = Tracheal epithelial cells Viral abbreviations (Reading from the left): HIV= human immunodeficiency virus, Cox B3 =Coxsackie B3 virus; RSV = respiratory syncytial virus; LCMV= Lymphocytic Choriomeningitis Virus; HMPV= Human metapneumovirus; Ebola= Ebola virus; Influenza = Influenza A virus; Sendai = Sendai virus, HCoV = human coronavirus; IFNG = interferon gamma; SARS = severe acute respiratory syndrome coronavirus; HCMV = human cytomegalovirus; MCMV = mouse cytomegalovirus; Dhori = Dhori virus; EBV = Epstein-Barr virus; HepC = hepatitis C virus; KSHV= Kaposi’s sarcoma-associated herpesvirus; HSV-1 = herpes simplex; Norwalk = Norwalk virus (Norovirus); Ad5 = adenovirus 5; SIV= Simian immunodeficiency virus; poly(I:C)= Polyinosinic:polycytidylic acid (a viral mimic stimulating Toll-like TLR3 receptors); NDV = Newcastle disease virus; WestEq = Western equine encephalomyelitis virus; LASV = Lassa virus; dsRNA = double stranded RNA; HEV = hepatitis E virus.

The upregulated hippocampal genes in AD were also enriched in infection datasets for numerous bacteria as well as to fungal species (*C. albicans* and *C.neoformans*) and in those related to bacterial endotoxin or sepsis and to nematode/trematode or protozoan infection datasets (FDR p < 0.05) (Fig 5). This also applied to diverse lipopolysaccharide datasets and responses to Toll-like receptor ligands, CpG oligonucleotide (a ligand for TLR9, which mediates cellular response to unmethylated CpG dinucleotides in bacterial DNA (definition from Refseq)) and R848 (a ligand for TLR7/TLR8 both of which recognize RNA released from pathogens that enter the cell by endocytosis [92]) (Fig). With the exception of *H.Pylori*, *P.Gingivalis* and *Borrelia burgdorferi* and *C.albicans* or *C.Neoformans*, none have been implicated in AD.

**Fig 5.**
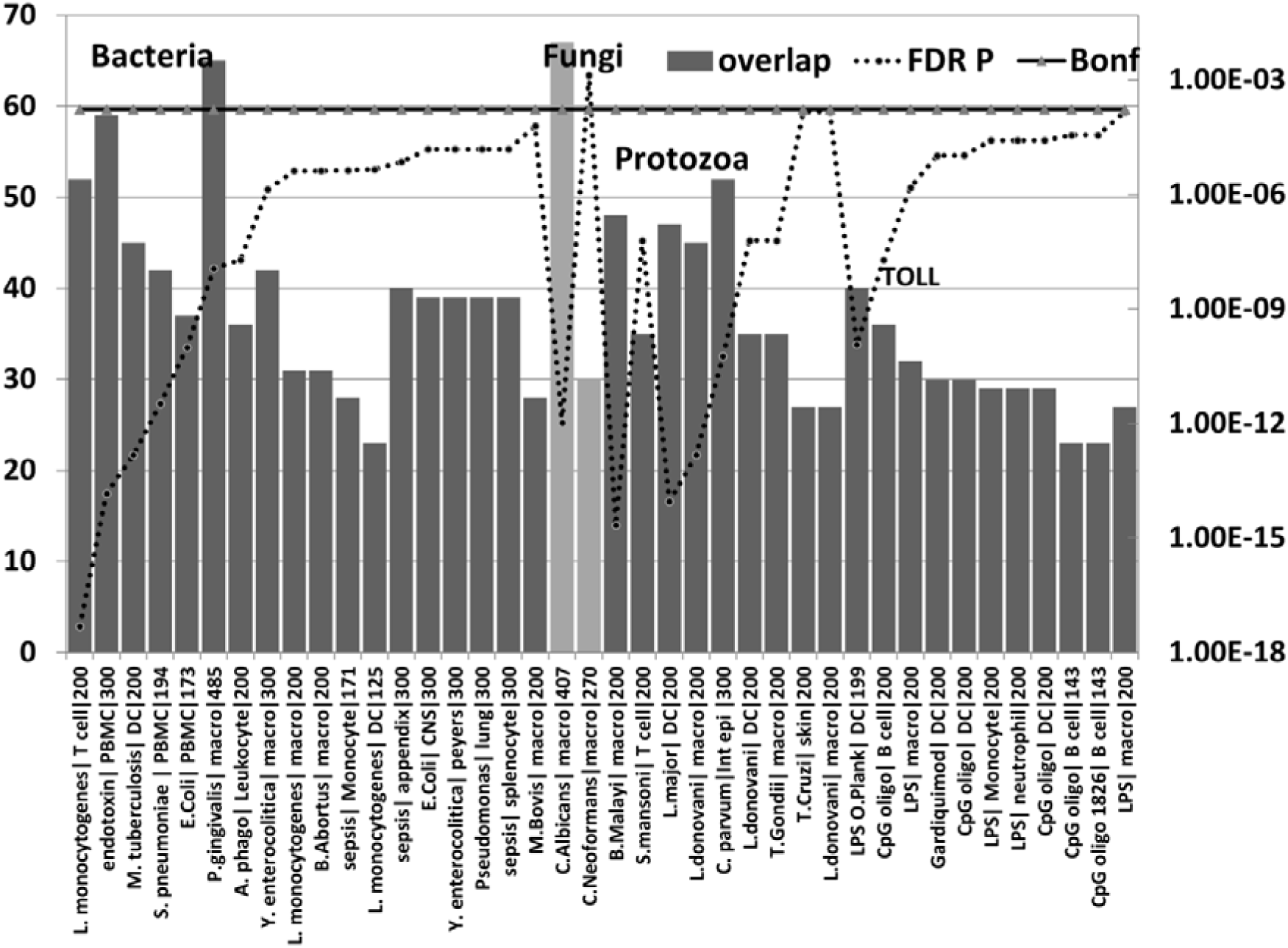
The number of upregulated genes (bars) from the AD hippocampal transcriptome that overlap with upregulated genes in bacterial (first batch), fungal (pale bar =*C.albicans*, *C.Neoformans*), nematode (B.Malayi) /trematode (S.Mansoni), or protozoan microarray datasets (see methods). The effects Lipopolysaccharides and other Toll receptor ligands are also shown. For each datapoint, the name of the pathogen or ligand is shown, followed by the cell type and the total number of upregulated genes in the comparator datasets (limited by MSigDb or GEO). The significance of enrichment (right axis) represents the FDR corrected p value from the hypergeometric test. All values except for *C.Neoformans* are below the Bonferroni correction level. Pathogen or ligand abbreviations (from left) *L. monocytogenes* = *Listeria monocytogenes*; endotoxin = gram-negative bacterial wall component; *S.pneumoniae* = *Streptococcus pneumoniae*; *E.Coli* = *Escherichia coli*; *P. gingivalis* = *Porphyromonas gingivalis*; *A.phago* = *Anaplasma phagocytophilum*; *Y. enterocolitica* = *Yersinia enterocolitica*; *M.Bovis* = *mycobacterium bovis*; *C.albicans* = *Candida albicans*, *C.Neoformans* = *Cryptococcus neoformans*, *B.Malayi* = *Brugia malayi* (filarial parasite causing elephantiasis); *S.mansoni* = *Schistosoma mansoni*; *L. donovani* = *Leishmania donovani*; *C. parvum* = *Cryptosporidium parvum*; *L. Major* = *Leishmania major*; *T.Gondii* = *Toxoplasma Gondii*; *T. Cruzi* = *Trypanosoma Cruzi*; LPS = lipopolysaccharide; LPS O.Plank = Oscillatoria Planktothrix (cyanobacteria lipopolysaccharide) CpG oligo = CpG Oligodeoxynucleotide (TLR9 ligand); Gardiquimod = TLR7 ligand; Cell type abbreviations as for Fig 4. CNS = central nervous system; peyers = peyers patch; Int epi = intestinal epithelial cells;

Together these data suggest a significant parallel between the upregulated genes in the AD hippocampus and the responses to multiple and diverse infectious agents with little overall discrimination between viral, bacterial, fungal or protozoan types of infection. Multiple pathogens have been detected in the AD brain (see Table 1) and the diversity of these infection related overlaps with the AD hippocampal transcriptome suggests that many other pathogens could induce similar pathological transcriptome changes. Microbiome studies in the AD brain and periphery will help to elucidate the role of multiple pathogens.

### Pathogen interactomes are enriched in the proteins found in AD amyloid plaques and neurofibrillary tangles (Fig 6)

**Fig 6.**
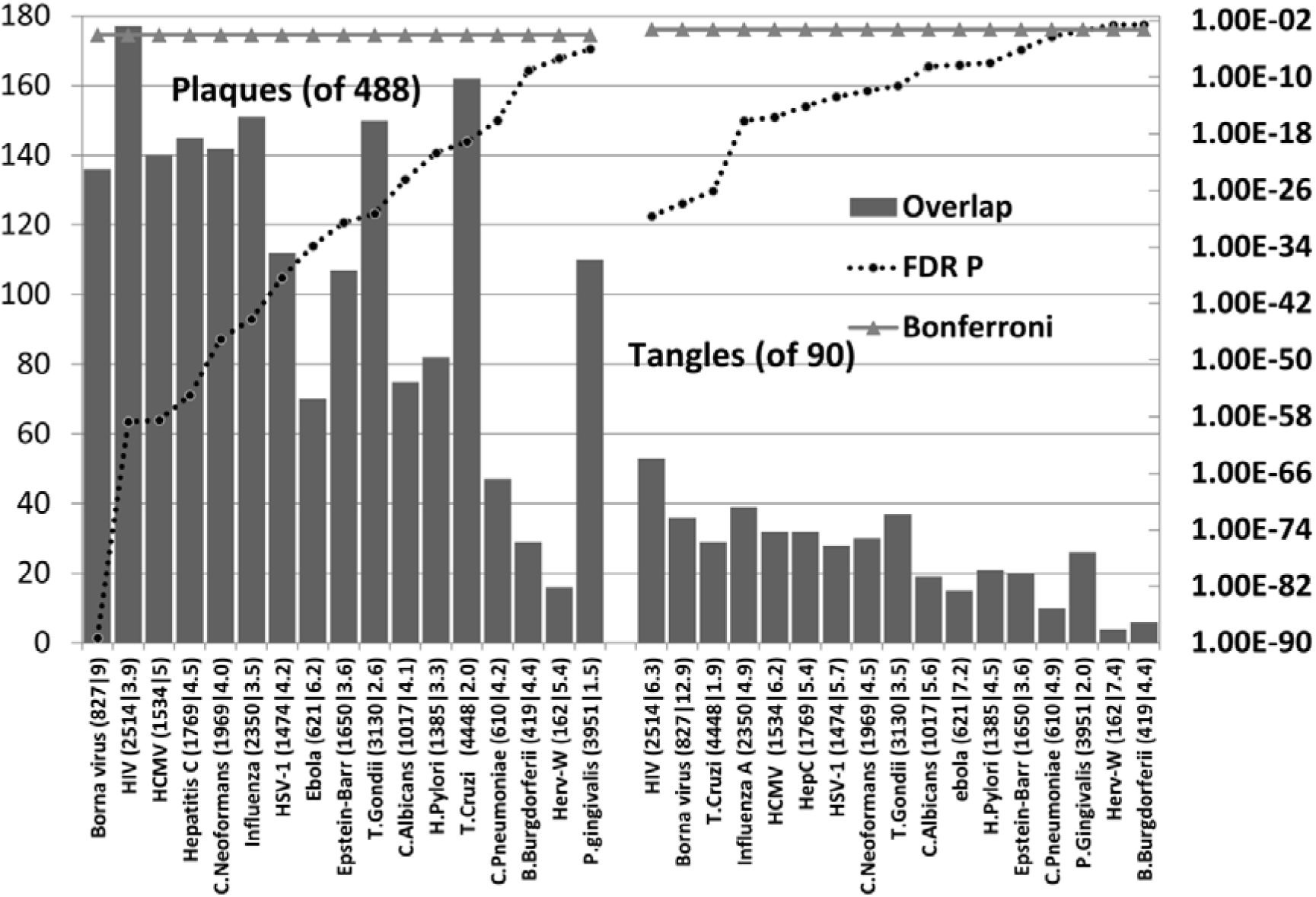
Host pathogen interactome enrichment in a set of 488 proteins isolated from amyloid plaques in the AD brain or from 90 proteins isolated from neurofibrillary tangles. The identities on the X-axis are appended with the total number of genes in each interactome followed by the enrichment ratio. The FDR p value for enrichment, derived from the hypergeometric distribution, is shown on the right hand axis (log scale) which is set to a maximum of 0.05.

All pathogen interactomes were significantly enriched in proteins found in plaques and all except HERV-W and *B.Burgdorferi* interactomes significantly enriched in tangle proteins (below the Bonferroni cut-off level). The Borna virus and HIV-1 ranked highly in both cases. There is only one publication relating to Borna virus effects on beta-amyloid and none could be found for tangles. The microglial activation produced by the virus reduced brain parenchymal, but increased cerebral vascular beta-amyloid deposition, in APP transgenic mice [93]. The top agents relating to plaques were predominantly viral, while those relating to tangles were mostly viral, but included the parasites, *T. Cruzi* and *T.Gondii*.

Beta-amyloid is an antibacterial, antifungal and antiviral agent (Table 1). It has been shown that it binds to *C.albicans* and *S.Typhimurium* [2] and presumably to other microbes. Such microbes may well have sequestered host proteins specific to their particular life cycles during their passage to the cell, and this would partly explain the interactome enrichment. In addition to the plaque proteins relating to pathogen life cycles (for example receptor binding, endocytosis and transport between intracellular compartments or nuclear entry and subsequent translation in the case of HSV-1), the proteins found in plaques and tangles contain many related to the immune system, inflammation and autophagy, all of which play a general role in pathogen defence [8,24,25] as does beta-amyloid.

Viruses are transported via the microtubule network [94], which is also exploited by *C.Pneumoniae*, *T.Cruzi* and *T.Gondii* to reorganise cellular organelles to the pathogens' advantage [95,96]. Phosphorylated tau is a hallmark of neurofibrillary tangles and is induced by many pathogens (Table 1). Tau phosphorylation can also be induced by interferon gamma, an effect related to disinhibition of glycogen synthase kinase [97]. It is not clear whether or how such effects could influence the pathogens.

### AD genes are localised in the Blood brain barrier

30/78 AD genes are expressed in the BBB proteome dataset of mouse cerebral arteries [28] (Fig 1). The list below indicates the 30 BBB genes, annotated with the number of pathogen interactomes with which they overlap. Most BBB expressed genes interact with none or few pathogens (5 or less of the 17 studied), suggesting a subdivision of mainly BBB and mainly pathogen related. This could of course be confounded by missing data, as several of these genes are poorly characterised in terms of function. These 30 genes (N interactomes in brackets) are:- **PCNX (0), ABCA7 (1), ADAMTS20 (1), ATXN7L1 (1), TREML2 (1), AP2A2 (2), BCAM (2), CNTNAP2 (2), ECHDC3 (2), FRMD4A (2), GRIN3B (2), PAX2 (2), PICALM (2), DISC1 (3), LUZP2 (3), RELN (3), TTLL7 (3), FERMT2 (4), HMHA1 (4), MSRA (4), PPP1R3B (4), SASH1 (4), BIN1 (5), SORL1 (5), PVRL2 (7), MMP12 (8), CLU (9), PTK2B (10), BCL3 (13), SQSTM1 (13)**.

The BBB location of a high proportion of AD genes indicates an important function in relation to AD. Several studies have reported that disruption of the blood brain barrier is an important feature of AD [98–101]. Cerebral microbleeds and cortical siderosis (an increase in blood-derived iron deposition) are a feature related to BBB disruption in AD patients [102–104]. Many bacteria depend upon the availability of free iron and such effects may contribute to their sucessful colonisation in AD [105].

### Other environmental risk factors in AD disrupt the blood-brain barrier and BBB integrity is maintained by beneficial factors

AD susceptibility genes might have been selected for pathogen resistance rather than susceptibility (see above). In which case, what are the factors, in the aged, that nevertheless permit the cerebral invasion of a large variety of pathogens? (See Table 1) Certain viruses (e.g. HSV-1) can enter the brain via the olfactory or other neural routes, exploiting an ability to use the axonal transport system [106]. Some parasites [107] and bacteria (e.g. *C. Pneumoniae* [108,109]) have also found ways to circumvent the barrier systems that usually protect the brain.

Aging itself leads to blood brain barrier dysfunction [110] and immunosenescence is also a feature of ageing and AD. However, while immunosenescence can increase susceptibility to pathogens due to immunodefficiency, it is also accompanied by an increase in the pro-inflammatory activity of monocytes and macrophages which can lead to chronic low grade inflammation, termed “inflammageing”[111,112]. This increased inflammatory function also applies to microglia, the macrophage-like cells in the brain [113]. Certain AD gene variants are associated with enhanced pro-inflammatory responses (see above) and cerebral pathogen entry would thus be met with a doubly vigorous inflammatory response related to both immunosenescence and genetic variation. Persistently activated monocyte/macrophages have been observed in the blood of patients with early AD [114] and increased activation of microglia/macrophages, colocalized with the area of heavy beta-amyloid concentration, is also observed in the brains of AD patients [115].

Apart from pathogens, many other environmental risk factors have been reported in AD. These include diabetes, midlife hypertension or obesity, smoking and physical inactivity [116]. Other contributory factors include previous head injury [117], exposure to toxic metals (aluminium [118,119] or copper[120]), pesticides (organochlorine and organophosphate insecticides) [121,122], industrial chemicals (flame retardants) and air pollutants (particulate matter and ozone [123–126]). High levels of cholesterol or homocysteine [127–130]and low levels of folic acid [131,132] have also been associated with AD. In relation to cholesterol, atherosclerosis of the carotid arteries or of leptomeningeal vessels and in the circle of Willis has also been observed in AD. Such atherosclerotic effects can lead to chronic cerebral hypoperfusion [60,133,134].Sleep disruption or obstructive sleep apnoea are also associated with AD risk [135,136].

Factors reported to be of benefit, or that reduce the incidence of AD include the use of non-steroidal anti-inflammatories (NSAIDs) [137,138], and the early use of statins [139–141]. Statins also have antimicrobial effects against oral microorganisms including *Aggregatibacter actinomycetemcomitans* and *P. Gingivalis*, and against most dental plaque bacteria, including *Streptococcus mutans*. They possess antiviral properties against the human cytomegalovirus, HSV-1, hepatitis B and C viruses, and antifungal properties against *Candida albicans*, *Aspergillus fumigatus*, and Zygomycetes species [142].

Beneficial dietary factors in AD include caffeine [130], chocolate (versus cognitive decline in the nondemented aged)[143]) and the Mediterranean diet [144–146]. Melatonin [147,148], estrogen [149–151]and memantine [152,153] also have reported benefits in AD.

The environmental risk factors associated with AD disrupt the BBB, and BBB integrity is maintained by the beneficial factors (Table 2). While infections are random uncontrollable events, many of the other environmental risk factors are modifiable by lifestyle changes, for example diet, obesity, smoking and exercise, and it has been estimated that addressing such modifiable risk factors might result in a significant reduction in the incidence of AD [116]. Amelioration of BBB disruption has already been proposed as a potential therapy in AD, and several drugs including angiotensin receptor blockers, etodolac (NSAID), granisetron (5HT3 serotonin receptor antagonist) or beclomethasone (corticosteroid) [154,155] as well as other NSAIDS, statins and other drugs referenced in Table 2 might be considered as suitable candidates.

**Table 2.**
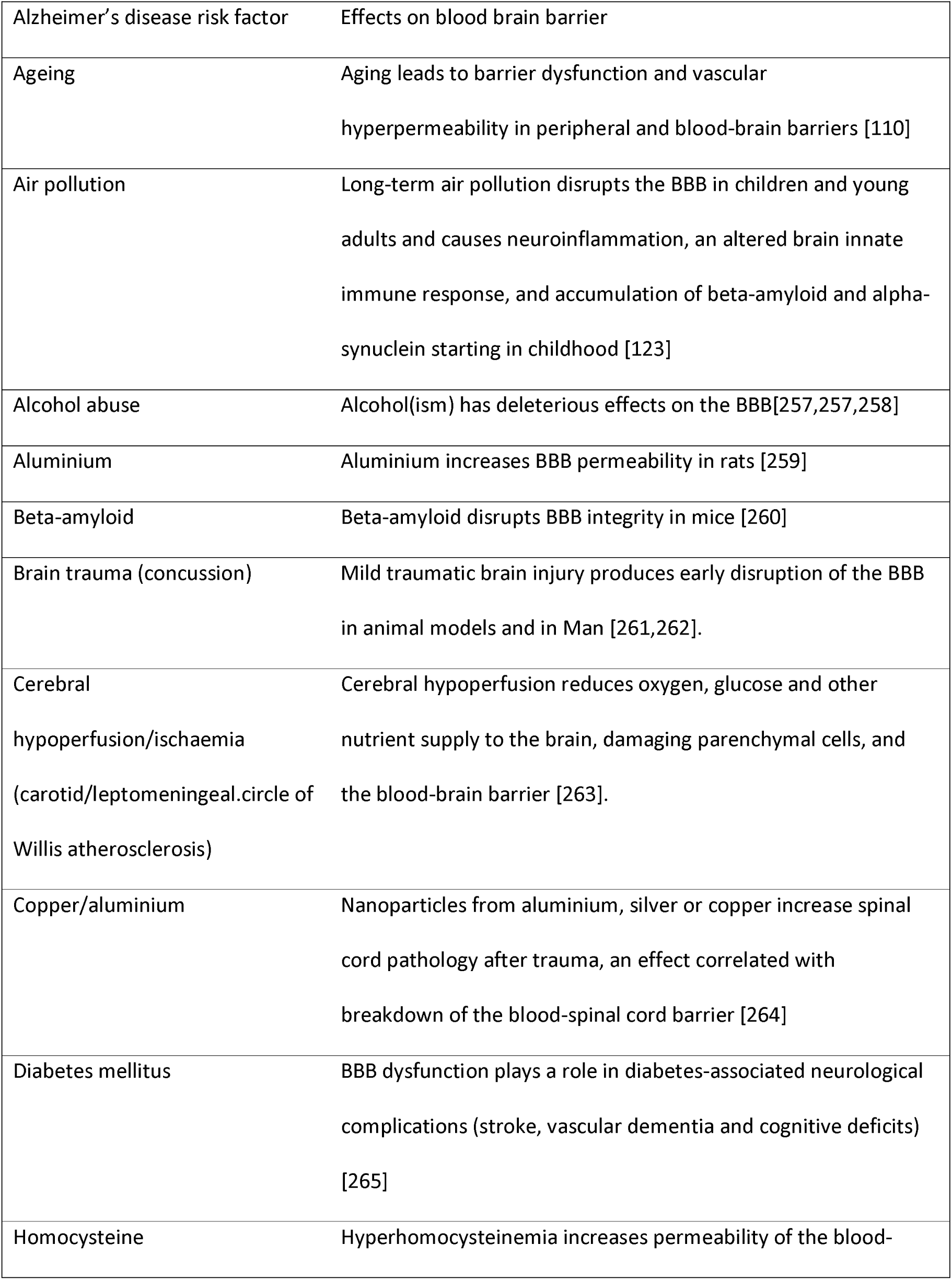
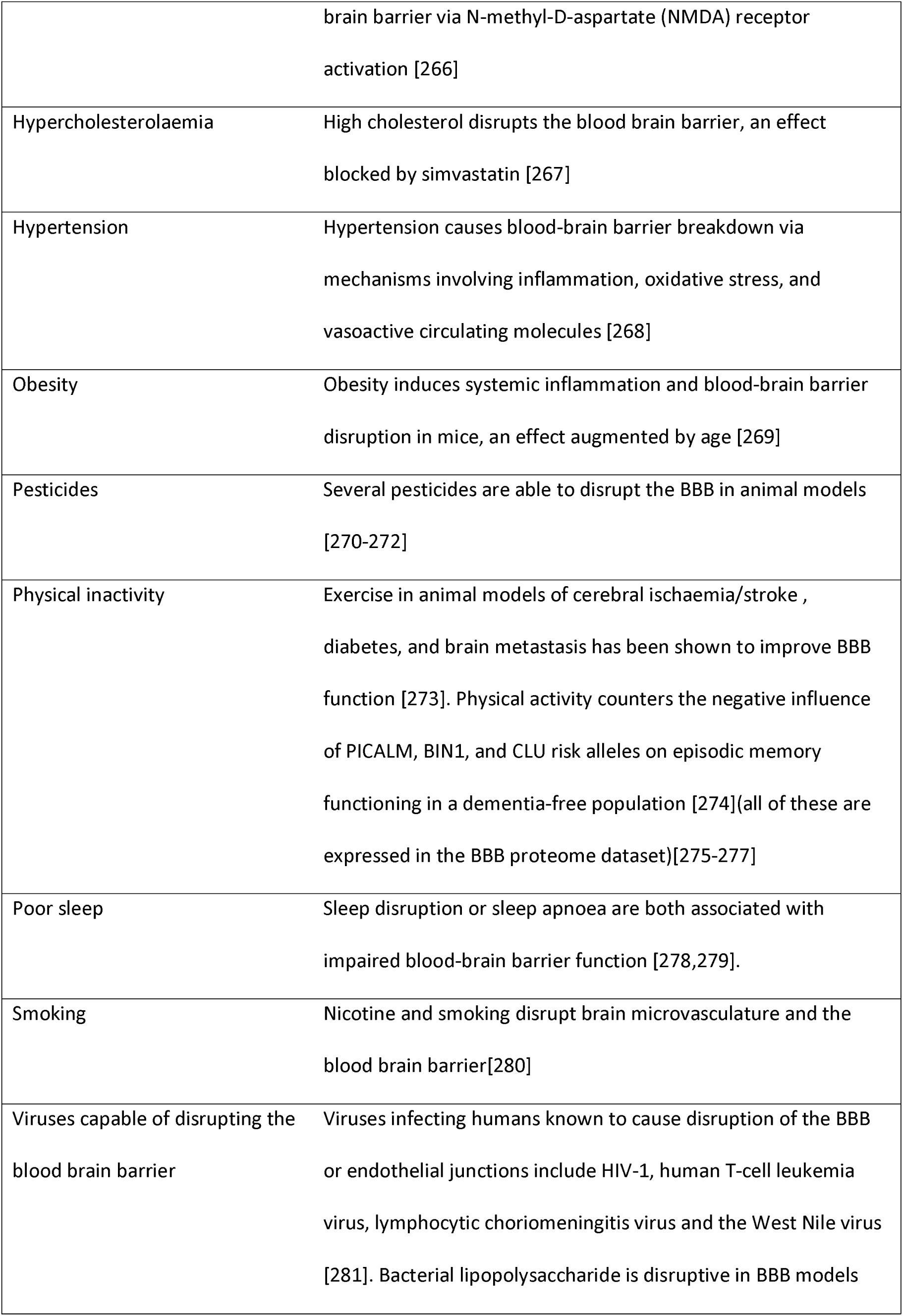
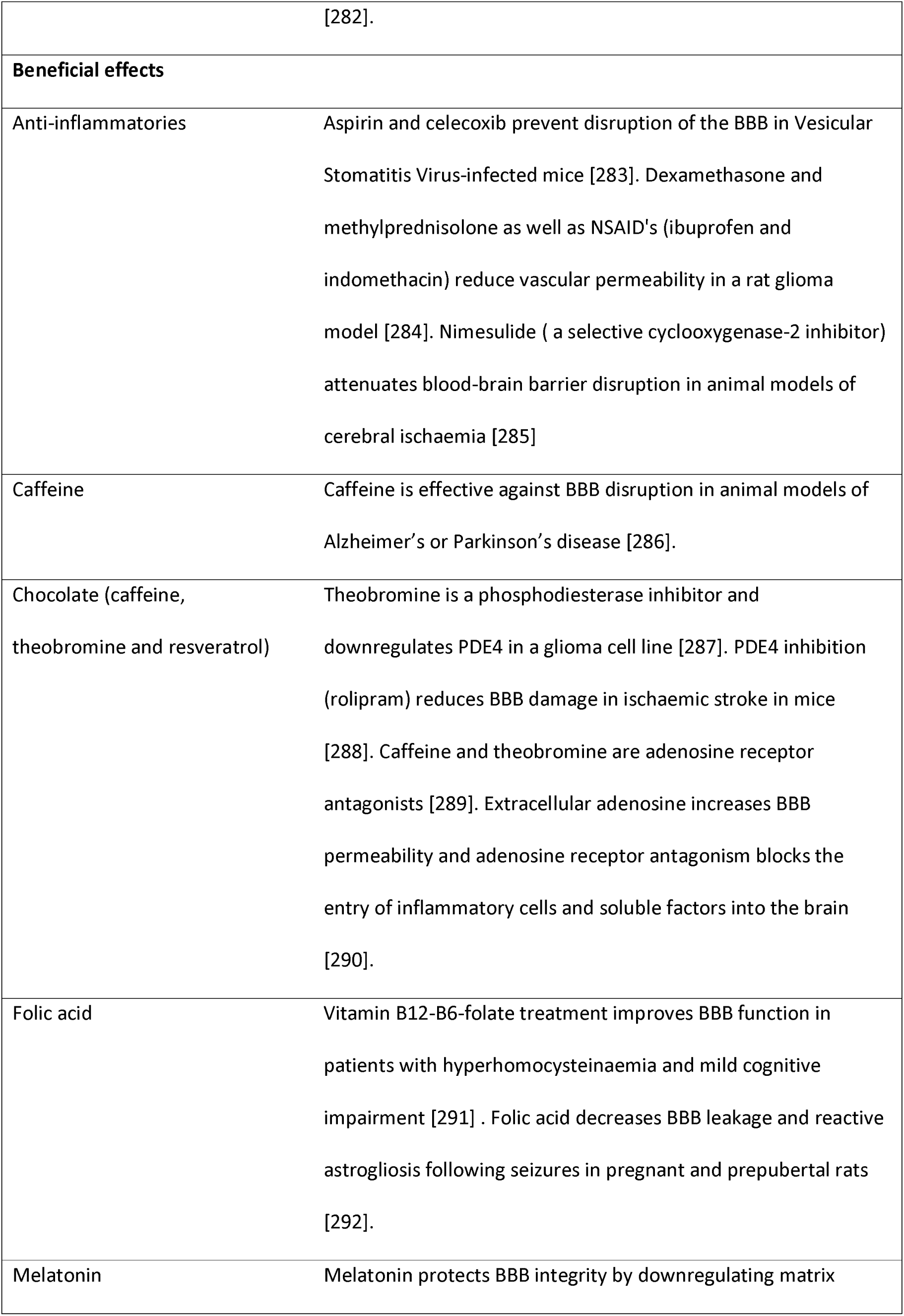
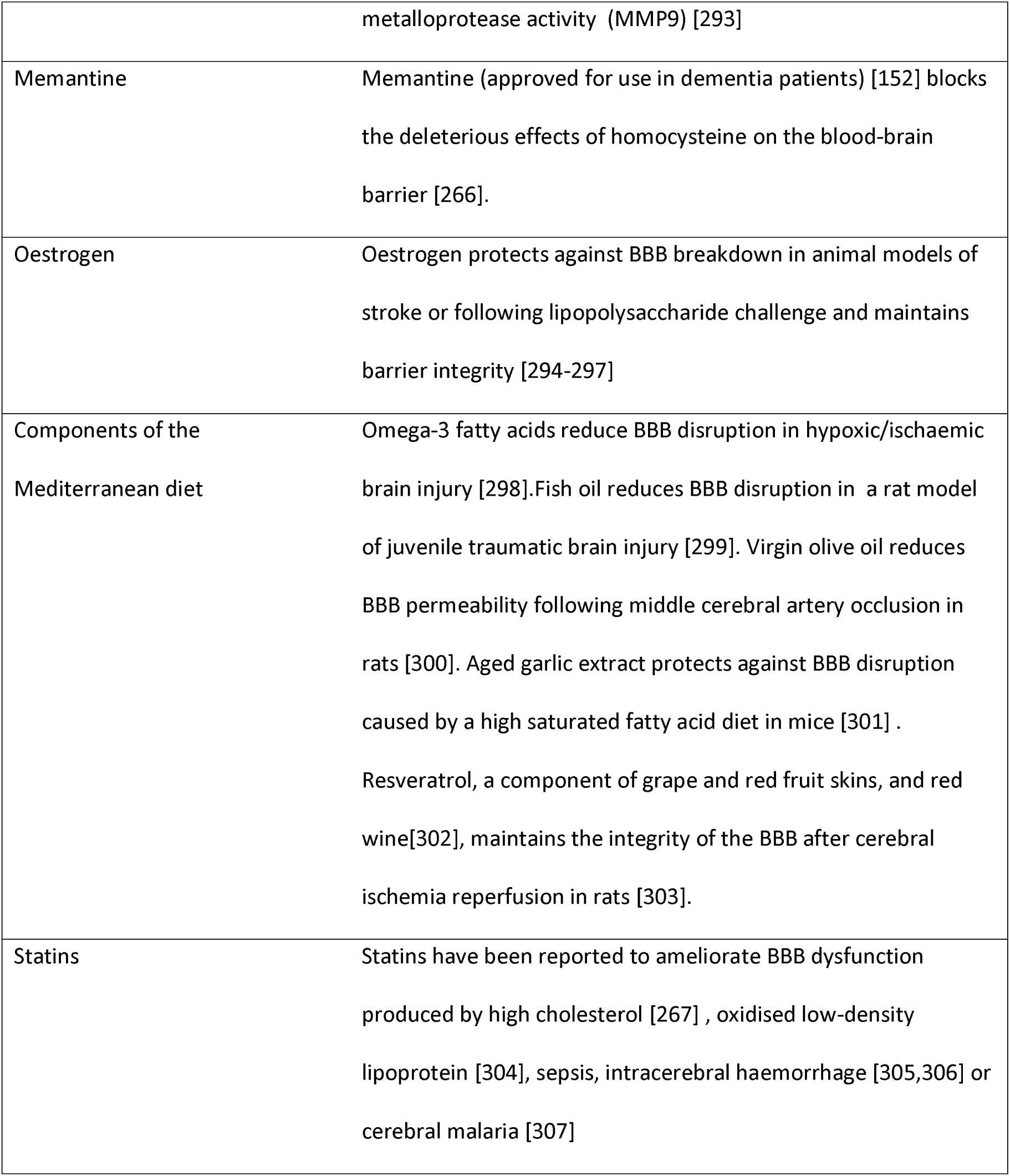
The effects of Alzheimer’s disease environmental risk factors and beneficial agents on blood brain barrier function.

### Diverse pathogen sensors and defenders relating to bacteria, viruses, parasites and fungi are upregulated in the AD brain, blood or CSF

We have evolved numerous pathogen detectors whose activation leads to stimulation of the immune system and to the production of defensive mechanisms, including inflammation and free radical attack. Multiple pattern recognition receptors including Toll-like receptors, C-type lectin receptors and nucleotide-binding oligomerization domain-like receptors (NOD-like) sense motifs in bacterial, viral, fungal and parasite proteins or other compounds or respond to foreign bacterial or viral DNA or RNA in cellular locations where host DNA or RNA should not exist [156–159].

Infection also activates inflammasomes, which trigger the maturation of proinflammatory cytokines, activating innate immune defences [160].

In addition to this, a large number of defensins and other antimicrobial peptides exist, targeting bacteria, fungi, parasites and viruses [161]. Beta-amyloid is one such [3].

EIF2AK2 (eukaryotic translation initiation factor 2 alpha kinase 2) better known as pkr, is activated by viral double stranded RNA and to bacterial RNA. This phosphorylates eif2alpha, leading to the arrest of the protein translation that is needed for viral replication. Pkr stimulation also leads to the production of interferon and to activation of the inflammasome [162–165]. Other viral RNA-sensors include RIG-I (coded by retinoic acid-inducible gene 1= DDX58), MDA5 (Melanoma Differentiation-Associated protein ; coded by IFIH1) and LGP2 (coded by DEXH-box helicase 58= DHX58) [92].

Indoleamine 2,3-dioxygenase 1 (IDO1) diverts tryptophan metabolism to N-formyl-kynurenine, (away from serotonin production). IDO1 upregulation is an important defence mechanism against pathogenic bacteria, many of which rely on host tryptophan. It is involved in antimicrobial defence and immune regulation, and its effects are not restricted to bacteria This IDO1 response is also deleterious to other pathogens and parasites, including T.Gondii, and to a number of viruses, including herpes simplex virus and other herpes viruses [166]. Kynurenine and kynurenic acid produced by IDO1 activation, are ligands for the aryl hydrocarbon receptor (AHR), which plays an important role in antimicrobial defence and immune regulation [167].

The function of these players with respect to the main pathogens studied above is reviewed in Supplementary Table 2, which also reports expression data in the Alzheimer’s disease brain, blood or CSF. Data derived from this table are illustrated in Figs 7 (viral) and 8 (bacteria, fungi and parasites).

**Figure 7:**
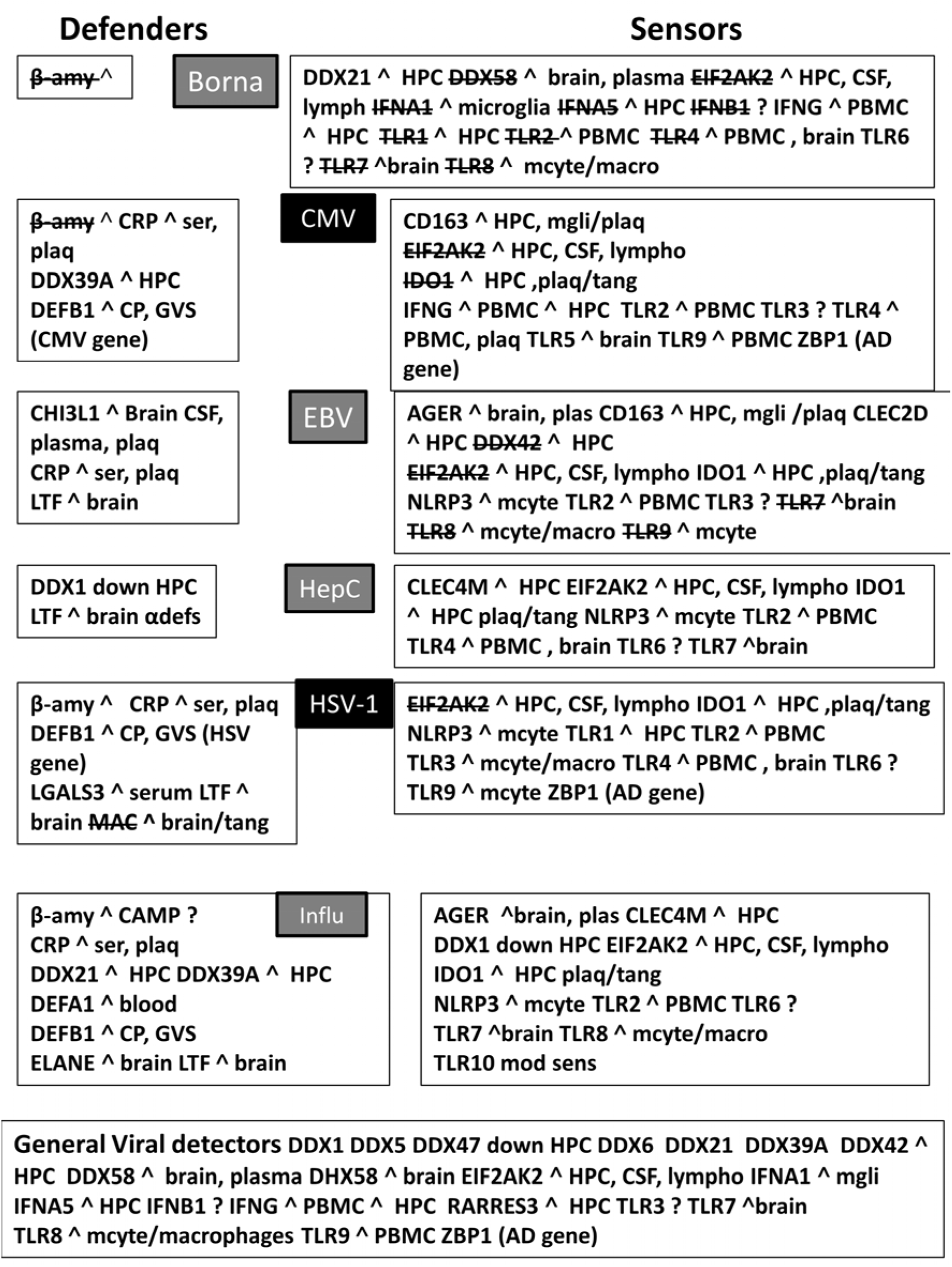
Fig 7 and 8. Viral (Fig 7) and fungal or bacterial (Fig 8) defenders and sensors and their expression (^ = upregulated; down = downregulated) in the brain, blood or cerebrospinal fluid of Alzheimer’s disease patients. CP = choroid plexus; CSF= cerebrospinal fluid; GVS= granulovacuolar degeneration; HPC = hippocampus; lympho = lymphocytes; macro = macrophages; mcyt=monocytes; mgli = microglia; PBMC = peripheral blood mononuclear cells; Plaq = amyloid plaques; Ser = serum; tang = tangles; αdefs or βdefs= unspecified alpha or beta defensins: AGER= advanced glycosylation end product-specific receptor (also known as RAGE); APCS= amyloid P component, serum ; CAMP = cathelicidin antimicrobial peptide (LL-37); Calpro= Calprotectin (S100A8/S100A9 dimer);CHI3L1 = chitinase 3 like 1 (aka YKL-40); C-type lectin = CLEC’s; CRP = C-reactive protein; DEAD box proteins = DDX’s; Defensins = DEFA’s, DEFB’s: EIF2AK2 = eukaryotic translation initiation factor 2 alpha kinase 2 (pkr); ELANE = elastase, neutrophil expressed ; IAPP = islet amyloid polypeptide (Amylin); IDO1= indoleamine 2,3-dioxygenase 1; Interferons = IFNA1, IFNA5, IFNB1, IFNG; LCN2 = lipocalin 2; LGALS3 = lectin, galactoside binding soluble 3; LTF = lactotransferrin; MAC = membrane attack complex (complement components C5b-C9); MRC1 = mannose receptor, C type 1; NAIP = NLR family, apoptosis inhibitory protein ; NLRP1 and 3 = NLR family pyrin domain containing 1 and 3; NOD1 and NOD2 = nucleotide binding oligomerization domain containing (1 and 2) ; RARRES2 and 3 = retinoic acid receptor responder (2 and 3) : S100’s= S100 calcium binding protein; Toll-like receptors = TLR1 to 10; ZBP1 Z-DNA binding protein 1. Gene = gene related to the respective pathogen in association studies or with Alzheimer’s disease (Gene AD). mod sens = modified senstitivity; The strikethrough’s (e.g. TLR1) represent a pathogen’s ability to inhibit or overcome the combative effects of the defensive or sensor protein. ? = unknown Borna = Borna virus; CMV = human cytomegalovirus; EBV = Epstein-Barr virus; HepC = Hepatitis C; HSV-1= Herpes simplex; Influ= Influenza A virus; Borrel= *Borrelia burgdorferi*; C.Alb= *Candida albicans*; C.Neo = *Cryptococcus neoformans*; C. Pneu = *Chlamydia pneumoniae*; H.Pyl= *Helicobacter pylori*; P.Ging = *Porphyrymonas gingivalis*; T.Gon = *Toxoplasma Gondii* Those shaded in black are those most implicated in Alzheimer’s disease (Table 1)

**Figure 8:**
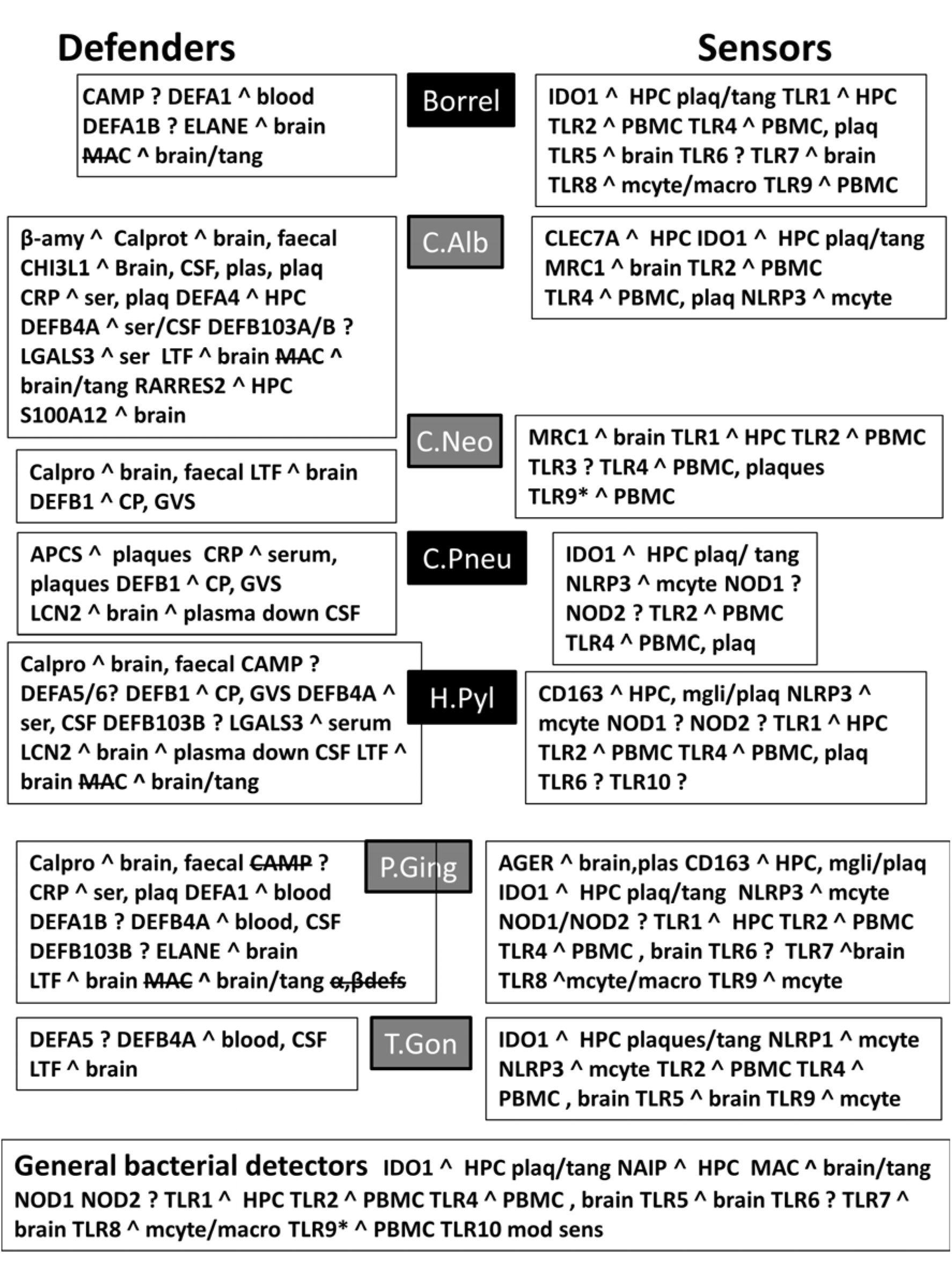
Fig 7 and 8. Viral (Fig 7) and fungal or bacterial (Fig 8) defenders and sensors and their expression (^ = upregulated; down = downregulated) in the brain, blood or cerebrospinal fluid of Alzheimer’s disease patients. CP = choroid plexus; CSF= cerebrospinal fluid; GVS= granulovacuolar degeneration; HPC = hippocampus; lympho = lymphocytes; macro = macrophages; mcyt=monocytes; mgli = microglia; PBMC = peripheral blood mononuclear cells; Plaq = amyloid plaques; Ser = serum; tang = tangles; αdefs or βdefs= unspecified alpha or beta defensins: AGER= advanced glycosylation end product-specific receptor (also known as RAGE); APCS= amyloid P component, serum ; CAMP = cathelicidin antimicrobial peptide (LL-37); Calpro= Calprotectin (S100A8/S100A9 dimer);CHI3L1 = chitinase 3 like 1 (aka YKL-40); C-type lectin = CLEC’s; CRP = C-reactive protein; DEAD box proteins = DDX’s; Defensins = DEFA’s, DEFB’s: EIF2AK2 = eukaryotic translation initiation factor 2 alpha kinase 2 (pkr); ELANE = elastase, neutrophil expressed ; IAPP = islet amyloid polypeptide (Amylin); IDO1= indoleamine 2,3-dioxygenase 1; Interferons = IFNA1, IFNA5, IFNB1, IFNG; LCN2 = lipocalin 2; LGALS3 = lectin, galactoside binding soluble 3; LTF = lactotransferrin; MAC = membrane attack complex (complement components C5b-C9); MRC1 = mannose receptor, C type 1; NAIP = NLR family, apoptosis inhibitory protein ; NLRP1 and 3 = NLR family pyrin domain containing 1 and 3; NOD1 and NOD2 = nucleotide binding oligomerization domain containing (1 and 2) ; RARRES2 and 3 = retinoic acid receptor responder (2 and 3) : S100’s= S100 calcium binding protein; Toll-like receptors = TLR1 to 10; ZBP1 Z-DNA binding protein 1. Gene = gene related to the respective pathogen in association studies or with Alzheimer’s disease (Gene AD). mod sens = modified senstitivity; The strikethrough’s (e.g. TLR1) represent a pathogen’s ability to inhibit or overcome the combative effects of the defensive or sensor protein. ? = unknown Borna = Borna virus; CMV = human cytomegalovirus; EBV = Epstein-Barr virus; HepC = Hepatitis C; HSV-1= Herpes simplex; Influ= Influenza A virus; Borrel= *Borrelia burgdorferi*; C.Alb= *Candida albicans*; C.Neo = *Cryptococcus neoformans*; C. Pneu = *Chlamydia pneumoniae*; H.Pyl= *Helicobacter pylori*; P.Ging = *Porphyrymonas gingivalis*; T.Gon = *Toxoplasma Gondii* Those shaded in black are those most implicated in Alzheimer’s disease (Table 1)

These figures show that sensors and defenders relating to multiple pathogens are upregulated in the AD brain, blood or CSF. These involve reactions to many different classes (bacteria, viruses, fungi and parasites) and there appears to be no discrimination, or focus on any particular type. This would concord with the multiple and diverse pathogen species that have been detected in the AD brain (Table 1) and with the relationship between the AD genes or the hippocampal transcriptome with multiple pathogen species.

## Caveats

This analysis is based on overlapping gene symbols rather than on specific polymorphisms. There is thus no indication of the physiological weight or importance of any gene/pathogen interaction, some of which will be more important than others. Pathogen effects may also be strain-dependent, and the size of the interactomes also varies widely. Within any large interactome there will be deleterious, neutral and beneficial effects. While HSV-1 infection causes beta-amyloid deposition and neurodegeneration [168], in its latent form, the virus can have neuroprotective effects. For example the viral latency transcript inhibits apoptosis and promotes neurite sprouting in neuroblastoma cells [169], protects neuronal C1300 and Neuro2A cells from granzyme B-induced apoptosis and CD8 T-Cell killing [170] and also protects trigeminal neurones from apoptosis [171]. The Bornavirus is capable of promoting hippocampal degeneration in Man [172]. In rats Borna virus infection decreases choline acetyltransferase activity in the cerebral cortex, horizontal diagonal band of Broca, hippocampus and amygdala [173] a situation similar to that observed in Alzheimer’s disease [174] but the inflammation and microglial activation it produces can also reduce beta-amyloid immunoreactivity in the brain parenchyma of Tg2576 mutant beta-amyloid mice [93]. Chronic, adult acquired T. Gondii infection causes neurologic and behavioural abnormalities secondary to inflammation and neuronal loss, in a strain-dependent manner [175]. T.Gondii infection in BALB/C mice induces neuroinflamation and learning and memory deficits. It also potentiates the toxic effects of low doses of intracerebrally administered beta-amyloid[176], but chronic infection can also increase beta-amyloid phagocytosis and clearance by recruited monocytes [177].

Dementia or neurodegeneration, in the absence of amyloid plaques is, by current clinical definition, not considered as Alzheimer’s disease, but as already noted, there is no inherent biological reason for this [178,179]. Such divergent effects might also be relevant to findings relating to the presence of amyloid plaques in the absence of dementia, as observed in the Nun study [180,181] or to diagnosed Alzheimer’s disease in the absence of beta-amyloid. A recent report showed that ~15% of patients clinically diagnosed with AD do not have amyloid deposits as indexed by positron emission tomography [182]. While some amyloid-negative patients could be re-diagnosed (~50%), the clinical follow-up using other criteria in other amyloid-negative patients continued to support the definition of Alzheimer’s disease.

There are also many inter-pathogen interactions relevant to this relatively small sample of the potential microbiome. For example HSV-1 infection activates replication of the Epstein-Barr virus, [183]. Gingipains or other proteases secreted by *P. Gingivalis* degrade multiple complement components [184] as well as alpha-and beta defensins [185],immunoglobulins, IgG1 and IgG3 [186] and interleukin-12, preventing its ability to stimulate interferon production [187]. Such effects enable the pathogen to counteract immune defence and would also impinge on the viability of many other pathogens.

HIV-1 is immunosuppressant and has been associated with many opportunistic pathogens including tuberculosis, toxoplasmosis, cytomegalovirus encephalitis and Cryptococcal brain invasion [188,189]. The human cytomegalovirus is also immunosuppressant via an ability to target MHC class I molecules for degradation [190]and to inhibit MHC class II antigen presentation [191]. Parasites, which maintain a long-term, if unwelcome presence in the host have also developed immunosuppressant and anti-defensive strategies[192,193]. In addition, the success of most pathogens depends upon their ability to subvert the defensive armoury of the host in some way.

The AD genes affect human processes relevant to the disease itself, but given that they are also part of pathogen interactomes, polymorphisms therein are also likely to affect pathogen life cycles or the ability of pathogens to promote diverse effects within the host. Apart from **APOE4** there are no studies relating to the effects of the AD gene variants on pathogens or their effects.

For these and many other reasons, it is perhaps unwise to rank the pathogens by order of importance in relation to their enrichment or p value in any of the data described above. Suffice it to say that diverse pathogens have been detected in the AD brain and all of the bioinformatics data presented above, whether related to genes, transcriptomes, plaques or tangles implicate multiple species of pathogens across viral, bacterial, fungal and protozoan classes.

While there are statistical limitations to this type of analysis, correction for false discovery followed by the Bonferroni correction has been conservatively applied. The relationship of AD to pathogens is supported by experimental observation (Table 1) and by the antimicrobial effects of beta-amyloid. This study also relies on multiple and diverse *in silico* bioinformatic analyses linking AD GWAS genes, plaques and tangles as well as the hippocampal transcriptome to multiple pathogen interactomes, and the upregulated AD hippocampal genes to multiple infection datasets from diverse pathogen species. Polymicrobial involvement is also supported by the diversity of bacterial, viral and fungal sensors and defenders that are upregulated in the AD brain, blood or CSF. Each comparison relates to single pathogens but given the diversity of pathogens detected in AD such effects are likely to be cumulative.

## Discussion

Multiple and diverse pathogens (bacteria, viruses, fungi and spirochetes) have been detected in the AD brain and many cause neurodegeneration, increase beta-amyloid deposition and tau phosphorylation or are killed/incapacitated by beta-amyloid, an antimicrobial peptide that is part of the innate immune defence system. Representatives of these pathogens target multiple AD GWAS genes, and their interactomes are enriched in genes related to the AD hippocampal transcriptome and to the proteins found in AD plaques and tangles. The upregulated genes of the AD hippocampal transcriptome also correspond to those upregulated by multiple species of viral, bacterial, fungal and protozoan pathogens or by interferon gamma and Toll-like receptor ligands.

The AD genes are preferentially localised in the bone marrow and other immunocompetent tissues, and in exosomes that are hijacked by pathogens for intercellular spread. They are also localised in the lateral ventricle and the hippocampus which abuts this area, a prime site of pathogen invasion via the choroid plexus and the blood/csf barrier.

The AD genes are enriched in global GWAS datasets relating to pathogen diversity, suggesting that some have been selected for pathogen resistance rather that susceptibility. This is supported by the old age of AD patients, indicating survival from the many infections that contribute to mortality in the younger population. **APOE4** variants protect against malaria and hepatitis C, and immune/inflammatory gain of function applies to **APOE4, CR1, TREM2** and presenilin variants, supporting this contention. Logically, any gene variant increasing the production of the antimicrobial peptide beta-amyloid in response to pathogens might also be considered as beneficial in these evolutionary terms. Apart from APOE4, there is however little data examining the effects of AD gene variants on pathogen life cycles or that relate specifically to pathogen responses.

Many AD genes are also localised in the blood brain barrier. This should provide an effective shield against many infections but it is disrupted by multiple environmental risk factors implicated in Alzheimer’s disease and protected by several factors reported to be beneficial in relation to Alzheimer’s disease, including NSAIDs, statins, oestrogen, memantine, melatonin, and components of the Mediterranean diet.

The relationship between pathogens and Alzheimer’s disease has a long history coupled with a degree of scepticism, perhaps related to an inability to fulfil Koch’s postulate. For example, the same pathogen is not always found in all AD brains, or in different laboratories. Laboratory confirmation in animal models may be impossible for certain pathogens, for example the Epstein-Barr or hepatitis C virus, that do not infect rodents. Nevertheless, the diversity of pathogens able to promote neurodegeneration, beta-amyloid deposition or to mimic the effects observed in the hippocampal AD transcriptome suggests that many candidates, alone or severally, could be involved in the pathogenesis of AD. A polymicrobial involvement seems likely given the multiple species detected in the AD brain. Evidently, this could be assessed by microbiome studies in the periphery or in postmortem brains.

Recent work suggests that the production of the antimicrobial/antiviral peptide beta-amyloid is an expected consequence of infection in general [2,3]. In the context of the amyloid hypothesis [194], this places pathogens upstream of the production of this toxic peptide, and logically as causal, both in terms of beta-amyloid production and in relation to Alzheimer’s disease.

Two separate case reports have shown remission from dementia or mis-diagnosed Alzheimer’s disease in patients subsequently diagnosed with and treated for *Cryptococcus neoformans* infection [50,51].

In a Greek study, *H. Pylori*-infected AD patients receiving the triple eradication regime (omeprazole, clarithromycin and amoxicillin) showed improved cognitive and functional status parameters where bacterial eradication was successful [195].*H. Pylori* eradication in AD patients with peptic ulcer was also associated with a decreased risk of AD progression in a Taiwanese study [196].

Taking all of the above into consideration the combined data suggest that polymicrobial brain invasion, enabled by environmentally-induced blood-brain barrier defects may be responsible for Alzheimer’s disease. This could essentially be mediated via activation of a hyper-efficient inflammatory network, including the call-up of beta-amyloid that, as a consequence, causes massive neuronal destruction in a tissue incapable of regeneration. The role of the innate immune system and the inflammatory response in neurotoxicity has recently been reviewed, and innate surveillance mediated cell death has been suggested as a plausible common pathogenic pathway responsible for many neurodegenerative diseases, including AD [197].

It is therefore not unreasonable to suggest that antibiotic, antifungal and antiviral agents, possibly in combination, tailored to the individual, might be able to halt, delay or perhaps even provide remission in patients with Alzheimer’s disease.

## Acknowledgements

Thanks are due to the many authors who have sent reprints and supplementary datasets that made this work possible. Thanks also to David Eby from the Broad Institute of MIT and Harvard for his help with the structure of the Molecular Signatures database.

The author reports no funding and no conflict of interest.

---

**Supplementary table 1:** Definitions of the Alzheimer’s disease susceptibility genes studied. While many other functions are recognised, for example relating to beta-amyloid, cholesterol, lipid and glucose metabolism or diabetes, *inter alia* [1-4], the properties isolated in this table focus specifically on immune and pathogen-related effects. The relationship between AD genes, the immune system and inflammation has also previously emphasised [5] and in a recent study from the Alzheimer’s Disease Neuroimaging Initiative, another subset of Alzheimer’s disease genes showed genetic overlap between Alzheimer’s disease and immune-mediated diseases [6].

| Gene<br>Symbol | Name | Immune or pathogen related properties |
| --- | --- | --- |
| ABCA7 | ATP-binding cassette, sub-family A<br>(ABC1), member 7 | Plays a prominent role in phagocytosis by macrophages (demonstrated with *Staphylococcus aureus*). This is an important line of general host defence against pathogens [7]. Overexpression of ABCA7 in HeLa cells resulted increases intracellular and cell surface ceramide and intracellular phosphatidylserine levels [8]. Ceramide reactivates the herpes simplex virus from latency [9] and is also incorporated into *C.Pneumoniae* inclusions [10]. APOA1 and APOE are substrates for ABCA7, and in cultured |
|  |  | HEK-293 cells, plasma membrane-situated ABCA7 increases the efflux of phosphatidylcholine and sphingomyelin efflux to APOA1 and APOE, with no effect on cholesterol efflux[11] . Sphingomyelin is enriched in extracellular herpes simplex viral membranes [12] . It is a receptor for the Helicobacter toxin VacA [13] and is also incorporated into inclusion bodies in *C.Pneumoniae* infected cells [14].<br>Phosphatidylcholine plays an important role in the fusion of herpes simplex glycoproteins B and H with the host cell lipid membrane, a process used in viral entry [15]. Phosphatidylcholine is also able to trigger capsular enlargement in *C.Neoformans* infection [16].<br>Cholesterol efflux to lipid-laden APOE, but not to lipid free APOE, is increased by ABCA7 expression in HEK-293 cells [17]. |
| ACE | angiotensin I converting enzyme | Modifies the C termini of peptides for |
|  |  | presentation by major histocompatibility complex class I molecules, which increases the efficiency of antigen-specific CD8+ T cell priming [18]. |
| ADAMTS20 | ADAM metalloproteinase with thrombospondin type 1 motif, 20 | Cleaves the chondroitin sulfate proteoglycan, versican[19] which interacts with myeloid and lymphoid cells promoting their adhesion and the production of inflammatory cytokines: Inflammatory agents, such as double-stranded viral RNA mimetics, stimulate stromal cells, smooth muscle cells and fibroblasts, to produce fibrillar extracellular matrices enriched in versican and hyaluronan that promote the adhesion of leukocytes [20] |
| AP2A2 | adaptor-related protein complex 2, alpha 2 subunit | Induces the renewal and maintenance of hematopoietic stem cells [21]. Required for binding of human immunodeficiency virus type 1 Nef and cooperative assembly of a CD4-Nef- |
|  |  | AP-2 complex [22]. |
| APOC1 | apolipoprotein C-I | APOC1 binds to lipopolysaccharide (LPS), an outer-membrane component of gram-negative bacteria and is involved in the presentation of LPS to macrophages. This improves the inflammatory response , thus protecting against infection [23]. **APOC1** is a component of high density lipoprotein: Herpes simplex is present in all lipoprotein blood fractions in blood (VLDL, LDL and HDL) and the lipid component of these lipoproteins binds to viral glycoprotein B [24] |
| APOE | apolipoprotein E | APOE4 favours cerebral access of HSV-1 in mice [25] and enhances C.pneumoniae adherence to host cells [26] and HIV-1 cell entry in vitro [27], but is protective against chronic hepatitis C virus infection [28]. The allele relates to increased viral load in HHV-6 infected epilepsy patients [29]. Hepatitis B pathology has a more benign course in ApoE2-E4 carriers |
|  |  | [30]. |
| ATXN7L1 | ataxin 7-like 1 | None found |
| BCAM | basal cell adhesion molecule<br>(Lutheran blood group) | Adhesion molecule involved in red blood cell adhesion to the vascular endothelium [31] Also plays a role in abnormal red blood cell adhesion in sickle cell disease (c.f. malaria) [32]. Acts as a receptor for Escherichia coli cytotoxic necrotizing factor 1, a toxin found in E.coli strains causing meningitis [33] |
| BCL3 | B-cell CLL/lymphoma 3 | BCL3 is essential for the development, survival and activity of adaptive immune cells. BCL3-deficient mice are more susceptible to bacterial and parasitic infection [34]. |
| BIN1 | bridging integrator 1 | BIN1 negatively controls the expression of indoleamine 2,3-dioxygenase IDO1 in cancer cells [35]. IDO1 activation diverts tryptophan metabolism to N-formyl-kynurenine, (away from serotonin production) .IDO1 upregulation is an important |
|  |  | defence mechanism against pathogenic bacteria, many of which rely on host tryptophan. It is involved in antimicrobial defence and immune regulation, and its effects are not restricted to bacteria This IDO1 response is also deleterious to other pathogens and parasites, including T.Gondii, and to a number of viruses, including herpes simplex virus and other herpes viruses [166]. Kynurenine and kynurenic acid produced by IDO1 activation, are ligands for the aryl hydrocarbon receptor (AHR), which plays an important role in antimicrobial defence and immune regulation [167].<br>A BIN1 isoform is required for macrophage phagocytosis, a key mechanism in the destruction of many pathogens [36] |
| CASS4 | Cas scaffolding protein family member 4 | One of a member of scaffold proteins are regulated by and mediating cell attachment, growth factor, and chemokine signalling [37] |
| CD2AP | CD2-associated protein | CD2AP and other endocytosis-associated proteins play a role in enteropathogenic *Escherichia coli* pedestal formation [38]: Also required for late endosomal trafficking of the *H. pylori* VacA toxin [39]. Clathrin and related proteins including CD2AP are involved in the recruitment of proteins that promote actin polymerization at the interface of T cells and antigen presenting cells [40]. Decreased CD2AP expression enhances the production of type I interferons in human plasmacytoid dendritic cells which secrete type I interferons in response to microbial stimuli [41] |
| CD33 | CD33 molecule | A member of the sialic acid binding Immunoglobulin g-like lectin (SIGLEC) family. CD33-related SIGLEC's regulate adaptive immune responses and are also important as macrophage pattern recognition receptors for sialylated pathogens, including enveloped viruses [42]. |
|  |  | CD33 binds to alpha 2-3- or alpha2-6-linked sialic acids (N-acetyl neuranimic acid) [43]. These residues bind to the influenza virus and the reovirus [44] and these particular sialic acids are expressed on the surface envelope glycoproteins (B, D and H) of the herpes simplex virion, and are required for viral entry into cells [45]. N-acetyl neuranimic acid is expressed by *C.Neoformans* and is involved in fungal adhesion to macrophages [46] and is also a component of the cell wall of *B.Burgdorferi* [47] while *Helicobacter pylori* adhesins also bind to this particular form of sialic acid [48,49] as does *P.Gingivalis*[50]. CD33 binds to sialic acid acquired by *P.aeruginosa* and to the HIV-1 gp120 protein [51] . |
| CDON | cell adhesion associated, oncogene regulated | A gene associated with the acquisition of *Staphylococcus aureus* bacteraemia [52] |
| CEACAM16 | carcinoembryonic antigen-related | The CEACAM family are docking sites |
|  | cell adhesion molecule 16 | for pathogenic bacteria [53] but this particular protein has not been characterised in relation to this effect |
| CELF1 | CUGBP, Elav-like family member 1 | A downstream effector of interferon beta signalling in macrophages [54]. |
| CLU | clusterin | Inhibits the membrane attack complex, composed of complement components C5 to C9. This is deposited on the bacterial surface forming channels that cause bacterial lysis [55,56] . |
| CNTNAP2 | contactin associated protein-like 2 | None found |
| CR1 | complement component (3b/4b) receptor 1 (Knops blood group) | Many pathogens are recognised by the complement system and coated with complement components C1q, C3b and iC3b. This “opsonisation” prepares the microbe for phagocytosis via binding of the complement components to complement receptors, including CR1 [57]. Receptor for the malaria pathogen *Plasmodium falciparum* [58], *Legionella pneumophila* [59], *Mycobacterium tuberculosis*[60] and *Cryptococcus neoformans* |
| CUGBP2<br>(changed to<br>CELF2) | CUGBP, Elav-like family member 2 | CUGBP2 silences the expression of cyclo-oxygenase 2 (PTGS2), thus regulating inflammatory processes [61]<br>CUGBP2 is regulated in response to T-cell signalling and increased CELF2 expression drives a network of activation-induced alternative splicing events in Jurkat cells [62]. |
| DISC1 | disrupted in schizophrenia 1 | DISC1 has many functions relevant to the psychiatric diseases in which it is implicated, among which is control of the intracellular traffic of mRNAs, neurotransmitter receptors, vesicles and mitochondria along the microtubule network [63,64] . Although DISC1 has not been related to any particular virus or pathogen, the microtubule network provides a set of railway tracks used by many viruses during their life cycles [65-67]. Such traffic is also important in the regulation of the immunological synapse and in the building of functional phagosomes [68] |
| ECHDC3 | enoyl CoA hydratase domain | Expressed in whole blood cells and |
|  | containing 3 | platelets, but no functional data available [69,70] .One of several genes downregulated by Trypanosoma Cruzi in mouse macrophages [71] |
| EPHA1 | EPH receptor A1 | Suppresses T cell activation and Th2 cytokine expression, while preventing activation-induced cell death in the lung [72]. Upregulated in dendritic antigen-presenting cells in response to the human papillomavirus E7 peptide [73] . Mice infected with M. tuberculosis displayed higher expression of EPHA1 and EPHA2 in monocytes as well as ephrinA1[74] |
| EXOC3L2 | exocyst complex component 3-like<br>2 | None found |
| FAM113B<br>(now<br>C5orf64) | chromosome 5 open reading frame<br>64 | This locus is considered non-coding by other groups due to a lack of experimental support for the protein, but NCBI annotates the protein because it meets minimal RefSeq quality criteria for representation. The coding status remains uncertain. [19 Nov |
|  |  | 2014](Refseq) |
| FANCD2OS | FANCD2 opposite strand | No functional publications |
| FERMT2 | fermitin family member 2 | None found |
| FLJ37543<br>(now<br>C5orf64) | chromosome 5 open reading frame<br>64 | None found |
| FRMD4A | FERM domain containing 4A | None found |
| GAB2 | GRB2-associated binding protein 2 | An adaptor protein involved in multiple receptor tyrosine kinase signalling pathways: phosphorylated by stimulation with growth factors-, cytokines-, Immunoglobulin Fc- and antigen receptors [75] Gab2 knockout mice show reduced inflammatory cytokine levels in, and are relatively protected against Mycobacterium tuberculosis infection[76]. |
| GRIN3B | glutamate receptor, ionotropic, N-methyl-D-aspartate 3B | None found |
| HLA-DRB1 | major histocompatibility complex, class II, DR beta 1 | Bind to pathogen antigens and present them to T-cells [77]. |
| HLA-DRB5 | major histocompatibility complex, class II, DR beta 5 |  |
| HMHA1 | histocompatibility (minor) HA-1 | When HA-1 peptide was added to mixtures of plasmacytoid DC dendritic cells and T cells, bystander suppression of the response to a colocalized recall Epstein-barr viral antigen occurred primarily via indolamine-2,3-dioxygenase (IDO1) production.<br>Bystander suppression is a process whereby Antigen-specific (adaptive) T regulatory cells inhibit the T effector cell response both to specific antigen and to a colocalized third-party antigen [78]: minor histocompatibility antigens refer to immunogenic peptides which, when complexed with MHC, can generate an immune response after recognition by specific T-cells. The peptides are derived from polymorphic intracellular proteins, which are cleaved by normal pathways of antigen processing (Definition from Uniprot). |
| HS3ST1 | heparan sulfate (glucosamine) 3-O- | Heparan sulfate biosynthetic enzymes |
|  | sulfotransferase 1 | are key components in generating a myriad of distinct heparan sulfate fine structures that carry out multiple biologic activities. The enzyme encoded by this gene is a member of the heparan sulfate biosynthetic enzyme family. It possesses both heparan sulfate glucosaminyl 3-O-sulfotransferase activity, anticoagulant heparan sulfate conversion activity, and is a rate limiting enzyme for synthesis of anticoagulant heparan. This enzyme is an intraluminal Golgi resident protein. [provided by RefSeq, Jul 2008]. Heparan sulphates act as attachment sites for many viruses [79,80] |
| IGH | immunoglobulin heavy locus | Forms the heavy chain of multiple antibodies [77]. |
| INPP5D | inositol polyphosphate-5-phosphatase, 145kDa | Phosphatidylinositol (PtdIns) phosphatase that specifically hydrolyses the 5-phosphate of phosphatidylinositol-3,4,5-trisphosphate (PtdIns(3,4,5)P3) to |
|  |  | produce PtdIns(3,4)P<sub>2</sub>, thereby negatively regulating the PI3K (phosphoinositide 3-kinase) pathways. Acts as a negative regulator of B-cell antigen receptor signalling. Mediates signalling from the FC-gamma-RIIB receptor (FCGR2B), playing a central role in terminating signal transduction from activating immune/hematopoietic cell receptor systems. Acts as a negative regulator of myeloid cell proliferation/survival and chemotaxis, mast cell degranulation, immune cells homeostasis, integrin alpha-IIb/beta-3 signalling in platelets and JNK signalling in B-cells. Regulates proliferation of osteoclast precursors, macrophage programming, phagocytosis and activation and is required for endotoxin tolerance. Involved in the control of cell-cell junctions, CD32a signalling in neutrophils and modulation of EGF-induced phospholipase C activity. Key regulator of neutrophil migration, by |
|  |  | governing the formation of the leading edge and polarization required for chemotaxis. Modulates FCGR3/CD16-mediated cytotoxicity in NK cells. Mediates the activin/TGF-beta-induced apoptosis through its Smad-dependent expression. May also hydrolyse PtdIns(1,3,4,5)P4, and could thus affect the levels of the higher inositol polyphosphates like InsP6.2 (Definition from Uniprot) |
| LUZP2 | leucine zipper protein 2 | None found |
| MEF2C | myocyte enhancer factor 2C | MEF2C orchestrates early B-cell development [81] and is also involved in the activation induced cell death of macrophages after priming with Salmonella typhimurium, type 5 adenovirus or Interferon-gamma [82]. Also a risk gene for periodontitis [83], a known risk factor for Alzheimer's disease [84] |
| MMP12 | matrix metalloproteinase 12<br>(macrophage elastase) | Degrades elastin, a matrikine derived from extracellular matrix proteins: These are implicated in inflammation, |
|  |  | immune responses, organ development, wound repair, angiogenesis, atherosclerosis, tumor progression and metastasis due to their ability to alter cellular migration, chemotaxis, and mitogenesis.[85] . Aging and various inflammatory diseases such as atherosclerosis, abdominal aortic aneurysms, chronic obstructive pulmonary diseases, cancer and type 2 diabetes are characterized by the destruction of elastin fibres [86] |
| MMP3 | matrix metalloproteinase 3<br>(stromelysin 1, progelatinase) | PolyI:C treatment (viral DNA mimic) increases the expression levels of Mmp3 mRNA and protein in astrocytes, but not microglia [87]. |
| MPZL1 | myelin protein zero-like 1 | Present in CD133(+) precursors (CD133= hematopoietic precursor antigen) and endothelial cells, and mainly in mesenchymal and committed myelomonocytic progenitor cells, and in erythroid precursor cell lines [88]. |
| MS4A3 | membrane-spanning 4-domains,<br>subfamily A, member 3<br>(hematopoietic cell-specific) | Modulates cell cycle progression in<br>hematopoietic cells [89] |
| MS4A4A | membrane-spanning 4-domains,<br>subfamily A, member 4A | Localised in Hematopoietic cells [89]:<br>Expressed in lung mast cells Silencing<br>MS4A4 promotes mast cell<br>proliferation and migration. Mast cells<br>express Toll receptors and play an<br>important role in pathogen recognition<br>and in acquired immunity against<br>parasitic infections [90,91]. |
| MS4A4E | membrane-spanning 4-domains,<br>subfamily A, member 4E | None found |
| MS4A6A | membrane-spanning 4-domains,<br>subfamily A, member 6A | Localised in Lymphoid tissues, Kidney<br>Colon and Wilm's tumor cells [89] |
| MSRA | methionine sulfoxide reductase A<br>Catalyses two reactions (from<br>KEGG)<br>(1) peptide-L-methionine +<br>thioredoxin disulfide + H<sub>2</sub>O =<br>peptide-L-methionine (S)-S-oxide +<br>thioredoxin; | Could have an important function as a<br>repair enzyme for proteins that have<br>been inactivated by oxidation.<br>Catalyzes the reversible oxidation-<br>reduction of methionine sulfoxide in<br>proteins to methionine (From Uniprot). |
|  | (2) L-methionine + thioredoxin<br>disulfide + H<sub>2</sub>O = L-methionine<br>(S)-S-oxide + thioredoxin |  |
| MTHFD1L | methylenetetrahydrofolate<br>dehydrogenase (NADP+ dependent)<br>1-like Catalyses the reaction<br>(KEGG)<br>ATP + formate + tetrahydrofolate =<br>ADP + phosphate + 10-<br>formyltetrahydrofolate | The protein encoded by this gene is<br>involved in the synthesis of<br>tetrahydrofolate (THF) in the<br>mitochondrion. THF is important in the<br>de novo synthesis of purines and<br>thymidylate and in the regeneration of<br>methionine from homocysteine<br>(Refseq) |
| NDUFAF6 | NADH dehydrogenase (ubiquinone)<br>complex I, assembly factor 6 | None found |
| NME8 | NME/NM23 family member 8 | The NME8 locus has been associated<br>in a genome-wide study with the<br>bacterial disease periodontitis [92] also<br>a known risk factor for Alzheimer's<br>disease [84]. |
| PAX2 | paired box 2 | PAX2 negatively regulates beta<br>defensin-1, an antimicrobial peptide<br>implicated in the resistance of<br>epithelial surfaces to microbial<br>colonization [93]. |
| PCDH11X | protocadherin 11 X-linked | None found |
| PCNX1 | pecanex homolog (Drosophila) | None found |
| PICALM | phosphatidylinositol binding<br>clathrin assembly protein | Involved in clathrin-mediated endocytosis, a process used by many viruses to gain entry to the cell [94]<br>(AP2A2 and BIN1 are also involved in this process)see KEGG pathway (red text genes)<br>http://www.genome.jp/kegg-bin/show_pathway?hsa04144+274+161 |
| POLN | polymerase (DNA directed) nu | POLN can perform translesion synthesis past thymine glycol, a common endogenous and radiation-induced product of reactive oxygen species damage to DNA. Thymine glycol blocks DNA synthesis by most DNA polymerases, but POLN was particularly adept at efficient and accurate translesion synthesis past a 5S-thymine glycol [95]. |
| PPP1R37 | protein phosphatase 1, regulatory subunit 37 | No publications |
| PPP1R3B | protein phosphatase 1, regulatory subunit 3B | None found |
| PTK2B | protein tyrosine kinase 2 beta | Involved in Toll-like receptor signalling (pathogen recognition receptors)(TLR2, TLR4) in macrophages [96]. Involved in the natural killer cell cytotoxic pathway [97] and in the microglial production of nitric oxide produced by lipopolysaccharide and interferon gamma [98] |
| PVR | poliovirus receptor | Mediates entry of the poliovirus and binds to NECTIN1 ( a receptor for HSV-1 and 2) [99] and NECTIN3 ( a receptor for HSV-1) [100] [101,102] |
| PVRL2 | poliovirus receptor-related 2 (herpesvirus entry mediator B) | Entry receptor for HSV-1 [103]. |
| RELN | reelin | Reelin plays a prominent role in the brain but also in the intestine where the reeler mutation down-regulates genes related to the immune response, inflammation, and tumor development [104] . Reelin deposits in the |
|  |  | hippocampus are a conserved neuropathological feature of aging, and such deposits are accelerated in adult wild-type mice prenatally exposed to a viral-like infection [105]. |
| RFC3 | replication factor C (activator 1) 3, 38kDa | The elongation of primed DNA templates by DNA polymerase delta and DNA polymerase epsilon requires the accessory proteins proliferating cell nuclear antigen (PCNA) and replication factor C (RFC).RFC3 is one of 5 subunits of this complex (Refseq). Host nuclear DNA processing factors are also recruited to viral genomes, RFC3 is one of many recruited to the HSV-1 viral genome [106] |
| RIN3 | Ras and Rab interactor 3 | RIN 3 inhibits mast cell migration toward stem cell factor, which recruits mast cells to sites of infection or injury, where they release pro-inflammatory substances [107]. |
| SASH1 | SAM and SH3 domain containing 1 | Scaffold molecule involved in Toll receptor (TLR4) signalling, a receptor involved in the recognition of bacterial |
|  |  | lipopolysaccharides[108]. |
| SCIMP | SLP adaptor and CSK interacting membrane protein | SCIMP is expressed in B cells and other antigen-presenting cells and is involved in major histocompatibility complex class II signalling [109]. |
| SLC24A4 | solute carrier family 24 (sodium/potassium/calcium exchanger), member 4 | None found |
| SLC4A1AP | solute carrier family 4 (anion exchanger), member 1, adaptor protein | None found |
| SORL1 | sortilin-related receptor, L(DLR class) A repeats containing | None found |
| SPPL2A | signal peptide peptidase like 2A | SPPL2A is a protease that cleaves CD74, the invariant chain of the MHCII complex, and an important chaperone regulating antigen presentation for the immune response. [110]. |
| SQSTM1 | sequestosome 1 | Autophagy can either promote or restrict viral replication. SQSTM1 is an autophagy receptor involved in the life |
|  |  | cycles of the Chikungunya virus[111], Coxsackievirus[112], Dengue virus [113], the encephalomyocarditis virus [114], enterovirus 71[115], hepatitis B [116] , HIV-1[117], Herpes simplex (HSV-1) [118], Kaposi's sarcoma virus [119], measles [120], Varicella zoster [121] and the West Nile virus [122] |
| STK24 | serine/threonine kinase 24 | Important regulator of neutrophil degranulation which results in the releases of proteases and other cytotoxic agents, including matrix metalloproteinases and myeloperoxidase These granule contents are antimicrobial, but can also cause tissue damage [123] |
| TOMM40 | translocase of outer mitochondrial membrane 40 homolog (yeast) | The influenza viral protein PB1-F2 translocates into mitochondria via TOMM40 channels and impairs innate immunity [124]. TOMM40 is required for replication of the African swine fever virus [125] |
| TREM2 | triggering receptor expressed on | A receptor for bacterial lipopolysaccharide that acts as a |
|  | myeloid cells 2 | phagocytic receptor for bacteria. It also inhibits the production of inflammatory cytokines induced by Toll like receptors [126-128]. |
| TREML2 | triggering receptor expressed on myeloid cells-like 2 | TREML2 (Triggering receptor expressed on myeloid cells (TREM)-like transcript 2) is expressed on T cells and regulates interleukin-2 and interferon-gamma production [129]. |
| TRIP4 | thyroid hormone receptor interactor 4: | None found: This protein is localized in the nucleus and contains an E1A-type zinc finger domain, which mediates interaction with transcriptional coactivators and ligand-bound nuclear receptors, such as thyroid hormone receptor and retinoid X receptor alpha, but not glucocorticoid receptor (Refseq). |
| TTLL7 | tubulin tyrosine ligase-like family, member 7 | None found |
| ZCWPW1 | zinc finger, CW type with PWWP domain 1 | None found |
| ZNF224 | zinc finger protein 224 | Wilms tumor 1 (WT1) recruits ZNF224 to the interferon regulatory factor 8 (IRF8) promoter [130]<br>The IRF family proteins bind to the IFN-stimulated response element (ISRE) and regulate expression of genes stimulated by type I IFNs, namely IFN-alpha and IFN-beta. IRF family proteins also control expression of IFN-alpha and IFN-beta-regulated genes that are induced by viral infection. [provided by RefSeq, Jul 2008] |

**Supplementary table 2:** A survey of the roles of diverse microbial sensors and defensive proteins. Their expression levels in the Alzheimer’s disease brain, blood, cerebrospinal fluid or other defined cells etc. are also reviewed.

| Gene | Function | Alzheimer's disease |
| --- | --- | --- |
| AGER advanced | Recognizes advanced | Increases in protein levels |
| glycosylation end product-specific receptor (more commonly known as RAGE) | glycosylation end products, members of the S100 protein family, beta-amyloid and amyloid fibrils, HMGB1, and β-integrin macrophage 1 antigen (Mac-1) [1]. Receptor for S100B , S100A4, 6,11,12,13, S100P [2], Expressed on endothelial cells macrophages, neutrophils, dendritic cells, T cells, B cells, alveolar type II cells and alveolar epithelial cells[3]. AGER(-/-) mice were relatively protected from influenza virus induced mortality showing improved viral clearance , enhanced cellular T cell response and activation of neutrophils [4]. AGER activation enhances the ability of neutrophils to eradicate bacteria (E.Coli) in vitro and in vivo via activation of NADPH | and in the percentage of AGER expressing microglia in the Alzheimer's disease brain linked with disease severity[10].Plasma protein levels increased in Alzheimer's disease [11] but decreased levels of a soluble isoform [12,13] |
|  | oxidase[5].<br>Involved in the adhesion of *Helicobacter pylori* to gastric epithelial cells [6]. P. gingivalis infection enhances AGER expression in Murine aortic endothelial cells [7]. Bacteria produce, metabolize and accumulate AGEs .*Escherichia coli* cells secrete AGEs [8]. Influenza A viral pneumonia is associated with enhanced AGER expression on endothelium and de novo expression on bronchial epithelium in mice[4]. The Epstein Barr viral protein LMP1 binds to the AGER promoter [9] |  |
| βAmyloid | Antimicrobial peptide with broad spectrum activity against bacterial (Enterococci, E.Coli, streptococci, staphylococci, pseudomonas, listeria) and | Key component of amyloid plaques |
|  | fungal (*Candida albicans*) species {Soscia, Kirby, et al. 2010 1912 /id} . It also has antiviral effects against Herpes simplex (HSV-1) {Bourgade, Le Page, et al. 2016 10595 /id}{Bourgade, Garneau, et al. 2015 10596 /id} and the influenza virus {White, Kandel, et al. 2014 8740 /id}. Borna virus infection induced microglial activation can reduce plaque formation in Tg2576 APP mutant transgenic mice [14] |  |
| APCS amyloid P component, serum (commonly known as SAP) | Binds to several bacterial lipopolysaccharides (S. pyrogens and rough strains of *E. coli*) preventing complement activation[15]. Increased levels of APCS in the atherotic plaques of *C. Pneumoniae* infected mice fed an atherogenic diet [16] . Binds avidly to *C. Albicans* | Protein levels elevated in the AD brain and associated with plaques, but low levels in plaques were seen in individuals with AD pathology without dementia [18] . |
|  | when amyloid is formed in fungal cell walls [17] |  |
| CAMP cathelicidin antimicrobial peptide (LL-37) | In addition to its antibacterial, antifungal, and antiviral activities, the encoded protein functions in cell chemotaxis, immune mediator induction, and inflammatory response regulation. [provided by RefSeq, Sep 2014].<br>[19]Antiviral versus influenza<br>Kills P.Gingivalis [20] but degraded by a P.gingivalis secreted protease (gingipain) [21]. DEFB1 and CAMP (cathelicidin/LL-37) kill H.pylori [22]. Borrelia burgdorferi is killed by human polymorphonuclear leukocyte granule components ( elastase ELANE, CAMP, bactericidal/permeability- | None found |
|  | increasing protein (BPI), and human neutrophil peptide-1)[23]. |  |
| CD163 CD163 molecule | Functions as an acute phase-regulated receptor involved in the clearance and endocytosis of hemoglobin/haptoglobin complexes by macrophages, and may thereby protect tissues from free hemoglobin-mediated oxidative damage. This protein may also function as an innate immune sensor for bacteria and inducer of local inflammation. [provided by RefSeq, Aug 2011]<br>Upregulated in the gastric mucosa of H. pylori infected children [24]. Upregulated by P.Gingivalis in periodontal ligament cells [25]. Kupffer cell/macrophage activation indicated by increased | Upregulated in the AD hippocampus [29].<br>Parenchymal microglia were immunoreactive for CD163 in all of 31 AD cases often associated with amyloid plaques [30] |
|  | CD163 is found in the livers of hepatitis C infected patients [26]. The cytomegalovirus encoded IL10 chemokine mimic upregulates CD163 in macrophages [27]. serum levels of soluble CD163 in Epstein-Barr virus positive children positively correlate with EBV-DNA copies [28] |  |
| CHI3L1 chitinase 3 like 1<br>(aka YKL-40) | Chitinases catalyze the hydrolysis of chitin, which is an abundant glycopolymer found in insect exoskeletons and fungal cell walls. The protein lacks chitinase activity and is secreted by activated macrophages, chondrocytes, neutrophils and synovial cells. The protein is thought to play a role in the process of inflammation and tissue remodeling. [provided by | CSF levels of CHI3L1 are associated with Alzheimer's disease [33-35]. Plasma levels are also increased and the protein is found in astrocytes near a subset of amyloid plaques (immunohistochemistry) [36] |
|  | RefSeq, Sep 2009]<br>CHI3L1 is induced by fungal infection (*Candida albicans*) and induces the antimicrobial peptides beta-defensin 3 and cathelicidin (CAMP) [31]. In transgenic mice expressing the Epstein-Barr virus LMP1 protein CHI3L1 is induced in the epidermis and is secreted and autoantibodies to CHI3L1 are generated [32] |  |
| CLEC2B C-type lectin domain family 2 member B | CLEC-2 is a HIV-1 attachment factor and platelets capture and transfer infectious HIV-1 via DC-SIGN and CLEC-2 [37]. Expression induced by infection of Akata cells [38] | Upregulated in the AD hippocampus [29] |
| CLEC2D C-type lectin domain family 2 member D | Expression upregulated by respiratory syncytial virus (RSV) infection, in the BEAS-2B respiratory epithelial cell line and | Upregulated in the AD hippocampus [29] |
|  | primary human bronchial epithelial cells [39].<br>Expression is induced in B cells and inflamed tonsils following viral infection ( Epstein-Barr virus or HIV infection) and in inflamed tonsils [40]. |  |
| CLEC4M C-type lectin domain family 4 member M (L-SIGN) | .....recognizes numerous evolutionarily divergent pathogens ranging from parasites to viruses, with a large impact on public health.....[provided by RefSeq, Feb 2009] CD209 (DC-SIGN) and CLEC4M (L-SIGN) are endocytic receptors for influenza A virus entry and infection, and for the Hepatitis C virus , HIV-1, Sindbis virus, and act as cofactors for cellular entry by Ebola virus.<br>CLEC4M also a receptor for Mycobacterium tuberculosis, | Upregulated in the AD hippocampus [29] |
|  | Schistosomes and<br>Leishmania infant [41-46] |  |
| CLEC7A C-type lectin<br>domain family 7 member A<br>(Dectin 1) | Functions as a pattern-recognition receptor that<br>recognizes a variety of beta-1,3-linked and beta-1,6-linked glucans from fungi<br>and plants, and in this way<br>plays a role in innate immune<br>response..... [provided<br>by RefSeq, Jul<br>2008].Activated by<br>C.albicans, and<br>Mycobacterium bovis [47] | Upregulated in the AD<br>hippocampus [29] |
| CRP C-reactive protein,<br>pentraxin-related | Involved in several host<br>defence related functions<br>based on its ability to<br>recognize foreign pathogens<br>and damaged cells of the host<br>and to initiate their<br>elimination by interacting<br>with humoral and cellular<br>effector systems in the blood.<br>Consequently, the level of | High serum levels associated<br>with AD (dependent on<br>methodology) [54], but levels<br>of CRP in a mild and<br>moderate dementia subgroup<br>were significantly lower than<br>that in the control group [55].<br>A recently developed high-<br>sensitivity (Hs) test reported<br>high serum levels of Hs-CRP |
|  | this protein in plasma increases greatly during acute phase response to tissue injury, infection, or other inflammatory stimuli.<br>[provided by RefSeq, Sep 2009]. Chlamydial lipopolysaccharide serum levels in coronary syndrome correlate with CRP levels [48]. High CRP levels observed in H.Pylori and C.Pneumoniae infection [49].<br>High antibody response to multiple pathogens (cytomegalovirus, herpes simplex virus-1, Hepatitis A virus, Helicobacter pylori and Chlamydia pneumoniae) associated with CRP in atherosclerosis patients [50].<br>Antibodies to P. gingivalis associate with high levels of SAA and high concentrations of CRP in periodontitis | in AD patients[56]. CRP staining of the hippocampal CA1/2 region correlates with Aβ staining in the AD brain [57]. |
|  | patients[51]. Serum CRP elevated in fungal esophagitis or enterocolitis due to C.albicans [52]. High serum levels of CRP found in numerous bacterial or viral infections : (Dengue virus , Cytomegalovirus , Epstein Barr virus, Parvovirus B19 , HSV-1 and -2 and Influenza A and B: [53] |  |
| DDX1 DEAD/H-box helicase 1 | DDX1, DDX21, and DHX36 helicases form a complex with the adaptor molecule TRIF to sense double stranded viral RNA, including Influenza and Poly-IC in dendritic cells [58].Binds to hepatitis C biotinylated RNA [59] | Down regulated in the AD hippocampus [29] |
| DDX18 DEAD-box helicase 18 | Few publications | Upregulated in the AD hippocampus [29] |
| DDX21 DEAD-box helicase 21 | DDX1, DDX21, and DHX36 helicases form a complex | Upregulated in the AD hippocampus [29] |
|  | with the adaptor molecule TRIF to sense double stranded viral RNA a in dendritic cells [58]. DDX21 inhibits replication of the influenza virus [60].Interacts with a Borna virus protein [61] |  |
| DDX27 DEAD-box helicase<br>27 | ? | Upregulated in the AD hippocampus [29] |
| DDX39A DEAD-box helicase 39A | Needed for the expression of Kaposi sarcoma-associated herpesvirus ORF57 [62]. The UL69 gene product of the human cytomegalovirus belongs to a family of regulatory proteins conserved among all herpesviruses and binds to DDX39A [63]. Mx proteins exert their antiviral activity against the influenza virus by interfering with the function of the RNA helicases DDX39B and DDX39A [64] . | Upregulated in the AD hippocampus [29] |
| DDX42 DEAD-box helicase<br>42 | The expression of N-terminal DDX42 binds to the NS4A protein of the Japanese encephalitis virus and DDX42 is able to overcome antagonism of interferon responses by the virus [65]. Also a potential target of an Epstein-Barr viral microRNA [66] . The Japanese encephalitis virus encodes for interferon antagonist proteins, one of which , NS4A, binds to DDX42 [65] | Upregulated in the AD hippocampus [29] |
| DDX47 DEAD-box helicase<br>47 | Interacts with the E1E4 protein of human papillomavirus type 16 [67] | Down regulated in the AD hippocampus [29] |
| DDX5 DEAD-box helicase 5 | This gene encodes a DEAD box protein, which is a RNA-dependent ATPase, and also a proliferation-associated nuclear antigen, specifically reacting with the simian virus 40 tumor antigen..... | Down regulated in the AD hippocampus [29] |
|  | [provided by RefSeq, Feb 2016] DDX3,5 and 6 play a role in hepatitis C viral replication [68]. DDX5 interacts with the SARS coronavirus [69] |  |
| DDX58 DEXD/H-box helicase 58 (commonly known as RIG-1) | DEAD box proteins, characterized by the conserved motif Asp-Glu-Ala-Asp (DEAD), are putative RNA helicases which are implicated in a number of cellular processes involving RNA binding and alteration of RNA secondary structure. This gene encodes a protein containing RNA helicase-DEAD box protein motifs and a caspase recruitment domain (CARD). It is involved in viral double-stranded (ds) RNA recognition and the regulation of immune response. [provided by | Expression increased in the temporal cortex and plasma of mild cognitive impairment patients with pathologic evidence of senile plaques and neurofibrillary tangles [71] |
|  | RefSeq, Jul 2008] The DDX58 -activating 5' triphosphate group is removed post-transcriptionally by a viral function and modified<br>DDX58 does not bind the RNAs of Hantaan virus, Crimean-Congo hemorrhagic fever virus or the Borna disease virus [70]. |  |
| DDX6 DEAD-box helicase 6 | The protein is an RNA helicase found in P-bodies and stress granules, and functions in translation suppression and mRNA degradation. It is required for microRNA-induced gene silencing. Multiple alternatively spliced variants, encoding the same protein, have been identified.<br>[provided by RefSeq, Mar 2012]. It also controls gene expression in RNA viruses | Upregulated in the AD hippocampus [29] |
|  | [72].<br>DDX3,5 and 6 play a role in hepatitis C viral replication<br>[68]. Binds to a Dengue virus RNA [73] |  |
| DEFA1 defensin alpha 1 | Found in the microbicidal granules of neutrophils and likely plays a role in phagocyte-mediated host defence. (from Refseq)<br>Defends against *S. aureus* , *E. coli* and *E. aerogenes* [74]<br>anthrax toxin, *C. Difficile* toxin B, diphtheria toxin, and *Pseudomonas* exotoxin A [75-77] and also inhibit the adenovirus, BK polyoma virus and HIV-1 [78-80].<br>Binds to *P. Gingivalis* [81]. | Defensins alpha1 and 2 (now coded only by DEFA1) are upregulated in Alzheimer's disease blood cells [84];<br>DEFA1/DEFA1B , DEFA3 and DEFB4A increased in sera and CSF of AD patients [85] |
|  | Antiviral versus Influenza A [82]. Alpha-defensin transcription activated by the hepatitis C core protein (specific gene symbol not possible) [83] |  |
| DEFA1B defensin alpha 1B | The protein encoded by this gene, defensin, alpha 1, is found in the microbicidal granules of neutrophils and likely plays a role in phagocyte-mediated host defense. Several alpha defensin genes are clustered on chromosome 8. This gene differs from defensin, alpha 3 by only one amino acid (from Refseq). Binds to P.Gingivalis [81]. Release induced by H.Pylori [86]. Borrelia burgdorferi is killed by human polymorphonuclear leukocyte granule components ( elastase | DEFA1/DEFA1B , DEFA3 and DEFB4A increased in sera and CSF of AD patients [85] |
|  | ELANE, CAMP, bactericidal/permeability-increasing protein (BPI), and human neutrophil peptide-1(DEFA1))[23] |  |
| DEFA3 defensin alpha 3 | Found in the microbicidal granules of neutrophils and likely plays a role in phagocyte-mediated host defense. (from Refseq)<br>Defends (relatively weakly) against S. aureus , E. coli and E. aerogenes [74] | DEFA1/DEFA3/ DEFB4A elevated in the serum and cerebrospinal fluid of AD patients [84] |
| DEFA4 defensin alpha 4 | Found in neutrophils; it exhibits corticostatic activity and inhibits corticotropin stimulated corticosterone production. [provided by RefSeq, Oct 2014]. Potent killer of Escherichia coli, Streptococcus faecalis, and Candida albicans [87]. | Upregulated in the hippocampus [29] |
| DEFA5 defensin alpha 5 | The protein encoded by this gene, defensin, alpha 5, is highly expressed in the | ? |
| | secretory granules of Paneth cells of the ileum. [provided by RefSeq, Oct 2014] Kills *H.Pylori* [88]. Unmethylated CpG motifs in *Toxoplasma gondii* DNA induce TLR9- and IFN-β-dependent expression of DEFA5 in intestinal epithelial cells.[89] | |
| DEFA6 defensin alpha 6 | The protein encoded by this gene, defensin, alpha 6, is highly expressed in the secretory granules of Paneth cells of the small intestine, and likely plays a role in host defense of human bowel. [provided by RefSeq, Oct 2014] Kills *H.Pylori* [88] | ? |
| DEFB1 defensin beta 1 | A gene associated with HSV-1 and cytomegalovirus seropositivity in children with acute lymphoblastic leukaemia [90], as well as with *H.Pylori* or chlamydial infections [91,92], also | Upregulated in the Alzheimer's disease choroid plexus and in granulovacuolar degeneration structures [95] |
|  | endowed with antimicrobial activity against *C.Neoformans* and other pathogens [93].DEFB1 and CAMP (cathelicidin/LL-37) kill *H.pylori* [22]. Protects mice from influenza pathogenesis with a mechanism other than inhibition of viral replication. plasmacytoid dendritic cells and monocytes increased production of DEFB1 peptide and mRNA as early as 2 h following infection of purified cells and peripheral blood mononuclear cells with influenza , HSV-1, and Sendai virus[94]. |  |
| DEFB103A defensin beta 103A | An antibiotic peptide which is induced by bacteria and interferon gamma, and which displays antimicrobial activity against *S. aureus*, *S. pyogenes*, *P. aeruginosa*, *E.* | ? |
|  | coli, and C. albicans.<br>[provided by RefSeq, Oct 2014] . |  |
| DEFB103B defensin beta 103B | This gene encodes defensin, beta 103, which has broad spectrum antimicrobial activity and may play an important role in innate epithelial defense. [provided by RefSeq, Oct 2014] Kills H.Pylori [96]. Binds to P.Gingivalis [81]. Antiviral versus Influenza A [97] | ? |
| DEFB4A defensin beta 4A | This gene encodes defensin, beta 4, an antibiotic peptide which is locally regulated by inflammation. [provided by RefSeq, Jul 2008]. Has potent antimicrobial activity against Gram-negative bacteria and Candida, but not Gram-positive Staphylococcus aureus [98]. kills H.Pylori [99] Also involved in defence against | DEFA1/DEFA3/ DEFB4A elevated in the serum and cerebrospinal fluid of AD patients [84] |
|  | Varicella zoster, human respiratory syncytial virus, HIV-1 and the Human papillomavirus [100-103].<br>Binds to P.Gingivalis [81].<br>slow-replicating type II and III T.Gonidii induce high levels of DEFB4A gene expression in human intestinal epithelial cells [104] |  |
| Defensins | A large number of antimicrobial peptides (almost 2000 animal derived peptides, 112 from Homo Sapiens) target bacteria, parasites, fungi or viruses [105]. Beta-amyloid can be considered as one such [106]<br>. |  |
| DHX58 DEXH-box helicase<br>58 | Detects double stranded viral RNA and activates antiviral responses [107,108]. | Upregulated in total brain and frontal lobe of AD patients [109] |
| EIF2AK2 eukaryotic translation initiation factor 2 alpha kinase 2 (commonly known as PKR) | Several stimuli including TNF and other cytokines, double stranded viral RNA or bacterial ligands acting via Toll receptors activate EIF2AK2 resulting in the | Upregulated in the AD hippocampus [29] and CSF [121] and activated in AD lymphocytes [122] |
|  | inhibition of protein synthesis necessary for viral replication. Activation also results in the production of interferons alpha and beta [110]. Activated by lipopolysaccharide or bacterial RNA or by the mycotoxin deoxynivalenol, shiga toxin, and ricin [111-115]. Activated by HCMV, but the virus possesses proteins able to antagonise EIF2AK2 [116]. Activated by HSV-1 which is also able to evade EIF2AK2 activation [117] and by hepatitis C and influenza viruses [118].<br>Epstein-Barr virus-encoded small RNAs bind the protein PKR and inhibit its activation [119]. Not activated by the Borna virus, suggesting an evasive strategy to abolish antiviral activities [120] |  |
| ELANE elastase, neutrophil expressed | Following activation, this protease hydrolyzes proteins within specialized neutrophil lysosomes, called azurophil granules, as well as proteins of the extracellular matrix. The enzyme may play a role in degenerative and inflammatory diseases through proteolysis of collagen-IV and elastin. This protein also degrades the outer membrane protein A (OmpA) of *E. coli* as well as the virulence factors of such bacteria as *Shigella*, *Salmonella* and *Yersinia*. [provided by RefSeq, Jan 2016] . Kills *Borrelia burgdorferi* [23]. *H. pylori* extract-activated human neutrophils result in endothelial cell detachment from human umbilical vein endothelial cells monolayers | Increased expression in the vessel wall of leptomeningeal vessels in AD . Arterial elastin degradation was observed from Braak stage III onward and correlated with Braak tau pathology [126]. In the brain parenchyma elastase immunoreactivity is restricted to neurons and is markedly elevated in a proportion of neurofibrillary tangle-bearing neurons [127]. |
|  | which can be blocked by an elastase antibody. The bacterium also inhibits elastase [123]. Elevated serum levels in patients with influenza virus-associated encephalopathy [124].<br>Periodontain, a protease secreted by P.Gingivalis, inactivates the human serpin, alpha1-proteinase inhibitor, the primary endogenous regulator of human neutrophil elastase, which may be responsible for increased elastase activity in periodontitis [125] . |  |
| Gamma-secretase | Localised in dendritic cells that scout for invading pathogens. Cleaves receptors for many pathogens including those for adenoviruses, C.Neoformans, cytomegalovirus, Epstein-Barr virus, Hendra virus, hepatitis C, HHV-6, HIV-1, HSV-1, influenza, rhinovirus, measles, Nipah virus, Papilloma virus, P.Gingivalis, rabies, S.Aureus and streptococci, Vaccinia and other pox viruses [128]. |  |
| IAPP islet amyloid | Commonly found in | Accumulates intraneuronally |
| polypeptide (Amylin) | pancreatic islets of patients suffering diabetes mellitus type II, or harboring an insulinoma. Studies suggest that this protein, like the related beta-amyloid (Abeta) associated with Alzheimer's disease, can induce apoptotic cell-death in particular cultured cells, an effect that may be relevant to the development of type II diabetes. This protein also exhibits a bactericidal, antimicrobial activity. [provided by RefSeq, Sep 2014]. Inhibits the growth of Staphylococcus aureus and Escherichia coli [129] | in brains of Alzheimer's disease patients, particularly in those with type-2 diabetes [130]. See review for common links between bacteria, diabetes and Alzheimer's disease [131] |
| IDO1 indoleamine 2,3-dioxygenase 1 | Catalyses the production of N-formylkynurenine from tryptophan. Expression is stimulated by interferon gamma and other inflammatory cytokines. | IDO1 expression is increased in the AD hippocampus and is associated with amyloid plaques and neurofibrillary tangles. Quinolinic acid immunoreactivity is localised |
|  | This diverts tryptophan metabolism away from serotonin production, towards kynurenines and can lead to overproduction of the kynurenic acid and quinolinic acid, N-methyl-D-aspartate receptor antagonist and agonist respectively. The subsequent depletion of tryptophan is deleterious to many microbes that depend upon this metabolite [132].<br>Diversion to the kynurenine pathway also produces metabolites activating the aryl hydrocarbon receptor which also plays a role in antimicrobial defence and immune activation. This pathway is relevant to anti-bacterial and antiviral effects[133]. Involved in *C.albicans* defence [134] and in the response to B. | in microglial and astrocytic cells around amyloid plaques and in the vicinity of neurofibrillary tangles [145-147]. |
|  | Burgdorferi [135].Restricts C.Pneumoniae replication in dendritic cells [136].Induced by HSV-1 [137], Influenza and hepatitis C infection [138,139] . Induced by C.Albicans at sites of infection and in dendritic cells and effector neutrophils [140]. IDO1 activation restricts HCMV replication, but the virus is able to counteract this block [141]. Expression increased by a DPG3 strain of P.Gingivalis [142] . Activated by T.Gondii infection in the mouse spleen [143] . Induced by the Epstein-Barr virus in human macrophages [144] |  |
| IFNA1 interferon, alpha 1 | The protein encoded by this gene is produced by macrophages and has antiviral activity. This gene is intronless and the encoded | The NK cell activity induced by either interferon-alpha (IFN-alpha) or interleukin-2 (IL-2) in DAT was also significantly lower than in |
|  | protein is secreted. [provided by RefSeq, Sep 2011]<br>Orthologs<br>a | the normal controls [148]<br>white matter microglia were intensely labeled for alpha-IFN [149] |
| IFNB1 Interferon beta 1 | The protein encoded by this gene belongs to the type I class of interferons, which are important for defense against viral infections. In addition, type I interferons are involved in cell differentiation and anti-tumor defenses. Following secretion in response to a pathogen, type I interferons bind a homologous receptor complex and induce transcription of genes such as those encoding inflammatory cytokines and chemokines.<br>Overactivation of type I interferon secretion is linked to autoimmune diseases.<br>Mice deficient for this gene display several phenotypes | Increased cytotoxic response by NK cells to IL-2 (mean increase +102%) and IFN-beta (mean increase +132%) in SDAT patients [151]. |
|  | including defects in B cell maturation and increased susceptibility to viral infection. [provided by RefSeq, Sep 2015]The Borna virus nucleoprotein inhibits type I IFN expression by interfering with the IRF7 pathway [150] . |  |
| IFNG Interferon Gamma | The active protein is a homodimer that binds to the interferon gamma receptor which triggers a cellular response to viral and microbial infections. Mutations in this gene are associated with an increased susceptibility to viral, bacterial and parasitic infections and to several autoimmune diseases. [provided by RefSeq, Dec 2015]. A P.Gingivalis protease, (gingipain) cleaves interleukin-12, reducing its | Increased spontaneous and IL-2-induced release of IFN-gamma and TNF-alpha from NK cells were found in DAT patients compared to healthy subjects. [154]: IFN-γ and TNF-α levels, in peripheral blood mononuclear cells, assessed in patients with AD in mild and severe stages, respectively, are higher than those observed in patients with moderate stage and MCI [155]. Increased IL2 and IFNG secretion from mononuclear cells observed |
|  | ability to stimulate IFNG production [152] .<br>Upregulated in the brains of Borna virus infected cats [153]. | in AD patients in the moderately severe stage of the disease [156] : IFNG levels increased in peripheral blood mononuclear cells [157]. No increase in plasma [158]<br>or CSF levels [159] : higher levels of IL-1beta (interleukin 1beta) ($P < .001$), IL-1beta to IL-1ra ratio ($P < .001$), tumor necrosis factor alpha ($P = .008$), IL-6 ($P = .04$), and interferon gamma ($P = .01$) in the non-afflicted offspring of patients with AD [160].<br>All participants with Apo $\epsilon 3/\epsilon 4$ or $\epsilon 4/\epsilon 4$ alleles showed a distinct biochemical profile characterized by low C-reactive protein and ApoE levels and by high cortisol, interleukin 13, apolipoprotein B, and gamma interferon |
|  |  | levels[161] . CSF interferon $\gamma$ was only detected in cytomegalovirus seropositive subjects and was significantly associated with neurofibrillary tangles [162] . Higher levels of IL-6 and IFN-γ were found more in the cultured T lymphocytes of the AD patients [163]. IFNA5 and IFNG upregulated in the AD hippocampus [29]. Infectious burden and IFNG levels associated with AD (HCMV, HSV-1, B. burgdorferi, C. pneumoniae and H. pylori) [164] |
| LCN2 lipocalin 2 | This gene encodes a protein that belongs to the lipocalin family. Members of this family transport small hydrophobic molecules such as lipids, steroid hormones and retinoids. The protein | Lcn2 levels are decreased in CSF of patients with mild cognitive impairment and AD and increased in brain regions associated with AD pathology in human postmortem brain tissue |
|  | encoded by this gene is a neutrophil gelatinase-associated lipocalin and plays a role in innate immunity by limiting bacterial growth as a result of sequestering iron-containing siderophores.<br>Mice lacking this gene are more susceptible to bacterial infection than wild type mice. [provided by RefSeq, Sep 2015] involved in host defence against *C.Pneumoniae* possibly by limiting the availability of iron to the pathogen [165].<br>Upregulated in the gastric mucosa of *H.Pylori* infected patients [166] | [167] .Plasma levels are increased in mild cognitive impairment [168]. |
| LGALS3 lectin, galactoside binding soluble 3 | The protein exhibits antimicrobial activity against bacteria and fungi..[provided by RefSeq, Oct 2014]<br>LGALS3 knockout mice are more susceptible to | Serum levels increased in AD [172] |
|  | C.Albicans infection [169].<br>Plays an important role in innate immunity to infection and colonization of H. pylori [170]. HSV-1 infection increases the carbohydrate binding activity and the secretion of cellular LGALS3 [171] |  |
| LTF lactotransferrin | Antimicrobial, antiviral, antifungal and antiparasitic activity has been found for this protein and its peptides. Alternatively spliced transcript variants encoding different isoforms have been found for this gene.<br>[provided by RefSeq, Sep 2014]. Kills T.Gondii and C.Albicans [173].<br>lactoferricin is generated by gastric pepsin cleavage of lactoferrin and kills albicans, C. tropicalis and C. neoformans[174].Neutralises | expression up-regulated in both neurons and glia in affected AD tissue [181] |
|  | HSV-1 and prevents replication [175]. Inhibits P.Gingivalis proteases [176]. Effective versus H.Pylori [177]. Inhibits influenza virus hemagglutination [178]. Antiviral versus hepatitis C [179]. Inhibits Epstein Barr virus infection [180] |  |
| MAC: Membrane attack complex: A complex composed of complement components C5b to C9 that attaches to bacteria, creating pores that kill by lysis [182]. | Activated by C.Albicans but secreted fungal proteases degrade C5 and can inhibit MAC formation [183,184]. Activated by P.Gingivalis which is also able to degrade C5 [185,186]. Kills H.Pylori in vitro but the pathogen evades MAC by binding to CD59, and inhibitor of MAC formation [187]. Attacks Borrelia burgdorferi , which retaliates via a protein (CspA) which binds C7 and C9 and blocks MAC assembly and membrane | The complement system is activated in the AD brain and MAC is abundantly present and associated with neurofibrillary tangles, in the neuronal cytoplasm, lipofuscin granules, lysosomes, dystrophic neurites within neuritic plaques, and neuropil threads [190-192] |
|  | insertion [188]. HSV-1 infected neuronal or skin cells activate complement and though initially resistant to MAC deposition the skin cells eventually succumb to MAC deposition. Neuronal Paju cells are more resistant but MAC is deposited on ~10% of these [189]. |  |
| MRC1 mannose receptor, C type 1 | The protein encoded by this gene is a type I membrane receptor that mediates the endocytosis of glycoproteins by macrophages. The protein has been shown to bind high-mannose structures on the surface of potentially pathogenic viruses, bacteria, and fungi so that they can be neutralized by phagocytic engulfment.[provided by RefSeq, Sep 2015].<br>Recognises C.Albicans [193]. Higher fungal burdens | mRNAs for TNF, AGI, MRC1 and CHI3L1; CHI3L2 were significantly increased in the AD brain [195] |
|  | for C.Neoformans in MRC1 knockout mice [194] |  |
| NAIP NLR family, apoptosis inhibitory protein | Senses bacterial flagellin [196] and type III secretion system needle proteins from several bacterial pathogens, including Salmonella typhimurium, enterohemorrhagic Escherichia coli, Shigella flexneri, and Burkholderia species [197]. Inhibits Legionella pneumophila infection [198]. | Upregulated in the AD hippocampus [29] |
| NLRP1 NLR family pyrin domain containing 1 | Activated by Bacillus anthracis lethal toxin, Toxoplasma gondii, muramyl dipeptide (a constituent of both Gram-positive and Gram-negative bacteria) [199]. NLRP1 and NLRP3 both activated by T.Gondii [200] | Monocyte expression of NLRP1, NLRP3, PYCARD, caspases 1, 5 and 8) and downstream effectors (IL-1β, IL-18) up-regulated in severe and mild AD [201] |
| NLRP3 NLR family pyrin domain containing 3 | Activated by Staphylococcus aureus , Candida albicans and | The NLRP1 and NLRP3 inflammasomes are both |
|  | the influenza virus as well as beta-amyloid [202]. Activated by C.Neoformans [203], C.Pneumoniae [204], H.Pylori [205] and by P.Gingivalis LPS [206] but also subject to proteolysis by the bacterium [207].<br>Activated and subsequently inhibited by HSV-1 [208]. Activated by the Hepatitis C virus [209] and by the Influenza A virus in dendritic cells [210]. An Epstein-Barr virus micro RNA can be secreted from infected B cells via exosomes to inhibit the NLRP3 inflammasome [211] | activated in AD monocytes [201] |
| NOD1 nucleotide binding oligomerization domain containing 1 | This protein is an intracellular pattern-recognition receptor (PRR) that initiates inflammation in response to a subset of bacteria through the detection |  |
|  | of bacterial diaminopimelic acid. Multiple alternatively spliced transcript variants differing in the 5' UTR have been described, but the full-length nature of these variants has not been determined. [provided by RefSeq, Oct 2009].<br>*P. gingivalis* outer membrane vesicles induce strong TLR2 and TLR4-specific responses and moderate responses in TLR7, TLR8, TLR9, NOD1 and NOD2 expressing-HEK-Blue cells [212]. *Nod1*(-/-) and *Nod2*(-/-) mice show delayed bacterial clearance of *C. pneumoniae* [213]. *H. pylori* activates the intracellular NOD1, NOD2, and NLRP3 [214] |  |
| NOD2 nucleotide binding oligomerization domain containing 2 | The protein is primarily expressed in the peripheral blood leukocytes. It plays a |  |
|  | role in the immune response to intracellular bacterial lipopolysaccharides (LPS) by recognizing the muramyl dipeptide (MDP) derived from them and activating the NFkB protein. Mutations in this gene have been associated with Crohn disease and Blau syndrome. Alternatively spliced transcript variants encoding distinct isoforms have been found for this gene.<br>[provided by RefSeq, Jun 2014]<br>*P. gingivalis* outer membrane vesicles induce strong TLR2 and TLR4-specific responses and moderate responses in TLR7, TLR8, TLR9, NOD1 and NOD2 expressing-HEK-Blue cells [212] |  |
| RARRES2 retinoic acid receptor responder 2: | The active protein has several roles, including that as an | Upregulated in the hippocampus [29] |
|  | adipokine and as an antimicrobial protein with activity against bacteria and fungi. [provided by RefSeq, Nov 2014]<br>Antimicrobial effects against *E. coli*, *S. aureus*, *P. aeruginosa* and *C. albicans* [215]. |  |
| RARRES3 retinoic acid receptor responder 3 | Viral RNA detector [216-220] | Upregulated in the hippocampus [29] |
| S100A4 S100 calcium binding protein A4 | Dimerises with S100A9 and stimulates AGER and TLR4 [221] | Upregulated in the hippocampus [29] |
| Calprotectin = S100A8+S100A9 | TLR4 agonist that is secreted during the stress response of phagocytes. Involved in promoting the inflammatory response to infections and a potent amplifier of inflammation [222].<br>Cytoplasmic calprotectin inhibits *C. Neoformans* growth [223]. Restricts *H. Pylori* growth [224]. Kills | Faecal levels increased in AD patients [227]. S100B, S100A9 and S100A12, but not S100A8, were consistently associated with the neuropathological hallmarks of AD in post-mortem brains [228] |
|  | Candida Spp Escherichia coli, Klebsiella spp, Staphylococcus aureus, and Staphylococcus epidermidis [225]. Confers resistance to P.Gingivalis [226]. |  |
| S100A8 S100 calcium binding protein A8 | See Calprotectin | S100B, S100A9 and S100A12, but not S100A8, were consistently associated with the neuropathological hallmarks of AD in post-mortem brains [228] |
| S100A9 S100 calcium binding protein A9 | See Calprotectin<br>S100A9 is also required for the maturation of TLR3 , which recognises viral double stranded RNA in the endosomal compartment of macrophages [229]. | Low CSF S100A9 and beta-amyloid levels in AD correlate with each other [230] |
| S100A11 S100 calcium binding protein A11 | Expression increased in the blood of infectious myocarditis patients (staphylococcal IE and streptococcal) [2,231] | Upregulated in the hippocampus [29] |
| S100A12 S100 calcium | S100A12 has antifungal | S100B, S100A9 and |
| binding protein A12 | activity against *Candida albicans*, *C. krusei*, *C. glabrata* and *C. tropicalis* and *Listeria monocytogenes* but not *Escherichia coli* K-12 or *Pseudomonas aeruginosa*[232]. Induced in response to *H. pylori* infection and inhibits bacterial growth by binding nutrient zinc [224]. | S100A12, but not S100A8, were consistently associated with the neuropathological hallmarks of AD in post-mortem brains [228]<br>Upregulated in the hippocampus [29]. |
| S100B S100 calcium binding protein B | Pathogenic bacteria increase S100B expression in human enteric glial cells where S100B integrates bacteria-induced Toll-like receptor signalling [233]. Forms complexes with TLR2 ligands, particularly fungal RNA and inhibits TLR2 via AGER (advanced glycosylation end product-specific receptor), dampening pathogen-induced inflammation. In addition, | Low serum S100B levels in AD patients [237].<br>S100B, S100A9 and S100A12, but not S100A8, were consistently associated with the neuropathological hallmarks of AD in post-mortem brains [228] |
|  | upon binding to nucleic acids, S100B activates intracellular toll receptors which feedback to inhibit S100B transcription [234].<br>Low blood levels of S100B are a marker for invasive aspergillosis [235]. S100B expression is reduced in Borna virus-infected brains and no upregulation of the expression of S100B, or RAGE, was observed in the persistently infected brains even when incited with several inflammatory stimuli, including lipopolysaccharide [236]. |  |
| TLR1 toll like receptor 1 | Recognises peptidoglycan , a component of bacterial cell walls and acylated lipoproteins as a heterodimer with TLR2[238,239].<br>cotransfection of TLR2- | Upregulated in the hippocampus [29] |
|  | TLR1 or TLR2-TLR6 required for the activation induced by *H. pylori* LPS preparations [240]. agonists of TLR1/2, TLR3, TLR4 and TLR9 increase the phagocytosis of encapsulated *Cryptococcus neoformans* [241]. *P. Gingivalis* fimbriae use TLR1 or TLR6 for cooperative TLR2-dependent activation of transfected cell lines while the bacterial lipopolysaccharide prefers TLR1 [242]. TLR1/TLR2 dimers recognise *Borrelia burgdorferi* [243]. Borna disease virus nucleoproteins and host NFKB1 share a common ankyrin-like motif. When THP1-CD14 cells were pre-treated with the viral nucleoprotein, NFKB1 activation by Toll-like receptor ligands was |  |
|  | suppressed (for TLR1/2; TLR4; TLR2/6; TLR2; TLR7/8). [244]. |  |
| TLR10 toll like receptor 10 | Involved in the response to influenza infection [245]. A TLR2/TLR10 heterodimer functions in *H. pylori* lipopolysaccharide and *Listeria monocytogenes* recognition[246,247] . | Upon Aβ stimulation, AD PBMCs generally down-regulated TLR ratios, whereas control PBMCs up-regulated TLR ratios. TLR3, TLR4, TLR5, TLR7, TLR8, TLR9, and TLR10 ratios exhibited the greatest difference between patients and control subjects [248] |
| TLR2 toll like receptor 2 | TLR2 and TLR4, acting via the adapter protein MyD88, signal responses to *Cryptococcus neoformans*, *Aspergillus fumigatus* and *Candida albicans* [249]. TLR2 and TLR4 are activated by *H. Pylori* [250]. Activated by herpes simplex (HSV-1) and *Listeria monocytogenes* in microglial cells [251,252] Activated by | TLR2 and TLR4 expression are increased in AD peripheral blood mononuclear cells [266] |
|  | *Porphyromonas gingivalis*<br>[253]. Stimulated by the hepatitis C core protein<br>[254]. TLR2 is induced by *Haemophilus influenza* (bacterium) [255] . TLR2 and TLR9 synergistically stimulate innate antiviral activities, thereby protecting against HSV infection in the brain[256]: TLR2 TLR4 and TLR9 ligands promote the microglial uptake of beta-amyloid [257] . Amyloids from bacterial curli fibrils (from *E. coli*, *Salmonella*, and some *Enterobacteriales*) activate TLR2 [258]: TLR2 recognizes many microbial components. including lipoproteins/lipopeptides from various pathogens, peptidoglycan and lipoteichoic acid from Gram-positive bacteria, |  |
|  | lipoarabinomannan from mycobacteria, glycosylphosphatidylinositol anchors from Trypanosoma cruzi, modulin from Staphylococcus epidermis, zymosan from fungi and glycolipids from Treponema maltophilum, and lipopolysaccharides preparations from Leptospira interrogans, Porphyromonas gingivalis and Helicobacter pylori [259]. HSV-1 glycoprotein B activates NF-κB activation through TLR2/TLR6 but not with TLR1 although it coimmunoprecipitates with TLR1,2 and 6[260] Activated by C.Pneumoniae which also activates TLR4 but to a lesser extent [261,262]. the production of tumor necrosis factor (TNF) $\alpha$ by |  |
|  | macrophages in response to *Toxoplasma gondii* glycosylphosphatidylinositols require the expression of both Toll-like receptors TLR2 and TLR4 [263]. Recognises HCMV [264]. Epstein-Barr virus activates TLRs, including TLR2, TLR3, and TLR9 [265]. |  |
| TLR3 toll like receptor 3 | Recognises double stranded viral RNA [267]. Antiviral against HSV-1 and upregulated by the virus in neural stem cells, resulting in beta-interferon induction [268]. TLR3 and TLR4 activate cholesterol-25-hydroxylase producing 25-hydroxycholesterol [269], which along with 27-hydroxycholesterol inhibits the replication of enveloped and non-enveloped viruses [270]. TLR3 and TLR9 | TRL3- and TLR8-expressing Monocytes/macrophages are increased in Alzheimer's disease patients and in mild cognitive impairment [272] |
|  | recognise HCMV [271] |  |
| TLR4 toll like receptor 4 | Lipopolysaccharide [273]<br>leptospiral LPS<br>Campylobacter jejuni [274]<br>Helicobacter pylori [250]<br>C.Neoformans<br>glucuronoxylomannan [275]<br>TLR2 and TLR4 activation<br>reduce Hepatitis B infection [276]<br>TLR4 896 A>G increased risk for all parasitic infections (ORG 1.59; 95%CI 1.05-2.42), malaria (1.31; 95%CI 1.04-1.66), brucellosis (2.66; 95%CI 1.66-4.27), cutaneous leishmaniasis (7.22; 95%CI 1.91-27.29), neurocysticercosis (4.39; 95%CI 2.53-7.61), Streptococcus pyogenes tonsillar disease (2.93; 95%CI 1.24-6.93) , typhoid | TLR2 and TLR4 expression are increased in AD peripheral blood mononuclear cells [266].<br>TLR4 expression increased in the Alzheimer's disease brain in regions of beta-amyloid deposition[283] |
|  | fever (2.51; 95%CI 1.18-5.34) and adult urinary tract infections (1.98; 95%CI 1.04-3.98), but was protective for leprosy (0.36; 95%CI 0.22-0.60). TLR4 1196 C>T effects were similar to TLR4 896 A>G for brucellosis, cutaneous leishmaniasis, leprosy, typhoid fever and S. pyogenes tonsillar disease, and was protective for bacterial vaginosis in pregnancy (0.55; 95%CI 0.31-0.98) and Haemophilus influenzae tonsillar disease (0.42; 95%CI 0.17-1.00). The majority of significant associations were among predominantly Asian populations and significant associations were rare among European populations.<br>Hepatitis C viral protein |  |
|  | NS5A downmodulates NKG2D on natural killer cells via the TLR4 pathway [254]. TLR2 and TLR4 activated by HSV-1 in astrocytes [277]. *P. gingivalis* GroEL protein may contribute to cardiovascular disorders by increasing TLR4 expression [278]. *P. gingivalis* outer membrane vesicles induce strong TLR2 and TLR4-specific responses and moderate responses in TLR7, TLR8, TLR9, NOD1 and NOD2 expressing-HEK-Blue cells [212]. Senses the *C. Pneumoniae* heat shock protein [279] and a bacterial phospholipase D [280].<br>Phagocytosis of *B. burgdorferi* by microglia increases expression of TLR1, -2, 4 and 5 [281]. Induced by HCMV [282] |  |
| TLR5 toll like receptor 5 | Recognises bacterial flagellin [284]. Microglia and astrocytes respond to *B. burgdorferi* through TLR1/2 and TLR5. Phagocytosis of *B. burgdorferi* by microglia increases expression of TLR1, -2, 4 and 5 [281].<br>*Toxoplasma gondii*- derived profilin triggers human TLR5-dependent cytokine production [285]. HCMV infection potentiates TLR5 ligand-stimulated cytokine production [286]. | Upregulated relative to aged controls in the AD hippocampus and superior frontal gyrus [287] |
| TLR6 toll like receptor 6 | TLR2/TLR6 dimers recognise bacterial lipoproteins (from Refseq) but are also activated in response to viral infection (Dengue virus, hepatitis C, HIV-1, influenza, inter alia ) [288-291]. HSV-1 glycoprotein B activates NF-κB activation through | NF |
|  | TLR2/TLR6 but not with TLR1 although it coimmunoprecipitates with TLR1,2 and 6[260].<br>P.Gingivalis fimbriae use TLR1 or TLR6 for cooperative TLR2-dependent activation of transfected cell lines [242]. Involved in responses to B .Burgdorferi outer surface protein A lipoprotein[292]. |  |
| TLR7 toll like receptor 7 | Senses single stranded RNA viruses in endosomes [293].<br>TLR7 and TLR8 act as endosomal recognition receptors for a number of ssRNA viruses including influenza, HIV-1, VSV, Sendai virus, coxsackie B virus, coronaviruses (mouse hepatitis virus and severe acute respiratory syndrome coronavirus), and flaviviruses (HCV, dengue virus and | Upregulated relative to aged controls in the AD superior frontal gyrus [287] |
|  | West Nile virus) [294]. P. gingivalis outer membrane vesicles induce strong TLR2 and TLR4-specific responses and moderate responses in TLR7, TLR8, TLR9, NOD1 and NOD2 expressing-HEK-Blue cells [212]. Borrelia burgdorferi induces the production of type I interferons by human dendritic cells via TLR7 and TLR9. Indoleamine 2,3-dioxygenase (IDO1) induction and kynurenine production were mediated by the same TLR7-dependent recognition process [135]. TLR7 stimulates the expression of Epstein-Barr virus latent membrane protein 1 in infected cells [295]. Epstein-Barr virus inhibits the stimulatory effect of TLR7/8 and TLR9 |  |
|  | agonists in human B lymphocytes enabling evasion of the immune system [296] |  |
| TLR8 toll like receptor 8 | An endosomal receptor that recognizes single stranded RNA viruses such as Influenza, Sendai, and Coxsackie B viruses. Also recognises bacterial RNA from streptococci [297] and Staphylococcus aureus [298]. P. gingivalis outer membrane vesicles induce strong TLR2 and TLR4-specific responses and moderate responses in TLR7, TLR8, TLR9, NOD1 and NOD2 expressing-HEK-Blue cells [212]. TLR8 is activated by Borrelia burgdorferi RNA in the phagosome of human monocytes[299] . | TRL3- and TLR8-expressing Monocytes/macrophages are increased in Alzheimer's disease patients and in mild cognitive impairment [272] |
| TLR9 toll like receptor 9 | This gene is preferentially expressed in immune cell | The rs187084 variant homozygote GG was |
|  | rich tissues, such as spleen, lymph node, bone marrow and peripheral blood leukocytes. Studies in mice and human indicate that this receptor mediates cellular response to unmethylated CpG dinucleotides in bacterial DNA to mount an innate immune response. [provided by RefSeq, Jul 2008]. TLR3, TLR7, TLR8, and TLR9 also detect distinct forms of viral nucleic acids [294]. TLR2 and TLR9 protect against HSV-1 infection in the mouse brain [256] *P. gingivalis* outer membrane vesicles induce strong TLR2 and TLR4-specific responses and moderate responses in TLR7, TLR8, TLR9, NOD1 and NOD2 expressing-HEK-Blue cells [212]. Unmethylated | significantly associated with a decreased AD risk in a Chinese study. This protective variant related to increased TLR9 expression in peripheral blood monocytes [301] . Transcription of TLR3, TLR4, TLR5, TLR7, TLR8, TLR9, and TLR10 following beta-amyloid stimulation is depressed in mononuclear cells of AD patients [248] |
| | CpG motifs in *Toxoplasma gondii* DNA induce TLR9- and IFN-β-dependent expression of DEFA5 in intestinal epithelial cells.[89].Upregulated in dendritic cells by *C.Pneumoniae* nasal infection [300] | |
| ZBP1 Z-DNA binding protein 1 | This gene encodes a Z-DNA binding protein. The encoded protein plays a role in the innate immune response by binding to foreign DNA and inducing type-I interferon production. Alternatively spliced transcript variants encoding multiple isoforms have been observed for this gene. [provided by RefSeq, Dec 2011]. ZBP1 recognises foreign DNA in the cytosol and inhibits HSV-1 replication[302].HCMV induces the interferon | ZBP1 was identified as an Alzheimer's disease susceptibility gene using hippocampal atrophy as a quantitative Trait [304] |
|  | response via ZBP1[303]. |  |

